# RIPPLET: Mutation-Only Gene and Pathway Profiling for Precision Oncology

**DOI:** 10.1101/2025.09.17.676675

**Authors:** Fabius Wiesmann, Cassandra Litchfield, Maud Mayoux, Tumor Profiler Consortium, Andreas Wicki, Burkhard Becher, Viktor Hendrik Koelzer, Holger Moch, Marija Buljan, Sonia Tugues, Bettina Sobottka

## Abstract

Clinical implementation of comprehensive genomic profiling, via whole-genome (WGS) or whole-exome sequencing (WES), is constrained by sparse mutation burdens and analytic pipelines reliant on matched transcriptomes. Currently, gene-centric analysis prevails, but overlooks the complex, multigene and pathway perturbations shaping tumor biology. We introduce RIPPLET, a DNA-only framework converting somatic variants into quantitative gene-impact scores and topology-aware pathway-perturbation profiles. By integrating tissue-specific protein–protein interaction networks with cohort-informed reweighting, RIPPLET prioritizes likely functionally relevant alterations. Applied across 33 TCGA cancer types, RIPPLET surpasses four state-of-the-art multi-omic driver-prioritization tools in recovering cancer type-specific drivers. In a cohort of metastatic cutaneous melanomas, it identifies pathway signatures that predict drug response, provide prognostic insight and distinguish immune-infiltration phenotypes without RNA data, independently validated on an in-house cohort. RIPPLET enables DNA-only inference of tumor-specific gene and pathway dysregulation, aligning with clinical sequencing workflows and offering a scalable precision-oncology strategy in transcriptome-limited settings.

## Introduction

Despite advances in molecular profiling and targeted therapies, precision oncology still faces major challenges such as treatment resistance, therapeutic failure, and variable responses. Most genome-interpretation methods are gene-centric, emphasizing recurrent, high-frequency drivers and missing the network-level interplay that shapes tumor behavior. Mutations rarely act in isolation: distinct profiles can converge on shared dysfunctions, and identical mutations yield different outcomes depending on context, as with *BRAF* in colorectal cancer versus melanoma^1^. Whole-genome sequencing (WGS) provides a comprehensive view of somatic alterations: SNVs/InDels, copy-number alterations (CNAs), and other structural variants including fusions (FUS). Unlike transcriptomics, DNA profiling is stable, reproducible, and aligned with clinical decision standards^2–6^. Yet driver discovery remains difficult due to mutation sparsity, inter-patient heterogeneity, and rare, non-recurrent events that evade frequency-based methods. The driver/passenger dichotomy is also blurring: purported passengers can cumulatively shape the microenvironment, clonal evolution, progression, and therapy response^7–10^.

These challenges argue for a systems-level perspective. Cancer arises from coordinated dysfunction of interconnected molecular modules rather than isolated hits^11^. Network-based approaches model these interactions: propagation methods diffuse mutational signal across protein–protein interaction (PPI) networks to expose impacted subnetworks^12–14^. Algorithms such as PRODIGY^15^, IMCDriver^16^, PersonaDrive^17^, and PDRWH^18^ integrate mutations with expression to nominate sample-specific drivers. However, reliance on matched RNA limits clinical utility due to variability, batch effects, and the scarcity of high-quality RNA in routine diagnostics. Methods operating directly on genomic data to produce sample-specific outputs remain comparatively underdeveloped. Further, by default, many tools depend on global, context-agnostic PPIs, risking misattributed importance. And while some combine networks with curated pathways, they typically return ranked driver gene lists rather than quantitative, sample-level pathway dysregulation. Because diverse alterations converge on a limited set of oncogenic pathways, pathway-level readouts are more reproducible and informative ^19–21^. The success of pathway analysis in transcriptomics underscores the need for an analogous, DNA-only framework.

We therefore developed RIPPLET (**R**andom-walk-based **I**nference of Gene & **P**athway **P**erturbations integrating Sample-**L**evel and Cohort-wide **E**ffec**T**s), which transforms a tumor’s somatic profile into personalized scores of gene and pathway dysregulation. RIPPLET, a DNA-only framework, combines tissue-contextualized PPIs with cohort-wide mutational patterns to place individual variants in biological context, clarifying their likely impact and uncovering shared mechanisms across tumors. Across 33 TCGA cancer types, RIPPLET outperformed four state-of-the-art methods in cancer-specific, patient-level driver identification. In metastatic cutaneous melanoma (SKCM-MET, n=372; 50 in-house^22–24^ and 322 TCGA^19^), RIPPLET produced pathway scores with predictive, prognostic, and immune-modulatory relevance. By enabling transcriptome-independent, pathway-level interpretation of tumor genomes, RIPPLET aligns with clinical sequencing workflows and supports downstream precision-oncology applications, including therapeutic prioritization, prognostic assessment, and trial stratification.

## Results

### The RIPPLET algorithm

RIPPLET is an R framework that infers patient-specific gene and pathway perturbations from DNA alone. It runs in four steps (**Fig. 1**): (1) build a filtered binary mutation matrix from WGS/WES calls (SNVs/InDels, CNAs, fusions); (2) diffuse each tumor’s mutations with Random Walk with Restart (RWR) on a tissue-specific PPI, yielding network-smoothed profiles; (3) reweight profiles by cohort similarity so similar tumors reinforce shared signals and suppress idiosyncratic noise, producing a continuous gene-impact matrix; (4) project gene scores onto canonical pathways and compute topology-aware pathway scores per sample using directed pathway structure, integrating gene magnitude and positional influence.

**Fig. 1.**
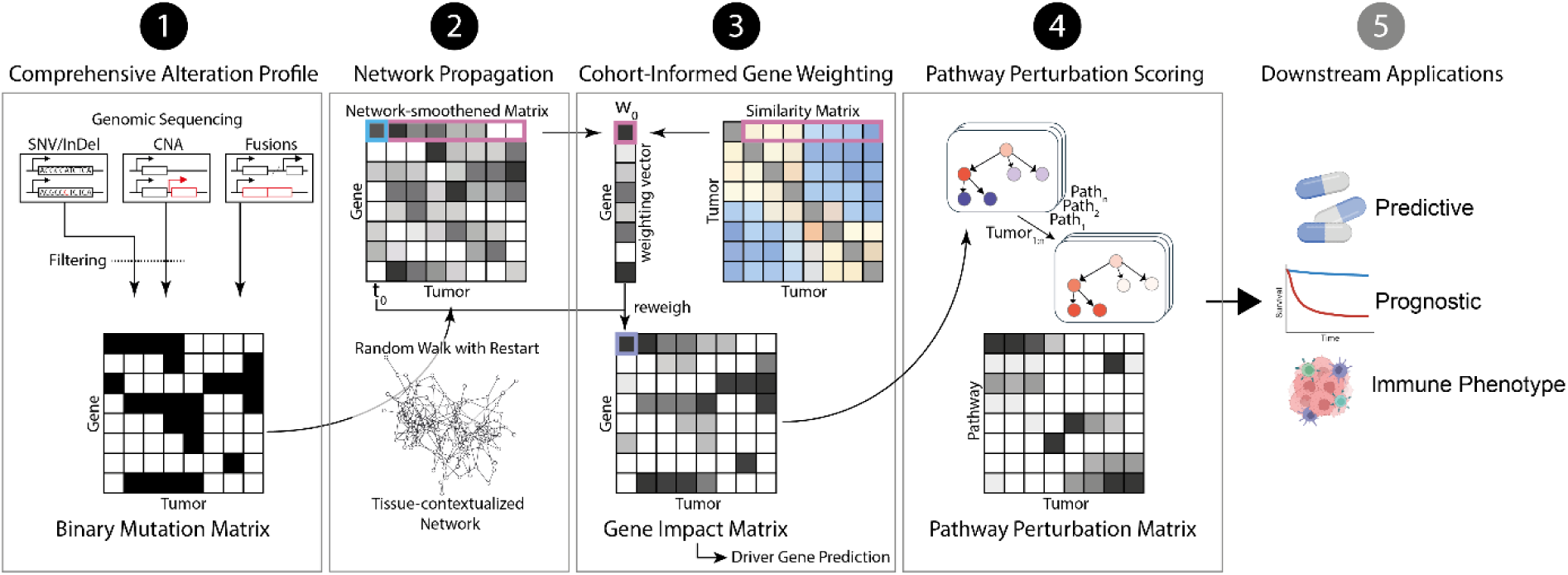
RIPPLET workflow for DNA-only, sample-specific driver gene and pathway discovery. RIPPLET transforms filtered somatic alteration calls into individualized gene- and pathway-dysregulation scores through four steps: **(1)** SNVs/InDels, CNAs and gene fusions, derived from WGS or WES, are stringently filtered and binarized into a tumor-by-gene matrix. **(2)** Each tumor’s binary profile seeds a Random Walk with Restart on a tissue-specific protein–protein interaction network, producing a smoothed mutation score for every gene that reflects both direct hits and their network neighbors. **(3)** Tumors with similar smoothed profiles inform a weighting vector for each index sample, iteratively reweighting scores to yield a continuous gene impact matrix in which each entry quantifies the likelihood of driving that tumor’s phenotype. For each patient, we rank genes in descending order to obtain personalized driver rankings. **(4)** Gene impact scores are mapped onto directed pathway graphs and aggregated, considering each gene’s position within the pathway, to produce a dysregulation score for every sample-pathway pair. Scores are aggregated and normalized to produce the pathway perturbation matrix. **(5)** Application of RIPPLET to a cohort of metastatic cutaneous melanoma samples revealed pathway dysregulation signatures that predicted therapeutic response, stratified patients by survival outcomes, and correlated with distinct tumor immune-phenotype profiles.

### Binary mutation matrix generation

We improved signal-to-noise by filtering for likely functional mutations before building the binary matrix. For SNVs/InDels, we retained only protein-altering variants that meet REVEL^25^, clinical, and allele-frequency thresholds (Methods), excluding long mutation-prone genes unless supported by significant interactome enrichment (**Supplementary Note 1**; **Supplementary Fig. 2–4, 9**). In SKCM-MET, this removed 53% of SNV/InDel calls while preserving genes linked to melanoma migration/invasion (**Supplementary Fig. 3**; **Supplementary Table 3**). For CNAs, often broad even after cohort-level consensus regions are defined^26,27^, we applied an allelic-heterogeneity filter^28^ to avoid overcalling in recurrent segments, cutting candidates from ∼55 to ∼8 per locus and eliminating 81% of amplifications and 60% of deletions (**Supplementary Fig. 5–6**). Gene fusions remain unfiltered due to the lack of standardized impact metrics and their disproportionate oncogenic potential despite rarity^29–32^. Overall, filters cut integrated mutation events by 55% and affect 33% fewer genes (**Supplementary Fig. 6**).

In SKCM-MET, mean area under the precision–recall curve (AUPRC) for frequency-based driver prioritization increased from 0.081 to 0.140 (on Known Cancer Gene reference for melanoma, KCG-MEL). Adding CNAs and fusions recovered additional biology (e.g., *PTEN*, *MTAP*, *SMYD3*, *RSF1*, *ATAD2*, *CDKN2A*, *CCND1*, *PAK1IP1*) and contributed to this performance (AUPRC = 0.133 for SNV/InDel layer data only; **Supplementary Fig. 7–8**). We observed similar gains across 33 TCGA cohorts, where mean AUPRC rose from 0.125 to 0.182 when evaluated against KCG-SPEC (cancer-type–specific drivers), with improvements in 31 cohorts (**Supplementary Table 4**). Thus, pre-filtering removed noise while preserving, and enhancing, biologically relevant signals.

### RIPPLET outperforms state-of-the-art methods in cancer type-specific driver gene identification

We benchmarked RIPPLET in SKCM-MET and across 33 TCGA cancer types by comparing its sample-level driver rankings to three curated references: KCG-SPEC (KCG-MEL for SKCM-MET), KCG-PAN (pan-cancer drivers), and KTG (immune-modulatory tumor genes). Performance was assessed using the rank-aggregation evaluation framework of Erten et al.^17^ (Methods).

In SKCM-MET, RIPPLET achieved high precision, recall, and F1 scores across all reference sets (**Supplementary** Fig. 18), and outperformed (F1 metric) four state-of-the-art methods (PRODIGY^15^, IMCDriver^16^, PersonaDrive^17^, and PDRWH^18^) in prioritizing melanoma-specific driver genes (**Fig. 2a**). Importantly, all four comparators require matched expression data, and for PersonaDrive and PRODIGY curated directional pathway maps, limiting their scalability. Against KCG-PAN and KTG, RIPPLET closely followed the top-performing alternatives (**Fig. 2b**; KTG: **Supplementary Fig. 19a**). Cohort-informed re-weighting was essential, as its removal led to a substantial decline in performance (**Supplementary Fig. 14**). Across patients, top-ranked genes were predominantly those mutated in the input matrix, while non-mutated genes, scored via network context, ranked lower (**Supplementary Fig. 15**). This pattern supports RIPPLET’s emphasis on identification of primary drivers.

**Fig. 2.**
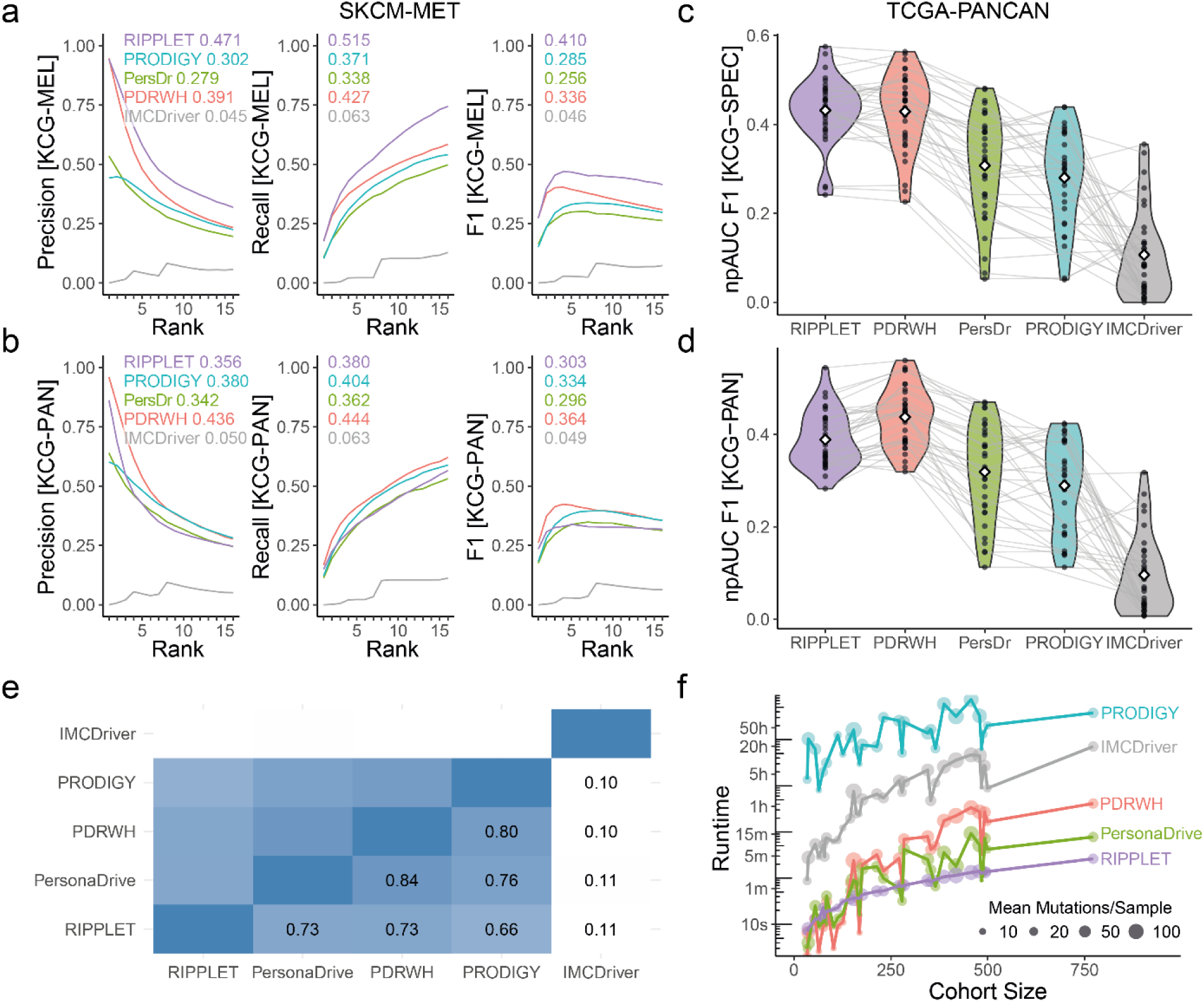
Benchmarking RIPPLET for personalized driver gene recovery. (**a**) SKCM-MET using KCG-MEL: normalized partial AUC (npAUC) of precision, recall and F1 computed to rank 16, determined using the PersonaDrive approach^17^ (Methods). Final per-sample ranks used for performance evaluation are in **Supplementary Table 5**, although RIPPLET’s gene-impact scores remain ≥ 0.01 through ∼rank 150 for SKCM-MET (**Supplementary** Fig. 16). For fairness to expression-dependent tools (PRODIGY^15^, IMCDriver^16^, PersonaDrive [PersDr]^17^, PDRWH^18^), all methods are evaluated on SKCM-MET[sub] (n = 320) with matched mutation and expression, although RIPPLET itself is DNA-only. All tools were run on identical tissue-contextualized networks. (**b**) RIPPLET npAUC on SKCM-MET[sub] against KCG-PAN. (**c–d**) Pan-cancer evaluation across 33 TCGA tumor types using (c) KCG-SPEC and (d) KCG-PAN. PRODIGY requires paired tumor–normal expression and is reported across 29 studies. (**e**) Averaged Jaccard index of predicted personal drivers (restricted to known drivers in KCG-SPEC) up to rank thresholds; RIPPLET shows high overlap with other tools in the known drivers it identified. (**f**) Algorithm runtime (log-scale) across TCGA versus cohort size, with points annotated by mean mutations per sample; unlike others, RIPPLET is insensitive to mutation burden. Full runtimes are in **Supplementary Table 6**.

Averaged across 33 TCGA cancer types, RIPPLET achieved the highest mean recovery of cancer-specific drivers (npAUC = 0.432, KCG-SPEC; **Fig. 2c**), leading in 19/33 cancers (56%; **Supplementary Table 5**) despite heterogeneous driver sets (**Supplementary Fig. 1**). On KCG-PAN/KTG it closely trailed the top tool (PDRWH; **Fig. 2d**; **Supplementary Table 5**), underscoring robust, transcriptome-independent performance across tumor types (npAUC[F1] = 0.389 vs. 0.437). Predicted drivers were highly concordant: up to each cohort-specific rank cutoff, mean pairwise Jaccard similarity on KCG-SPEC drivers showed strong overlap between RIPPLET and other top methods (**Fig. 2e**). Tissue-specific PPI contextualization improved performance in 31/33 cohorts (**Supplementary Fig. 17**) and increased SKCM-MET npAUC from 0.338 to 0.405 (+19.8%; Methods). Crucially, RIPPLET was also the most computationally efficient, completing per-sample predictions in 1min8s on average (8-core CPU, 64 GB RAM; 7s–4min23s), ∼3× faster than the next-best method and >2,500× faster than PRODIGY, enabling scalable clinical deployment (**Fig. 2f**; **Supplementary Table 6**).

Among the top five RIPPLET-ranked genes across 372 SKCM-MET tumors (maximum npAUC[F1;KCG-MEL] = 0.460, **Fig. 2b**), a total of 186 unique candidate drivers were identified. While 57 were present in the canonical cancer driver gene sets (either KCG-MEL or KCG-PAN), the remaining 129 presented potentially novel predictions in SKCM-MET.

Notably, 99/129 (76.7%, permutation P = 0.026) of our novel candidates were supported by at least one line of evidence for oncogenic relevance (**Fig. 3**): 51.9% were cited in CancerMine^33^ (hypergeometric test P = 2.24×10^-5^), 59.7% surpass the OncoScore^34^ threshold (P = 0.776), and 10.1% score as essential in CRISPR viability screens of skin-derived cell lines (P = 0.931). Of the novel predictions, 49.6% directed interacted with either KCG-MEL or KCG-PAN genes (permutation P = 0.108), indicating that they often reside in functionally similar network contexts. Clinically, 20.9% are deemed druggable in DGIdb^36^ (P = 3.0×10^-4^), while 1.6% have direct links to approved or in-development agents in the TARGET^35^ database (P = 0.190). Remarkably, 99.5% of tumors harbor at least one druggable hit among their top five RIPPLET predictions, and 94.9% still do when excluding the guideline-indicated^37^ drug targets BRAF, KIT, and MAP2K1/2 (both P < 2.2×10^-16^). Likewise, 93.8% of tumors carry a clinically approved target (72.8% after excluding those consensus targets; both P < 2.2×10^-16^). Together, these findings underscore RIPPLET’s ability to identify both diverse oncogenic drivers and actionable targets in metastatic melanoma.

**Fig. 3.**
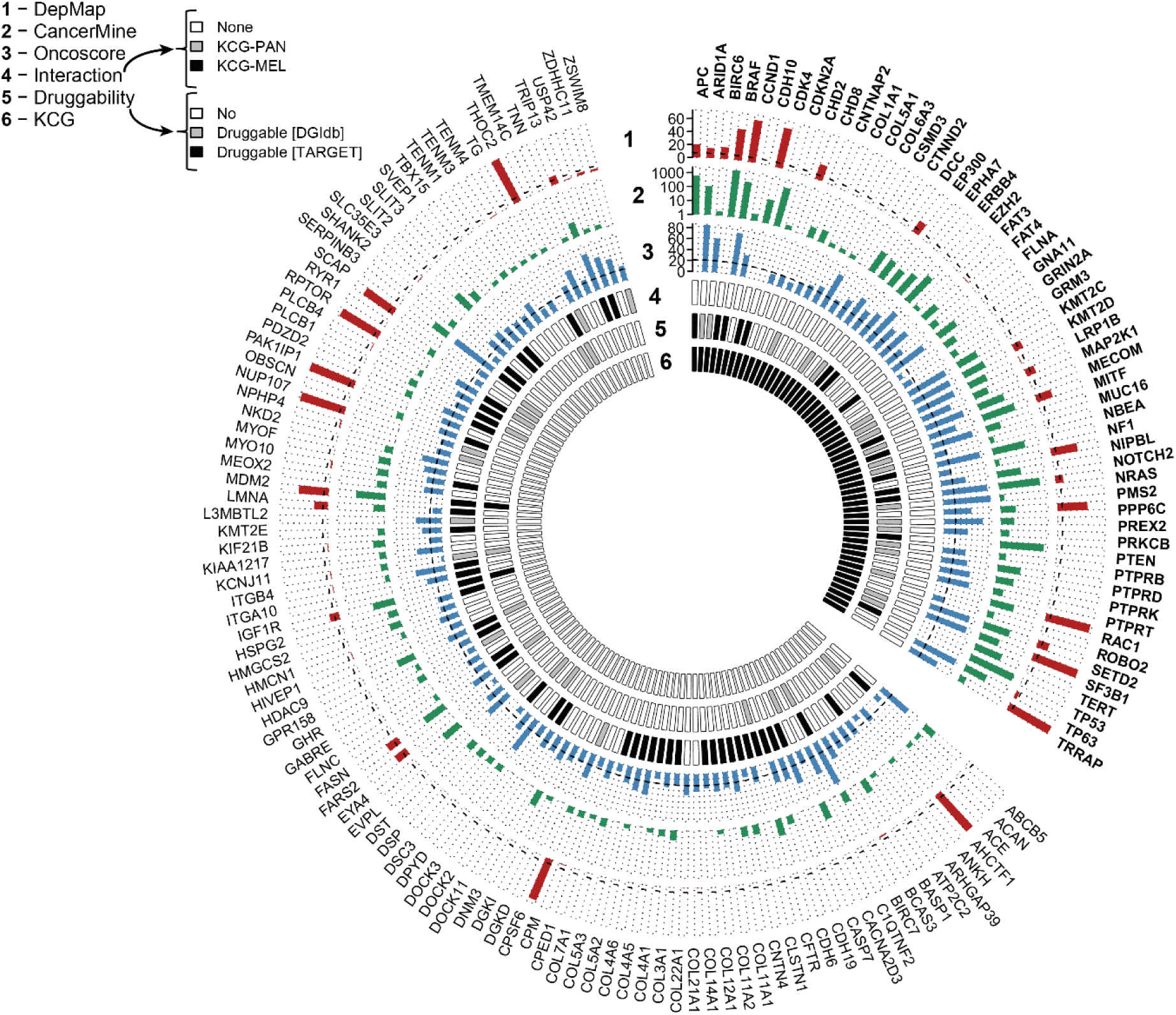
Personalized driver predictions in metastatic melanoma. Radial barplot depicting 186 unique genes prioritized in top-five sample-specific drivers by RIPPLET across 372 TCGA SKCM-MET tumors. The plot is divided into canonical drivers (n =57; bold) and novel predictions (n = 129), based on overlap with known cancer gene sets (KCG-MEL ∪ KCG-PAN). For each gene, concentric rings encode multiple lines of supporting evidence: **(1)** Gene essentiality in skin-derived cell lines from DepMap (red bars; dotted ring indicates mean count = 7.06); **(2)** CancerMine^33^ citation count (green); **(3)** OncoScore^34^ metric for oncogenic potential (blue; threshold = 21.09, dotted ring); **(4)** Protein-protein interaction with canonical drivers (white = none, grey = interacts with KCG-PAN, black = interacts with KCG-MEL); **(5)** Druggability annotation, classified as no evidence (white), DGIdb-reported (grey), or listed in the TARGET^35^ database (black); **(6)** Canonical driver status, highlighting genes previously implicated in melanoma (KCG-MEL) or pan-cancer (KCG-PAN) contexts. Together, this integrative visualization summarizes the functional and therapeutic relevance of both known and novel candidate drivers inferred from tumor-specific mutational profiles.

### Pathway analysis reveals tumor heterogeneity and improves DNA-only therapy prediction

RIPPLET’s pathway aggregation distills sparse, heterogeneous mutations into pathway profiles. In the SKCM-MET cohort, each tumor harbors on average only 1.12 high-impact (GIS ≥ 0.1) genes per perturbed pathway, yet across patients, every pathway is affected by ∼5.73 distinct genes (**Fig. 4a,b**). This pattern, intratumor mutual exclusivity (few concurrent hits per pathway) alongside interpatient heterogeneity (different genes disrupting the same pathway), dramatically enhances cohort cohesion: average inter-tumor cosine similarity rises from 0.28 at the gene level to 0.50 at the pathway level, a gain seen in all 33 TCGA cancer types and not driven by feature selection alone (**Fig. 4c, Supplementary Fig. 21**). Pathway aggregation thus reveals shared functional disruptions that remain undetectable at the single-gene level.

**Fig. 4.**
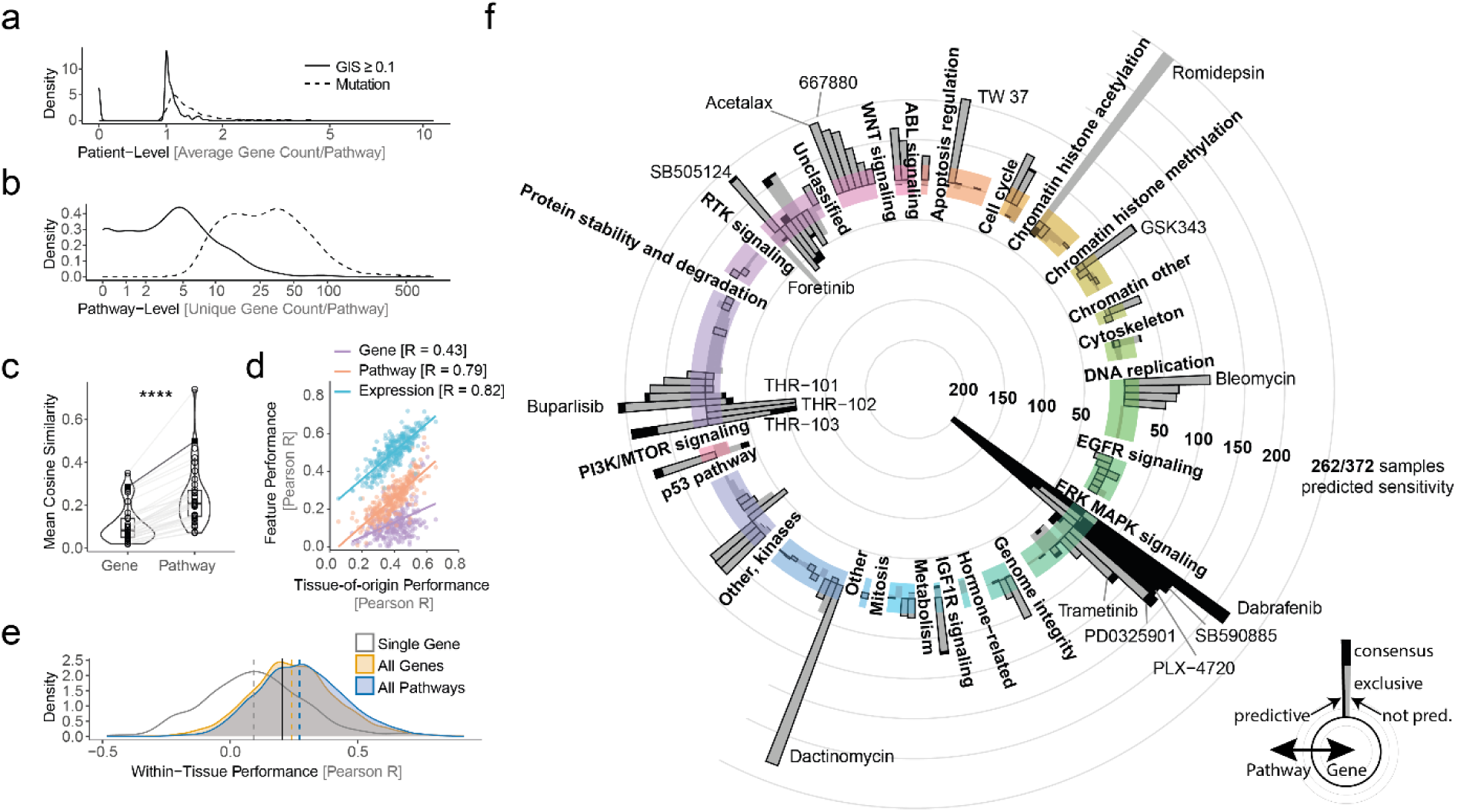
RIPPLET links heterogeneous mutations to drug response through pathway-level signatures. **(a)** Per-sample averaged pathway counts of mutated genes (dashed line) and highly impacted genes per RIPPLET (solid line, GIS ≥ 0.1), indicating mutual exclusivity. **(b)** Distribution of the mean number of uniquely mutated or impacted genes per perturbed pathway, highlighting cancer heterogeneity at the pathway level. **(c)** Violin plots of cosine similarity across PANCAN (33 cancer types) and the SKCM-MET cohort, comparing gene-(GIS) versus the pathway level. Pathway-based profiles showed significantly higher cohort cohesion with consistent gains across all cancer entities (paired Wilcoxon P < 2.2×10^-16^). The SKCM-MET cohort is indicated as a black square and line. Control groups are shown in **Supplementary** Fig. 21 supporting that this observation is not due to feature selection alone. **(d)** Per-drug predictive performance (Pearson correlation of predicted versus observed IC_50_ values) of feature layers (gene-, pathway, or expression-based predictors) versus tissue labels in tissue-agnostic elastic net (TA-EN) analysis of GDSC. Pathway-based predictors outperform gene-level models but still fall short of expression-based approaches. We verified that the observed high pathway correlation with tissue-of-origin predictive performance was not driven by tissue-specific contextualization of the gene impact scores underlying our pathway scoring (**Supplementary** Fig. 23). **(e)** Within-tissue performance of tissue-guided elastic net (TG-EN) models across 18 cancers. Densities of Pearson R for single gene (grey), all genes (orange), and all pathways (blue); dashed lines = means, solid black = predictive threshold. Pathways outperform gene-level models despite using the same underlying mutational information (mean R 0.278 vs 0.250; paired Wilcoxon P=1.1×10^-13^) and narrow the gap to expression (0.425; not shown). **(f)** Circular bar plot showing the number of SKCM-MET patients (bar length, of 372) predicted to be sensitive to each drug either by pathway (outer bars; TG-EN models) or mutation (inner bars) per drug (grouped by cellular process). For mutations, any patient with a mutation in a drug-indicated target gene is deemed sensitive. Grey segments = exclusive calls, black = consensus. Only drugs passing predictive thresholds at either mutational level (see Methods) or pathway-level (above the predictive threshold) are shown, and these are respectively highlighted by a black border around their bars. Drug labels are shown for drugs where more than 20% of the cohort were deemed sensitive.

We next evaluated whether these shared pathway perturbations could improve drug response predictions. We used RIPPLET’s inherent transfer learning ability to compute tumor-informed gene impact scores for 630 cancer cell lines from GDSC^38^ and collapsed them into pathway signatures for pan-cancer, tissue-agnostic elastic net (TA-EN) models (Methods). Pathway-based predictors achieved predictive performance for 204 of 295 drugs (mean Pearson’s R = 0.259, above the R ≥ 0.205 threshold (**Supplementary Fig. 22a**))^39^, nearly six-fold more than mutation-only models (35/295 drugs; mean R = 0.116). They recaptured ∼69% of expression-based predictive power (295/295 drugs; mean R = 0.503) and mirrored tissue-of-origin (TOO) predictive signals nearly as well (pathway R = 0.790 vs. expression R = 0.817; mutation R = 0.430) (**Fig. 4d**). However, much of the pan-cancer predictive signal was driven by capturing these TOO differences rather than mechanistic signals, as indicated by low within-tissue performance (**Supplementary Fig. 22c**). To improve this, we trained tissue-guided (TG-EN) models for 18 cancer types (≥15 cell lines each) (Methods). Therein, pathway features outperformed mutations in 12/18 tissues (mean within-tissue R = 0.278 vs 0.250, paired Wilcoxon P = 1.1×10^-13^) despite using the same underlying information and substantially closed the gap to expression-based predictors (mean R = 0.425) (**Fig. 4e**). This underscores that aggregating mutations at the pathway level enhances drug response prediction in both pan-cancer and tissue-specific contexts.

Finally, we applied the melanoma-tuned TG-EN models to predict therapies for actual metastatic melanoma tumors. We first projected the 372 SKCM-MET tumors into the same pathway space as 54 melanoma cell lines. Clustering revealed five distinct mixed groups of tumors and cell lines (C1–C5), clearly separated by consensus drivers (**Supplementary Fig. 25-26**). *BRAF*-mutant clusters (C1, C3) were predicted sensitive to MAPK inhibitors (BRAF/MEK), whereas *NRAS*-mutants or triple-wild-type (*BRAF*, *NRAS*, *NF1*)^19^ clusters (C2, C4, C5) showed diverse predicted responses, including drugs targeting PI3K/mTOR, p53, DNA-replication, and other cytotoxics (**Supplementary Fig. 26**). *NRAS*-mutant clusters (C2, C4) were predicted resistant to BRAF inhibitors but sensitive to MEK inhibitors, consistent with cell-line and clinical data^40^. Crucially, all 146 patients lacking guideline-recommended biomarkers (*BRAF*, *MAP2K1/2*, or *KIT*)^37^ were predicted to respond to at least one RIPPLET-nominated therapy (predictive pathway models only, n = 133). Further, even assuming every target mutation confers sensitivity, pathway-based predictions still yielded more patient-specific drug hits than mutation calls (mean 9.97 overall; 8.66 for pathway models meeting our predictive threshold versus 6.76 mutation-only; Wilcoxon P < 2.2×10^-16^, predictive-only P = 2.7×10^-8^; **Fig. 4d**). Yet, we observed that just 19.9% of drugs in the GDSC panel showed significant mutation-response associations and overall low predictive performance (**Fig. 4e, Supplementary Fig. 27**), underscoring the limits of single-gene biomarkers. Together, these results demonstrate that pathway-level modeling overcomes genomic heterogeneity and substantially improves DNA-based therapy prioritization in precision oncology.

### Identification of a prognostic pathway signature in metastatic melanoma

To explore whether RIPPLET-derived pathway perturbation profiles carry prognostic information in metastatic melanoma, we applied the method to the SKCM-MET[TCGA] cohort, identifying 10 Reactome pathways significantly associated with overall survival (OS) status (Wilcoxon P ≤ 0.05). From these, LASSO regression selected six protective features that all correlated with improved survival when disrupted, which were summed up into a composite Pathway Risk Score (PRS) for each patient, weighing each pathway score by its multivariate Cox coefficients (**Supplementary Table 9**). Among these, the *Toll Like Receptor 4 (TLR4) Cascade* emerged as an independent prognostic factor (HR = 1.28 × 10^-4^, P = 0.032), aligning with previous evidence of its known role in melanoma progression^41–43^.

In the SKCM-MET[TCGA] cohort, PRS-high patients exhibited significantly shorter median OS (72 days, log-rank P =0.00048) (**Fig. 5a**), and multivariate analysis confirmed the PRS as an independent prognostic factor after adjustment for tumor mutational burden (TMB; clinical threshold High=TMB ≥ 10 mut/Mb)^44,45^, age and sex (multivariate HR = 2.53; 95% CI 1.58– 4.07; P <1.20× 10^-4^) (**Fig. 5c)**, with time-dependent AUC of 0.58, 0.67, and 0.69 at 100, 200, and 300 days (**Supplementary Fig. 28**). These findings were consistent in the SKCM-MET[TUPRO] cohort where PRS-high patients had shorter OS (503 days vs. unreached, log-rank P = 0.038) (**Fig. 5b)** and comparable predictive accuracy (AUC of 0.60-0.72 at 400-600 days, **Supplementary Fig. 28**). Notably, the PRS stratification was not confounded by mutation status of common melanoma drivers: neither *BRAF* nor *NRAS* mutations were significantly associated with PRS group in either cohort (Fisher’s exact test, all P > 0.18). While TMB was inversely associated with risk group in SKCM-MET[TCGA], this was not observed in our validation cohort (**Supplementary Fig. 29**), further supporting the PRS as an independent biomarker in metastatic melanoma.

**Fig. 5.**
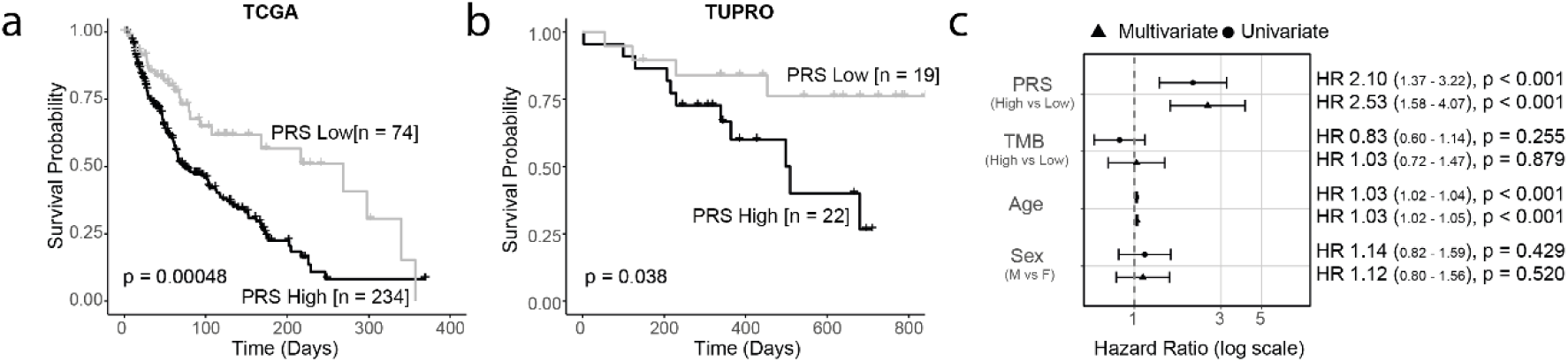
Prognostic value of the RIPPLET-derived Pathway Risk Score (PRS) in metastatic melanoma. **(a)** Kaplan-Meier curves for overall survival (OS) in the TCGA SKCM-MET cohort stratified by PRS. High risk patients (n = 234) had significantly shorter median OS (72 days) than low-risk patients (n = 74; 269 days; log-rank P = 4.8×10^-4^). **(b)** Validation in the independent TUPRO cohort where PRS high-risk patients (n = 22) also showed reduced OS compared to PRS-low (n = 19; log-rank P = 0.038), median OS was unreached in the low-risk and overall groups. **(c)** Forest plot of univariate and multivariate Cox models in SKCM-MET[TCGA] showing PRS as an independent predictor of mortality (multivariate HR = 2.53; 95% CI 1.58–4.07; P <1.20× 10^-4^) adjusted for TMB, age, and sex.

### Pathway-level predictors of tumor immune infiltration

We previously observed that RIPPLET can prioritize tumor immune microenvironment (TIME)-modifying genes (**Fig. 2a, Supplementary Fig. 19**). To comprehensively examine tumor-intrinsic regulation of immune phenotypes, we stratified the SKCM-MET cohort (n = 278; 238 SKCM-MET[TCGA], 40 SKCM-MET[TUPRO]) into inflamed, excluded and deserted subtypes based on transcriptomic^46^ and immunohistochemistry (IHC) data^47^. Inflamed tumors were associated with the most favorable overall survival, and deserted tumors with the poorest (SKCM-MET[TCGA] log-rank P < 1×10^-3^; SKCM-MET[TUPRO] P = 0.013; **Fig. 6a,b**). Although *BRAF* mutations are significantly associated with immune phenotype across all samples (Chi-sq P = 0.0107), especially non-inflamed (Fisher’s P [excluded] = 0.037, P [deserted] = 0.009), these poorly distinguished inflamed from non-inflamed tumors (Matthews correlation coefficient = 0.029). Further, consistent with the lack of significant association between tumor mutational burden (TMB) and immune subtype (Kruskal–Wallis P = 0.226; **Fig. 6c; Supplementary Fig. 30**), TMB showed near-random discriminative performance in classifying inflamed versus non-inflamed (excluded & deserted) immune phenotypes (area under the receiver-operating characteristic curve (AUROC) = 0.531 [TCGA], 0.517 [TUPRO]; **Fig. 6d**), confirming its limited utility in this setting.

**Fig. 6.**
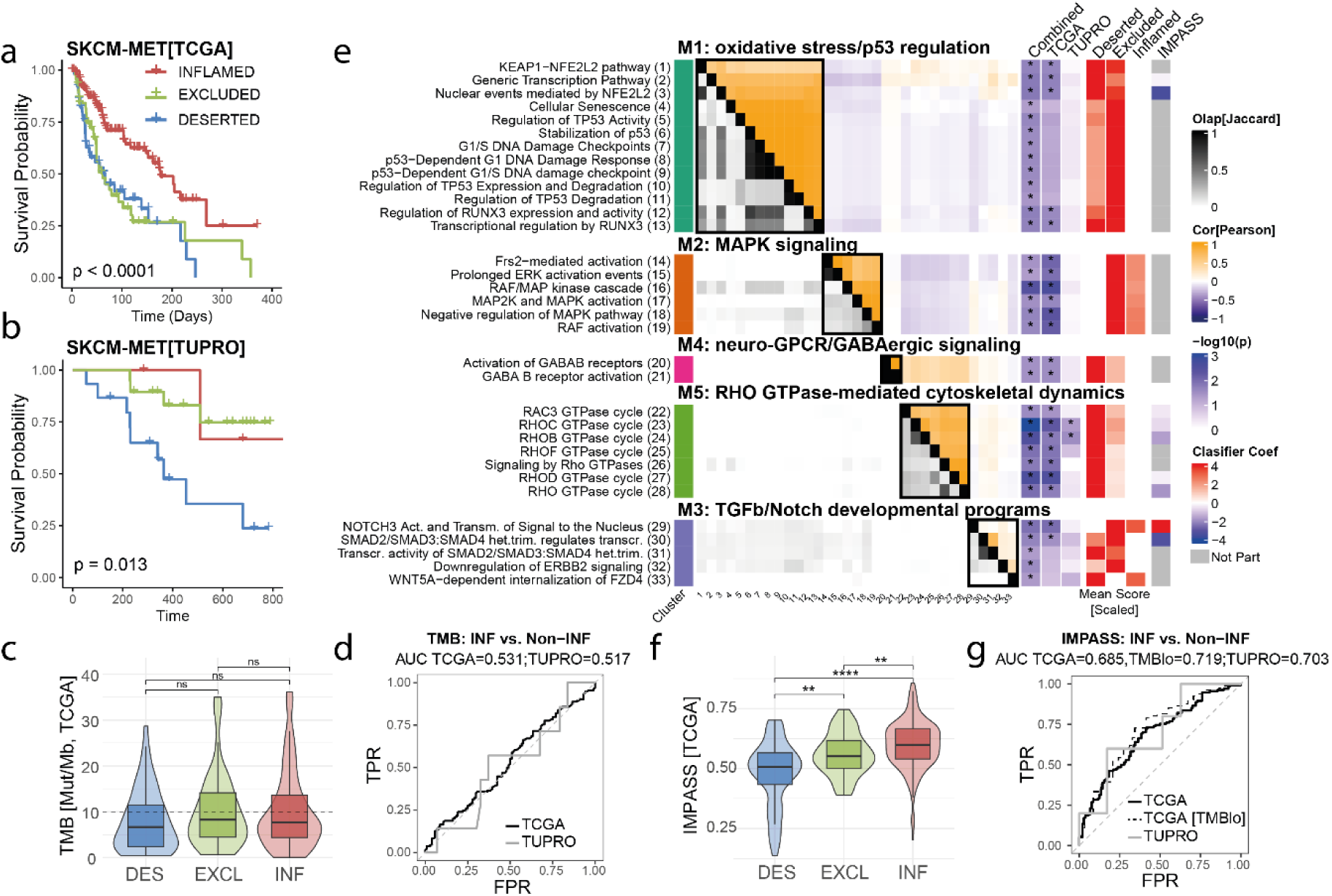
RIPPLET pathway signatures stratify immune phenotypes and reveal immune-modulatory biology. (a,b) Kaplan-Meier curves for overall survival (OS) in the (A) TCGA (n = 227 with immune & survival data) and (B) TUPRO (n = 48) cohorts, showing improved prognosis for inflamed and poorest survival for deserted tumors (log-rank P < 0.001 and P = 0.023 respectively). **(c)** Tumor mutational burden (TMB, mutations/Mb) by immune phenotype across both cohorts. No significant association was observed (Kruskal-Wallis P = 0.082). Boxes represent median ± IQR; whiskers extend to 1.5× IQR. **(d)** ROC curves assessing the ability of TMB to distinguish inflamed versus non-inflamed (excluded+deserted) tumors. Classification performance was near-random (AUROC: TCGA = 0.531; TUPRO = 0.517). **(e)** Hierarchical clustering of RIPPLET pathway scores reveals five immune-modulatory modules, including M1 = oxidative stress/p53 regulation, M2 = MAPK signaling, M3 = TGFβ/Notch signaling, M4 = GABAergic signaling, M5 = cytoskeletal dynamics. Matrix shows pathway gene overlap (Jaccard index), score correlation (Pearson), and association with immune phenotype (-log10(P), Kruskal-Wallis). Pathway perturbation patterns are scaled from 0 (white) to 1 (red). Unscaled data is shown in **Supplementary** Fig. 34. **(f)** Violin-Boxplots of immune-modulatory pathway score (IMPASS) across immune phenotypes in TCGA, with higher values associated with inflamed tumors (all P < 0.01, Wilcoxon). **(g)** ROC curves showing IMPASS performance in classifying inflamed versus non-inflamed tumors across TCGA (AUROC = 0.685), TUPRO (AUROC = 0.703), and the TMB-low subset of TCGA (AUROC = 0.719).

Using RIPPLET-derived pathway scores, we detected 33 pathways with phenotype-specific perturbation in the full SKCM-MET cohort (TCGA-only: n = 24; TUPRO-only: n = 11; Kruskal–Wallis P ≤ 0.05; **Fig. 6e**). Unsupervised clustering grouped these into five functional modules for which we derived module scores (**Methods**), four remaining significant after Benjamini-Hochberg correction (P.adj < 0.05) (**Supplementary Figure 31**). Module M2 (MAPK signaling; Kruskal-Wallis P.adj = 0.021) was notably less perturbed in deserted tumors, while module M5 (RHO GTPase signaling; Kruskal-Wallis P.adj = 0.015) scored significantly higher. Within M5, the RHOB and RHOC cycles reached nominal significance in both cohorts (P = 0.016, 0.005; BH-adjusted P > 0.45), implicating cytoskeletal remodeling next to MAPK signaling as an additional barrier to immune-cell infiltration.

To formalize the link between pathway dysregulation and immune phenotype, we developed the Immune Phenotype Assignment Signature (IMPASS) in metastatic melanoma using LASSO regression, trained on SKCM-MET[TCGA] with cross-validation for parameter selection and out-of-set performance testing, and using [TUPRO] for independent validation (**Supplementary Fig. 35**). IMPASS effectively distinguished inflamed from non-inflamed tumors across both cohorts (**Fig. 6f/g, Supplementary Figure 32**) (AUROC = 0.685 SKCM-MET[TCGA], 0.703 SKCM-MET[TUPRO]). Performance did not improve when combined with TMB (**Supplementary Figure 33**). Importantly, in tumors with low TMB (<10 mut/Mb), IMPASS alone achieved strong predictive power (AUROC SKCM-MET[TCGA] = 0.719; not enough samples in SKCM-MET[TUPRO]), identifying likely immunotherapy responders that TMB alone would miss (**Fig. 6g**)^47^. A dedicated classifier further distinguished excluded from deserted tumors (AUROC = 0.828 SKCM-MET[TCGA], 0.590 SKCM-MET [TUPRO]), again outperforming TMB (**Supplementary Figure 36**) and highlighting RIPPLET’s ability to resolve immune states beyond canonical mutation burden.

## Discussion

The present study introduces RIPPLET, a DNA-only framework that translates somatic variant calls into concise, patient-specific driver gene rankings and pathway perturbation profiles. RIPPLET’s key innovation lies in its generalizable architecture, avoiding reliance on expression or multi-omic inputs in favor of network propagation combined with cohort-informed weighting to quantify mutation impact. By drawing on recurring network perturbations across patients, which has been shown to enhance robustness and noise tolerance^17,18^, RIPPLET prioritizes gene alterations in a single, cohesive model. In practice, this end-to-end approach not only streamlines variant interpretation in settings where RNA data are unavailable or impractical, but also matches or outperforms expression-dependent methods in recovering cancer type-specific drivers, operates with dramatically reduced runtimes, and delivers consistent, cancer-agnostic gene relevance rankings across 34 diverse cohorts.

RIPPLET goes beyond individual genes by generating topology-aware pathway scores, showing that disparate mutational profiles across patients converge into common cellular process disruptions^20,21^. In this study, we leveraged these pathways alongside RIPPLET’s inherent transfer learning to align in vitro cell line models with patient tumors, markedly improving drug-response prediction from DNA data alone compared to the gene-centric models. The advantage of gene expression likely stems from expression profiles capturing genomic alterations and tissue identity while also delineating dynamic cellular states^48^. Across and within tissue types, pathway-level features consistently outperform gene-centric scores and rival expression-based signatures, while also uncovering network dependencies that single-gene or high-dimensional mutation approaches tend to miss.

Tumor mutational burden (TMB) belongs to one of the few FDA-approved immunotherapy biomarkers, yet high TMB yields inconsistent responses across, and even within, cancer types ^44,45,49–51^. In metastatic melanoma, patients meeting the clinical TMB threshold show no survival stratification. By contrast, a RIPPLET-derived composite pathway risk score robustly stratified survival in metastatic melanoma across both cohorts, supporting that pathway-level scoring may produce a more reliable biomarker. Moreover, TMB fails to predict immune infiltration in SKCM-MET, whereas RIPPLET exploits somatic disruptions in tumor-intrinsic immune-modulator pathways to infer immune phenotypes. This ability to translate genomic data into an immune landscape profile could markedly improve patient stratification and therapeutic decision-making in DNA-only settings.

Despite these strengths, RIPPLET has several limitations. Firstly, its weighting scheme is inherently cohort-driven and may underperform in small or compositionally biased datasets, risking the omission of low-frequency but clinically meaningful drivers. Secondly, whilst collapsing mutations to the gene level reduces dimensionality and data sparsity, it obscures allele-specific effects, functional variant classes, and mutational clonality. Thirdly, by binarizing alterations without respect to variant class or predicted effect, we assume equivalent phenotypic impact across diverse genomic events. Lastly, due to prioritizing a genomic-only rather than multi-omic approach, RIPPLET may fail to capture potential context- or state-dependent signaling dynamics. These may only be evident in other omic layers that are better at capturing dynamic states (e.g. transcriptomics).

In summary, RIPPLET advances the field by offering a reproducible, interpretable and computationally efficient framework that translates a patient-specific catalogue of somatic mutations into functional pathway-level consequences. Aligned with the goals of precision oncology, it enables biologically informed, patient-specific predictions of therapeutic response, prognosis, and immune phenotype, with the potential to support more tailored clinical decision-making.

## Materials & methods

### Patient cohorts & data collection

We applied RIPPLET to all 33 cancer types in The Cancer Genome Atlas (TCGA)^52^ to assess its broad utility. We further focused specifically on metastatic skin cutaneous melanoma, analyzing samples from two sources: our in-house Swiss Tumor Profiler Consortium (TUPRO) dataset (denoted SKCM-MET[TUPRO]^22–24)^ and metastatic samples of the publicly available TCGA skin cutaneous melanoma cohort (denoted SKCM-MET[TCGA]). Together, these melanoma samples are referred to as SKCM-MET.

### TCGA Data Sources

Somatic mutation calls were obtained from the MC3 (Multi-Center Mutation Calling in Multiple Cancers) dataset (MAF v0.2.8, accessed from https://gdc.cancer.gov/about-data/publications/pancanatlas). Copy number alterations were downloaded from Broad GDAC Firebrowse (http://firebrowse.org/) (results of GISTIC2 pipeline). Gene fusions were obtained from ChimerDB4.0^53^. mRNA expression data for tumor and healthy tissues were downloaded from UCSC Toil RNAseq Recompute hub (Dataset: TCGA TARGET GTEx; RSEM processed)^54^. Curated clinical data was downloaded from University of California Santa Cruz (UCSC) Xena^54,55^. We assigned MSI and hypermutation status based on the supplement of Bailey *et al*^56^. For the TCGA-SKCM cohort, immune phenotype labels were kindly provided by Mlynska *et al* upon request^46^. Study-specific sample counts with complete mutation (MC3), copy-number (GISTIC), and gene-fusion (ChimerDB) data are provided in **Supplementary Table 1**. For SKCM-MET[TCGA], we selected only metastatic samples from TCGA-SKCM (sample type code = 06, Metastatic).

### TUPRO Data Sources

Samples were collected and analyzed as recently described^24^, including immune phenotyping via IHC^47^ and somatic variant calling by whole genome sequencing (WGS)^23^.

### Gene Annotation

Ensembl (v113) Biomart was accessed via *biomaRt* R package (v2.54.1) to retrieve 19’335 protein-coding genes and associated metadata including gene length, HUGO (HGNC) symbol, and Stable Gene, Transcript and Protein Identifiers. Protein-coding genes were defined as those with at least one transcript biotype annotated as protein_coding and located on autosomes or sex chromosomes (1-22, X, Y). We defined gene length as the maximum coding sequence (CDS) length among all protein-coding transcripts per gene.

### Benchmarking and Evaluation Datasets

**Reference Driver Gene Sets** To benchmark RIPPLET’s ability to recover established cancer genes, we compiled several gold-standard reference sets. Genes for each set were identified based on COSMIC Cancer Gene Census (v101, Tiers 1–2), IntOGen (v2023.05.31), NCG canonical drivers (v7.1) and Bailey et al.^56^. Detailed filtering criteria, matching terms, and the resulting gene counts for each set are provided in **Supplementary Table 2**. This strategy resulted in two pan-cancer gene sets: pan-cancer cancer driver genes (KCG-PAN, n = 321 genes) and tumor immune microenvironment modifiers (KTG, n = 201). Additionally, we created cancer-type specific consensus gene sets for all 33 TCGA cancers (KCG-SPEC; respective gene count in **Supplementary Table 1**). For SKCM-MET, we use the same reference as for TCGA-SKCM (we refer to the melanoma-specific consensus benchmark as KCG-MEL, n = 137). We also evaluated RIPPLET’s gene-level output against literature-derived gene sets from CancerMine (https://zenodo.org/records/4925789, cancermine_collated.tsv, accessed 18.09.2024), which includes curated annotations of drivers, oncogenes and tumor suppressors.

### Driver Gene Scoring

To further prioritize and score novel candidate drivers, we used the *OncoScore* R package (v1.26.0), which assigns a cancer relevance score based on citation frequencies in biomedical literature (analysis performed on 13.11.2024)^34^.

### CRISPR-based gene essentiality

Gene essentiality data was obtained from the DepMap Portal, listing gene essentiality across 1’150 cell lines integrated using Chronos (DepMap Public 24Q2; CRISPRGeneEffect.csv, accessed 2024-04-18). Celligner^57^ data for pan-cancer comparison was also retrieved from the DepMap Portal.

### Druggability Assessment

Druggability of prioritized genes was assessed using two curated resources. First, we queried DGIdb^36^, restricting to cancer-specific interactions (anti_neoplastic = TRUE), which yielded 2’090 gene–drug relationships. Second, we consulted the TARGET database^35^ to identify genes with existing genomically guided therapies, resulting in a set of 121 actionable targets.

### Genomics of Drug Sensitivity in Cancer (GDSC)

Drug sensitivity profiles for 295 anticancer drugs across 969 cancer cell lines were downloaded from the Genomics of Drug Sensitivity in Cancer (GDSC2) portal (Release 8.5, October 2023; https://www.cancerrxgene.org/)^38^. Drug response was measured as the natural log-transformed half maximal inhibitory concentration (LN_IC50). Matched genomic information, including somatic mutation calls (v20250318), GISTIC scores from Affymetrix SNP6.0 arrays (v20191101), gene fusion predictions (v20250210), and gene expression data (RMA-normalized as per Iorio *et al*.)^39^, were retrieved via the Cell Model Passports (https://cellmodelpassports.sanger.ac.uk/) and GDSC portals (https://www.cancerrxgene.org/).

### Ethics

The study was conducted in accordance with the ethical standards of the TUPRO program (BASEC approval 2018-02050, 2018-02052, 2019-01326, 2024-01428). We also gratefully acknowledge The Cancer Genome Atlas (TCGA) Research Network for generating and sharing the genomic and clinical data used in this work. All TCGA datasets were obtained from publicly accessible repositories under the TCGA Data Use Certification and in compliance with TCGA’s publication guidelines; because these data are openly available, no additional institutional review board or ethics committee approval was required for their analysis.

### Protein-protein interaction networks

We downloaded the latest STRING^58^ protein-protein interaction network (v12.0) and retained only high-confidence interactions with a score ≥ 700 across all data channels. Protein identifiers from the Ensembl namespace (v113) were mapped to HUGO (HGNC) gene symbols. Duplicate edges, self-interactions, and redundant nodes were removed. The final high-confidence global (or context-unaware) network comprised |V| = 15’722 nodes and |E| = 225’835 unique interactions.

Given the high tissue-specificity of cancer drivers (**Supplementary Fig. 1**), the global network risks both false positives and misattributing gene importance, and tissue-specific networks were shown to be useful for prioritizing candidate disease-associated genes^59^. To generate tissue-contextualized networks for each TCGA and the SKCM-MET cancer cohorts, we employ a node removal strategy and filtered the global graph using expression data from the TOIL RNA-seq recompute dataset^59^. For each cancer entity, we retained only proteins expressed (mean transcripts per million (TPM) ≥ 1) in either cancer samples (TCGA) or corresponding normal tissues (Genotype-Tissue Expression (GTEx); if available). Known cancer type-specific cancer genes with TPM ≤ 1 were retained. Using the R *igraph* package (v1.5.1), we selected only the largest connected component to ensure steady-state convergence for network propagation. All RIPPLET networks were stored as sparse adjacency matrices A (symmetric, with Aᵢⱼ = 1 if nodes i and j are connected, 0 otherwise). Corresponding edge lists were used for benchmarking of comparator methods. Final network sizes for each cancer type are provided in **Supplementary Table 1**.

### Harmonization of genomic input data

We converted TCGA MC3 mutation calls to VCF format using maf2vcf (https://github.com/mskcc/vcf2maf) and processed all 33 tumor types (see **Supplementary Table 1** for abbreviations). Only variants labeled “PASS” in the “FILTER” column were retained for all cancer types except ovarian serous cystadenocarcinoma (OV) and acute myeloid leukemia (LAML), for which we allowed “wga”^60^. Variants were annotated as described previously^23^. Pathogenicity of missense variants was assessed with REVEL^25^, and variants with missing REVEL scores (due to Ensembl versioning differences) were assigned the highest REVEL score at the genomic position, following gnomAD’s approach (as described in gnomAD v4.1 updates, https://gnomad.broadinstitute.org/news/2024-05-gnomad-v4-1-updates/). ClinVar classifications (annotated as described previously^23^) were grouped into four categories: benign/likely benign (BLB), pathogenic/likely pathogenic (PLP), conflicting significance (both BLB and PLP terms present), and others (no BLB/PLP or conflicting status). Tumor mutational burden (TMB) for all samples was computed as the number of coding mutations (Missense_Mutation, In_Frame_Del, Nonsense_Mutation, In_Frame_Ins, Frame_Shift_Del, Frame_Shift_Ins) per genome (32.102474 Mb)^61^. To mitigate noise from hypermutated tumors, we removed hypermutated samples by applying the Tukey outlier criterion (Q3 + 1.5 × IQR) to non-silent somatic mutation counts and excluded any sample exceeding this threshold. For TCGA cases, we directly retrieved the hypermutator status from the supplementary data of Bailey et al.^56^. For SKCM-MET[TCGA], this resulted in 322 samples with complete data, of which 238 carried immune-phenotype annotations^46^. The same hypermutation filtering was applied to the SKCM-MET[TUPRO] cohort, resulting in 50 metastatic cutaneous melanoma samples with complete data including immune phenotype labels.

### Variant filtering

#### Small variants (SNV/InDel)

Somatic short variants (SNVs and InDels < 50bp) were filtered to retain variants likely to impact protein function and tumor biology.

#### Long Gene Filter

To reduce false-positive driver predictions from mutation-prone long genes, we first identified a set of very long genes using Tukey’s rule on log-transformed gene lengths and added FrequentLy mutAted GeneS (FLAGs)^62^, omitting any KCG-PAN entries from the list (**Supplementary Note 1, Supplementary Fig. 2-3**). We then assessed the functional relevance of retained long genes by comparing their mutational load within their first-degree interactors to a length-biased null model. For each sample, binarized gene alteration counts were randomly redistributed across all network genes with replacement, reassigning mutation status with a probability proportional to gene length. All SNVs and InDels in protein-coding regions (including synonymous alterations as indicators of mutational processes) were included to model the background mutation rate. For each long gene, we computed interactor mutation counts from 1’000 shuffled matrices across the cohort and retained only those with statistically elevated observed counts (Benjamini-Hochberg-adjusted P ≤ 0.05). Only mutations in these genes were retained, all others set to zero.

#### Variant Consequence Filter

We retained only non-silent somatic variants in protein-coding regions, specifically the consequences: Frame_Shift_Del, Frame_Shift_Ins, In_Frame_Del, In_Frame_Ins, Missense_Mutation, Nonsense_Mutation, Nonstop_Mutation, Splice_Site, and Translation_Start_Site.

#### Missense Pathogenicity Filter

Missense variants were scored using REVEL, a meta-predictor of pathogenicity^25^. Since REVEL gives a probability of a variant being pathogenic, we determined suitable REVEL cutoff scores using metastatic melanoma and pan-cancer ClinVar annotations. Binary ClinVar labels were assigned to missense variants classified as benign/likely benign (BLB) or pathogenic/likely pathogenic (PLP) with REVEL scores. Cumulative empirical distribution functions for BLB and PLP variants were plotted, and variants below the REVEL threshold corresponding to a 0.05 PLP rejection rate were removed (SKCM-MET = 0.276, PANCAN = 0.386) (**Supplementary Fig. 4**). Missense variants without REVEL scores were retained.

### Population Frequency & Clinical Impact Filter

Variants with a minor allele frequency >0.1% in gnomAD (v4.1; exomes preferred) or classified as BLB in ClinVar were removed.

#### Gene fusions (FUS)

For TCGA gene fusions from ChimerDB v4.0, we retained only high-confidence fusion events from the ChimerSeq-Plus pipeline^53^. Both fusion partners were considered mutated. For SKCM-MET[TUPRO], we included all “unidirectional_gene_fusion” events (annotated via Nirvana)^23^.

#### Copy number alterations (CNA)

Somatic mutation via PAM, gene fusion, or CNA can have similar effects on cancer phenotypes, i.e., allelic heterogeneity (AH)^28^. To refine driver identification from CNAs, we extended the allelic heterogeneity framework by Striker *et al*^28^ to potentially nominate multiple high-AH genes per locus. Focusing on GISTIC2 wide peaks (q ≤ 0.05), we ranked genes by their integrated final frequency (IFF): the fraction of samples with any alteration. To calculate IFF, deep deletions (DEL ≤ –2) and high-level amplifications (AMP ≥ 2) were binarized from GISTIC2 (all_thresholded.by_genes). We then merged these CNA calls with RIPPLET-filtered SNVs/InDels and fusions via bitwise OR to build an integrated mutation matrix (assigning 1 if altered in any class, 0 otherwise), and computed amplification- and deletion-specific IFFs across the cohort. Within each peak, we retained the gene with the highest IFF plus any additional outliers whose modified z-score (based on the median absolute deviation (MAD) of IFFs) exceeded the threshold. Both the minimum locus size and this MAD cutoff were optimized by hypergeometric enrichment against the KCG-SPEC driver set. In SKCM-MET and TCGA pan-cancer, we applied the filter to peaks containing >10 protein-coding genes and selected those with modified z-scores > 3.5 (**Supplementary Fig. 5**).

### Integrated mutation matrix

After filtering, we built separate binary matrices for SNVs/InDels, fusions, amplifications and deletions. We then merged them into a single genes×samples matrix, assigning a 1 whenever a gene-sample pair harbored any alteration and 0 otherwise. We kept only samples with data in all three mutation categories (SNV/InDel, CNA, fusion) and at least one mutation in the corresponding cancer type-specific network. To assess the effect of pre-filtering, we compared the number of alterations and the count of unique genes in each layer before and after RIPPLET preprocessing. To isolate mutations unique to a given mechanism, we subtracted all other layers from the integrated matrix so that any remaining hits represent that layer’s exclusive contribution. Finally, we evaluated driver-calling performance against reference gene sets (KCG-SPEC) by computing the AUPRC of gene prioritization by mutation frequency.

### Personalized gene perturbation scoring through network propagation and cohort-informed weighting strategy

#### Network propagation with sample-specific alterations

To address mutational sparsity and inter-sample heterogeneity, we used the Random Walk with Restart (RWR) algorithm to spread the influence of each mutation profile over a protein–protein interaction network^63^. For each tumor, this produced a network-smoothed gene×sample matrix, in which the state of each gene is no longer binary but continuous reflecting proximity to mutated genes^64^. We iteratively computed

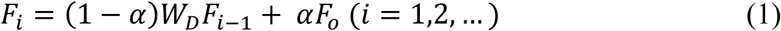

where F_0_ encodes the initial mutation status, α is the restart probability (0≤α≤1), W_D_ is the row-normalized adjacency (transition) matrix, and F_i_ denotes the distribution at iteration i^63,65^. Iteration continues updating F_i_ until convergence (|*F*_*i*_ − *F*_*i*−1_| < ε, here: ε = 1×10^-14^), at which point the stationary distribution F_S_ captures both direct and indirect mutation effects across the network. To construct *W*_*D*_, each row of the adjacency matrix A was divided by its node’s degree

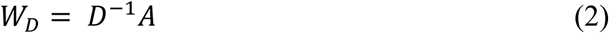

where D is the diagonal degree matrix (such that the entry d_ii_ is the sum of the i-th row of A). Row-wise normalization prevents bias toward highly connected nodes and ensures diffusion reflects input mutations rather than network topology alone (**Supplementary Note 4, Supplementary Fig. 10**)^65^. We set the default restart parameter α = 0.7 to balance local versus global diffusion (**Supplementary Fig. 11**) following prior work^63,66,67^ and observed result robustness for α in [0.9, 0.3] (**Supplementary Fig. 12**).

#### Cohort-informed reweighting via similarity in propagation space

Network-propagated scores from RWR alone neither capture shared perturbation patterns across patients nor reveal which alterations likely impact the phenotype. To enable cohort-informed gene prioritization, we apply truncated Singular Value Decomposition (SVD) to the propagated profiles, following the denoising strategy of Wang *et al.*^68^, and compute pairwise patient similarities in the resulting low-dimensional space. This approach captures functional convergence in network context, allowing similarity between tumors to be captured even in the absence of direct mutational overlap. We then pinpoint recurrently perturbed regions within these shared network contexts and use similarity-weighted frequencies to reweigh candidate drivers in individual samples. In this way, our method generalizes the classic mutation-frequency paradigm, leveraging cohort-derived similarity to highlight genes whose network-based perturbations are most likely to be functionally relevant.

To do so, each tumor-specific network-propagated vector F_S_ was normalized to sum to one (adding a small constant c=1/number of genes to avoid zeros) and log-transformed. We performed truncated SVD (20 components) via the *irlba* R package (v2.3.5.1), decomposing the matrix as:

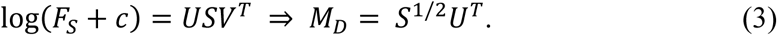

Here, M_D_ is a k×samples matrix in which each column represents a tumor in reduced-dimensional space. We compute pairwise cosine similarities between these columns and convert them to angular similarity,

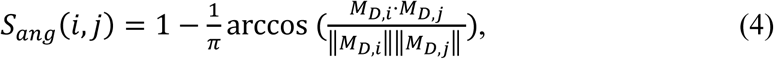

which ranges from 1 (identical) to 0 (opposite), with 0.5 orthogonal. Next, we refine each tumor’s propagated profile by borrowing information from its most similar peers. For each index tumor t_0_, we set *S*_*ang*_(*t*_0_, *t*_0_) = 0 to eliminate self-reinforcement and treat the remaining tumors as “information lenders”. Each lender *t*_*x*_ is assigned a contribution

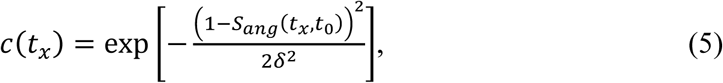

where δ controls the decay with increasing dissimilarity (default: δ=0.2, **Supplementary Fig. 13**)^18^, and normalize so ∑_*x*_ *C*_*x*_ = 1. Using the normalized contributions, we compute a cohort-derived weighting vector for the index tumor t_0_

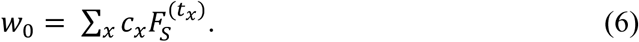

This vector encodes a gene-level weight based on cohort similarity. We then reweigh the index tumor’s own network-propagated F_S_ by multiplying it elementwise with w_0_. Finally, we linearly rescale the resulting vector to [0,1] so that the highest gene score is 1. Repeating this process for every tumor yields the final gene-by-sample impact matrix.

### Performance evaluation

#### Patient-specific benchmarking and personalized driver recovery

We benchmarked driver-prediction accuracy against patient-specific reference sets, defined as the intersection of a patient’s mutated genes (after RIPPLET preprocessing) with collected reference sets (KCG-MEL/KCG-SPEC, KCG-PAN, KTG). Mutated genes appearing in these datasets were treated as putative tumor-specific drivers^15^. Following the modified ranking-evaluation-aggregation protocol of PersonaDrive^17^, candidate genes were ranked per sample and precision, recall and F1 (**Supplementary Note 5**) were computed up to rank N, defined as twice the median number of personalized reference driver genes across the cohort after excluding samples with fewer than three patient-specific reference genes. If a tool returns fewer than N predictions for a given patient, we omit that patient from the performance calculations beyond the last available rank. To compare methods, we calculated the average partial AUC up to N of precision, recall and F1 via the trapezoidal rule and normalized it by the theoretical maximum (i.e. N) at that cutoff to obtain the normalized partial AUC (npAUC). Because N is defined by each cohort itself, npAUC provides a consistent, rank-invariant metric for comparing driver-prediction performance across different datasets. We focus on F1 as the primary performance metric (harmonic mean of the precision and recall).

#### Similarity of driver rankings

To assess concordance between driver gene–prioritization methods, we computed pairwise Jaccard similarity of their patient-specific driver sets. For each cancer type, we first determined the maximal evaluation rank defined by the KCG-SPEC reference framework^17^. For each patient and pair of methods with available predictions, we extracted the top-ranked drivers up to this cutoff, intersected them with the patient’s mutated gene set and the KCG-SPEC reference, and quantified overlap using the Jaccard index. Patient-level similarities were averaged within each cancer type and subsequently across all studies to obtain overall method concordance.

#### Comparative trial against state-of-the-art driver prediction tools

RIPPLET was benchmarked against four personalized driver prediction tools, PRODIGY^15^, IMCDriver^16^, PersonaDrive^17^ and PDRWH^18^, using identical sample selection, data inputs and PPI network references to ensure fair comparison. We restricted analyses to samples with both genomic and transcriptomic data (even though RIPPLET can score samples without expression data) and ran all methods on tissue-contextualized STRING v12.0^58^. Because matched normal samples from GTEx were unavailable for the TCGA-HNSC, -MESO, -THYM and -UVM cohorts, these cohorts were excluded from analysis with PRODIGY. For methods using additional pathway data for gene prioritization, PRODIGY was executed with default settings using Reactome pathways, while PersonaDrive employed the default kegg_pathways_v1 pathway collection. All tools were evaluated for cohort-averaged per-sample performance (npAUC of precision, recall, F1) and computational runtime.

#### Characterization of novel driver candidates in melanoma

In SKCM-MET, any gene ranked in the top driver predictions per patient (maximum F1 value) but missing from consensus drivers (KCG-MEL, KCG-PAN) was nominated as a candidate novel driver. We then classified genes as “melanoma essential” if their DepMap CRISPR gene effect score (CERES) was < -0.5 in more than the average of 7.06/73 skin-lineage cell lines^69^ and as literature-supported if they had ≥1 CancerMine^33^ citation or OncoScore^34^ ≥21.09. To test for enrichment, we performed one-tailed binomial tests comparing the number of hits among our 129 novel candidates against the background probability for each evidence set. In addition, we tested whether the 129 novel candidates were enriched for any independent line of support (DepMap essentiality, CancerMine annotation, or OncoScore ≥21.09). Enrichment was assessed against randomly sampled gene sets of equal size (100,000 permutations), with empirical P values defined as the fraction of permutations yielding at least as many supported genes. To assess translational potential, we annotated these candidates for druggability via DGIdb^36^ and flagged any with existing or emerging clinical applications in the TARGET database^35^. We used a hypergeometric model to estimate each patient’s chance of having at least one druggable or actionable gene among their top five candidates, then applied the upper tail of a binomial distribution to quantify the significance of observing this event in at least n out of all 372 patients. Finally, we mapped each candidate’s protein–protein interactions with KCG-PAN/MEL drivers to place them within the melanoma-oncogenesis network.

### Single-sample mutational pathway perturbation scoring

Somatic mutations occurring at different positions within a pathway can yield markedly different functional outcomes^70,71^. We adapted the Signaling Pathway Impact Analysis (SPIA) workflow to continuous gene impact scores derived from single-sample mutational profiles^72^. First, *graphite* (v1.52.0) was used to parse Reactome pathway data into directed graphs using the *prepareSPIA* function^73^. For each tumor, the previously generated gene impact scores were next projected onto the pathway graph to compute per-pathway perturbation scores, thereby capturing individualized pathway disruption patterns. The gene-level perturbation score (GPS) of a gene g in a pathway p is calculated as follows:

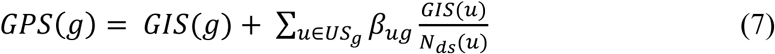

where *GIS(g)* represents the impact score of gene *g*^71,72^. The second term aggregates the impact scores of all upstream genes u that directly influence g, normalized by the number of downstream connections *N_ds_(u)*. Unlike SPIA, we define the interaction coefficient *β_ug_* as 1 when *u* and *g* directly interact in the pathway graph and 0 otherwise, disregarding directionality because we consider only positive mutational perturbations rather than signed RNA data^72^. This GPS is computed via the linear function adapted from the SPIA algorithm^71^. We apply a damping step to some pathways excluded by the original normalization constraints, preserving their outputs while enabling analysis (**Supplementary Fig. 20**). The final Pathway Perturbation Score (PPS) for each pathway *p* and sample *k* is computed by summing gene-level GPS values:

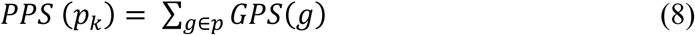

The Pathway Perturbation Score (PPS) quantifies the cumulative, weighted impact of somatic mutations on pathways. It accounts for both the overall effect of mutated genes and their positions, assigning higher scores to upstream mutations. To reduce noise from negligible gene perturbations, *GPSp,g* ≤ 0.01 were zeroed before computing PPS.

Since PPS depends on both pathway size and topology, we normalized each score by its theoretical maximum from a single-gene hit (MaxHit). For each pathway *p*, we iteratively set each gene’s GPS to 1, computed the resulting PPS, and took the maximum (*GPS_p,MaxHit_*) as the normalization constant. The normalized pathway perturbation score (NPPS) is then:

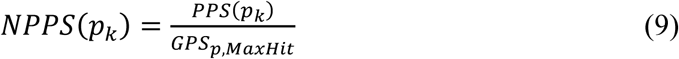

This normalization ensured that a single mutation with GIS = 1 in the most influential gene yields a normalized score of 1, with scores >1 indicating cumulative impact from multiple functionally important mutations. NPPS values thus range from 0 (no perturbation) to values potentially exceeding 1 in the presence of compounding upstream disruptions.

We focused our analysis on children of top-level pathways relevant to the given analyses from the Reactome database^74^. These include: Signal Transduction (R-HSA-162582), DNA Repair (R-HSA-73894), DNA Replication (R-HSA-69306), Circadian Clock (R-HSA-400253), Programmed Cell Death (R-HSA-5357801), Cellular responses to stimuli (R-HSA-8953897), Extracellular Matrix Organization (R-HSA-1474244), Cell-Cell Communication (R-HSA-1500931), Vesicle Mediated Transport (R-HSA-5653656), Chromatin Organization (R-HSA-48397269), Gene Expression (Transcription) (R-HSA-74160), Cell Cycle (R-HSA-1640170), Immune System (R-HSA-168256). After excluding pathways in levels 1 and 2 of Reactome hierarchy and those < 10 genes, 672 pathways were retained. For SKCM-MET-specific analyses (survival and immune associations), we further selected only pathways with (1) ≥ 70% of genes present in the tissue-contextualized network, and (2) a gene impact score of ≥ 0.2 in at least one patient, yielding a final set of 366 pathways.

### Drug response modeling and prediction

#### Pathway Level Convergence

To evaluate how diverse, patient-specific gene perturbations coalesce into common pathway-level disruptions, we first identified all pathways in the SKCM-MET cohort containing at least one gene with a Gene Impact Score (GIS) of 0.1 or higher. For each pathway, we then calculated two complementary measures: the average number of high-impact perturbations per tumor (i.e., genes with GIS ≥ 0.1 within that pathway, averaged across samples) and the average number of distinct GIS ≥ 0.1 genes per pathway across the cohort. We performed the same analyses considering all mutated genes. Next, we computed pairwise cosine similarities between samples using mutation (Mut), gene impact score (GIS) and pathway-level features in SKCM-MET and TCGA pan-cancer. For TCGA, we report the mean cosine similarity per cohort for each feature type. To rule out artifacts from differing feature counts, we repeated these comparisons with two alternative gene sets: (1) all genes annotated to our pathway collection (Mut[Path], GIS[Path]), and (2) the 672 most variably expressed genes (equal to the number of pathway features) (Mut[Var] and GIS[Var]) (**Supplementary Fig. 21**). Finally, to confirm that the convergence signal arises from pathway-specific gene composition, we generated randomized pathway definitions by permuting gene assignments within each pathway (while preserving its size and connectivity) and recalculated pairwise similarities.

#### Tumor-informed gene-impact scoring in GDSC2 cell lines

Large, well-annotated cell-line panels recapitulate patient tumor copy-number and mutational landscapes^39^. Leveraging this concordance, we analyzed 630 GDSC2 cell lines spanning 23 TCGA lineages with paired drug-response profiles (ln(IC50) for 295 compounds) and somatic mutation calls^38^ (**Supplementary Table 7**). Mutations were pre-filtered and binarized using RIPPLET’s pipeline (Methods; skipping the allelic heterogeneity filter due to limited data availability). Each line’s mutation vector was projected onto the corresponding lineage-specific protein–protein interaction network and smoothed by random walk with restart to obtain network-level gene scores. For tumor-informed transfer, we computed the similarity between each cell line’s smoothed profile and those of lineage-matched TCGA tumors to derive gene-weight vectors, which reweighted the cell-line scores to yield tumor-informed gene-impact scores. These scores served as inputs for pathway-perturbation and drug-response analyses. This design imports tumor-derived context and mitigates sparse per-entity cell-line numbers, consistent with the genomic concordance between tumors and cell lines^39^.

#### Tissue-agnostic elastic net (TA-EN) models

Our pan-cancer models leverage the fact that, despite tissue-specific initiation mechanisms and distinct genetic alterations, tumors converge on a limited set of core pathways^20,21^. Therefore, we expect that pathway-level features should transcend gene-level heterogeneity and better predict drug response. We trained pan-cancer elastic net regressions (R *glmnet* v4.1.8) to predict ln(IC_50_) for each drug using four feature sets: binary mutation calls, RIPPLET pathway scores, RNA-seq expression and one-hot–encoded tissue of origin (where each tissue type is represented by its own binary indicator column) by repeatedly splitting cell lines into 80% training and 20% test subsets. In each of two independent repetitions, we partitioned the training data into five folds for inner cross-validation: for α values from 0 to 1 in 0.1 increments and the corresponding λ sequence, we fit *cv.glmnet* and selected the α/λ pair minimizing mean squared error. We then refit an elastic net model on the full training portion using those optimal hyperparameters and predicted ln(IC_50_) in the held-out test fold. Predictive accuracy was quantified as the Pearson correlation between observed and predicted values. Averaging these correlations over all ten outer folds yielded a robust estimate of model performance for each drug-feature combination. To identify informative drug models, we pooled all genomic test-set R values, fitted a two-component Gaussian mixture (*mixtools* v2.0.0.1), and defined the predictive threshold θ at the intersection of the informative and non-informative density curves^39^.

#### Tissue-of-origin in drug response prediction

To quantify whether molecular models recapitulate tissue-driven predictive accuracy, we took the average Pearson’s R previously computed for each drug under both the tissue-only and each molecular feature model, and measured their concordance by correlating these values across all drugs^39^. Further, we evaluated *between-tissue* performance of TA-EN models following Llyod *et al*^75^ by first computing, for each drug, the mean observed and mean predicted ln(IC_50_) across all cell lines within each tissue lineage (restricting to tissues with ≥5 lines), and then calculating the Pearson correlation (R) between those tissue-level means. *Within-tissue* performance was quantified by computing, for each tissue (with ≥5 cell lines), the Pearson correlation between observed and predicted ln(IC_50_) across individual cell lines within that tissue.

#### Tissue-guided elastic net (TG-EN) models

In the clinic, oncologists often choose treatments by cancer type, but precision oncology demands finer discrimination among tumors within the same tissue^75^. To obtain models with high within-tissue performance, one approach is to train separate models per tissue, but GDSC’s limited sample sizes (mean ∼27 cell lines, range 1–62) make high-dimensional modeling (hundreds to thousands of features) prone to overfitting. To improve within-tissue accuracy without sacrificing sample size (by training only on a single tissue), we extended the tissue-guided LASSO approach of Huang et al.^76^ to elastic net (EN), which outperforms pure LASSO in drug-response modeling^48^. For each drug-tissue pair (for tissues with ≥15 CCLs, n = 18), we held out all CCLs of that tissue as a validation set and trained on the remaining CCLs from other tissues. Within each training set, we performed a grid search over the EN mixing parameter α (0–1, Δ = 0.1) and the λ path generated by *glmnet* (v4.1.8), selecting the α/λ combination that maximized Pearson’s correlation between predicted and observed ln(IC_50_) on the held-out tissue (i.e. tissue-guided). We then refit a single model on the full GDSC panel (all tissues) using those optimal hyperparameters, and report the (within-)tissue-specific R on its original validation CCLs. Thus, even though TG-EN uses gene, pathway or expression features of all CCLs in training, hyperparameters are selected in a tissue- and drug-specific manner^76^.

#### Performance assessment and mutation-only baseline

For each drug×tissue (tissues with ≥15 lines), we evaluated TG-EN models using either all genes or all pathways as predictors by computing Pearson’s R between predicted and observed ln(IC_50_) on the tissue held out for hyperparameter selection (α, λ; see TG-EN above). To place results on the same within-tissue scale, we included a single-gene baseline, reflecting the conventional molecular-pathology view, defined per drug×tissue as an “any drug-target mutated” indicator (≥1 indicated target mutated) and correlated it with ln(IC_50_) within tissue, retaining pairs with ≥5 mutant and ≥5 wild-type lines; for a binary predictor, Pearson’s R equals the point-biserial correlation. We pooled the resulting R across matched drug×tissue pairs to form the densities and compared models with paired Wilcoxon signed-rank tests. As a complementary, target-specific analysis, we tested whether mutations in a drug’s nominal target confer altered sensitivity. For this, we examined all 630 cell lines across 321 drug–target pairs in a tissue-agnostic fashion (191 compounds). Here again we retained only pairs with at least five mutant and five wild-type lines. For each remaining pair, cell lines were grouped by target-gene mutation status and compared using a Wilcoxon rank-sum test. Associations were considered mutation-dependent only if mutants showed significantly greater sensitivity (Cliff’s δ > 0 and P < 0.05).

#### Drug response prediction in SKCM-MET

We projected combined tumor- and CCL-RIPPLET pathway profiles into two dimensions via UMAP (*umap* v0.2.10.0) and identified joint clusters using k-means (*stats* v4.2.2). Within each cluster, we applied our TCGA-SKCM (melanoma) TG-EN models to predict ln(IC_50_) for 372 SKCM tumors across the 133 drugs that passed our model-performance threshold. Predicted ln(IC_50_) values were then z-score-normalized using each drug’s mean and standard deviation from the GDSC CCL panel. To binarize sensitivity, we derived drug-specific cutoffs from the CCL ln(IC_50_) waterfall curve^77^: for non-linear distributions (Pearson R ≤ 0.95), the inflection point (maximal perpendicular deviation below median) was used and converted to z-score; for linear distributions (r > 0.95), we applied a fixed threshold at ln(IC_50_) z-score ≤ –1.5, yielding 7.7% of cell-line interactions classified as sensitive. To benchmark against our pathway-based counts, we also called tumors sensitive whenever they harbored a mutation in the drug’s nominal target gene (RIPPLET-filtered). We then took all 153 drugs flagged by either approach and, for each patient, tallied the total predicted sensitivities, reporting both the overall counts (regardless of whether the corresponding pathway or gene-level model met its performance threshold) and for pathways those restricted to models that met performance thresholds.

#### Survival analysis

To ensure independence of observations in SKCM-MET[TUPRO], we retained only the earliest tumor specimen per patient, yielding 41 unique cases. All SKCM-MET[TCGA] have a unique patient identifier (n = 308 with complete survival data). Associations between RIPPLET pathway scores previously calculated (NPPS) and overall survival OS (alive vs. deceased at last follow-up) were first screen using the two-sided Wilcoxon rank-sum test. Pathways with P-values ≤ 0.05 were then subjected to L1-penalized Cox proportional-hazards modeling (LASSO, R *glmnet* v4.1.8) to guard against overfitting. Features selected by LASSO were entered into a multivariate Cox model to estimate weighted regression coefficients. For each patient, a pathway risk score (PRS) was computed as the LASSO-coefficient-weighted sum of NPPS. Optimal cutoffs were determined by maximally selected rank statistics (R *survminer* v0.4.9) on OS to dichotomize patients into PRS-high and PRS-low groups. Survival differences were assessed by Kaplan-Meier curves with log-rank tests, and model discrimination was evaluated by time-dependent AUCs at 100, 200, and 300 days (SKCM-MET[TCGA]) and 400, 500, and 600 days (SKCM-MET[TUPRO]) using *survivalROC* (v1.0.3.1) package. To assess clinical independence, we performed univariate and multivariate Cox regression, adjusting for age, sex, and TMB (dichotomized at FDA-approved cutoff ≥ 10 mut/Mb)^44,45^.

#### Immune analysis

Only tumors with complete genomic and immune-phenotype data were included (238 SKCM-MET[TCGA], 40 SKCM-MET[TUPRO]), with TUPRO cases restricted to ≤ 2 prior systemic therapies to limit treatment-related confounding. Immune phenotypes (inflamed, excluded, deserted) were derived from RNA-seq^46^ (TCGA) or pathologist-scored CD8 IHC^47^. Overall survival differences were assessed by Kaplan–Meier analysis with log-rank testing (R *survival* v3.5-7). Associations between canonical melanoma drivers (*BRAF*, *NRAS*, *KIT*, *MAP2K1*, *MAP2K2*) and immune phenotype were assessed by chi-squared tests, with per-phenotype associations by Fisher’s exact tests; discrimination of inflamed versus non-inflamed tumors (excluded + deserted) from binary mutation status was quantified using the Matthews correlation coefficient. Tumor mutational burden (TMB) associations with immune phenotype were tested by Kruskal-Wallis, with AUROC (R *pROC* v1.18.5) used for inflamed versus non-inflamed discrimination.

RIPPLET-derived per-sample pathway perturbations were compared across immune phenotypes using Kruskal–Wallis tests in SKCM-MET[TCGA], SKCM-MET[TUPRO], and the pooled cohort. Functional modules were defined by hierarchical clustering of pairwise Pearson correlations among selected pathways (R *hclust* v4.2.2, method = ward.D). Module scores were derived by first applying an inverse-normal rank transform to each pathway’s perturbation values across all samples: pathway values were ranked (average ranks for ties), converted to uniform quantiles, and then mapped to standard normal variates via the inverse Gaussian CDF. For each module, defined as a cluster of functionally related pathways, an individual sample’s module score was computed as the arithmetic mean of its transformed pathway values. Next, a LASSO-penalized logistic regression (R *glmnet* v4.1.8) was trained on SKCM-MET[TCGA] (5-fold cross-validation) to discriminate inflamed from non-inflamed tumors (excluded + deserted), yielding the IMPASS signature; a second classifier (PathClassifier2) distinguished excluded from deserted tumors. Both models were validated independently in SKCM-MET[TUPRO], with performance assessed by AUROC. To evaluate the combined predictive value of IMPASS and TMB, logistic regression meta-models (signature + TMB) were fitted separately in SKCM-MET[TCGA] and SKCM-MET[TUPRO], and AUROC determined.

## Data availability

All TCGA genomic and clinical data used in this study are publicly available (see Methods). The metastatic melanoma TUPRO cohort data are available as binary mutation matrix from the corresponding author upon request.

## Code availability

RIPPLET is implemented as an R package and available at https://github.com/wiesmannf/RIPPLET.

## Contributions

Author contributions: B.S. conceived the study. F.W. designed and implemented the methodology, performed the formal analyses, and prepared the figures and visualizations. C.L. led analysis of the SKCM-MET[TUPRO] cohort, reannotated and re-evaluated the TCGA genomic data, and harmonized datasets across cohorts. C.L. and the Tumor Profiler Consortium assembled and curated the SKCM-MET[TUPRO] dataset. C.L., M.M., A.W., B.B., V.H.K., H.M., and M.B. provided critical intellectual input and interpretation throughout. F.W. and B.S. drafted the manuscript; all authors revised the manuscript and approved the final version. S.T. and B.S. supervised the work.

## Acknowledgements

We thank Dagmar Seidl, Tara Rahimi, Susanne Dettwiler, and Fabiola Prutek for exceptional technical support. We are grateful to Agata Mlynska for providing immune-phenotype labels for SKCM-MET[TCGA]. We gratefully acknowledge the TCGA Research Network for generating and providing the data used in this study.

## Tumor Profiler Consortium

Rudolf Aebersold^5^, Melike Ak^34^, Faisal S Al-Quaddoomi^12,23^, Silvana I Albert^10^, Jonas Albinus^10^, Ilaria Alborelli^30^, Sonali Andani^9,23,32,37^, Per-Olof Attinger^14^, Marina Bacac^22^, Monica-Andreea Baciu-Drăgan^7^, Daniel Baumhoer^30^, Beatrice Beck-Schimmer^45^, Niko Beerenwinkel^7,23^, Christian Beisel^7^, Lara Bernasconi^33^, Anne Bertolini^12,23^, Bernd Bodenmiller^11,41^, Ximena Bonilla^9^, Lars Bosshard^12,23^, Byron Calgua^30^, Ruben Casanova^41^, Stéphane Chevrier^41^, Natalia Chicherova^12,23^, Ricardo Coelho^24^, Maya D’Costa^14^, Esther Danenberg^43^, Natalie R Davidson^9^, Reinhard Dummer^34^, Stefanie Engler^41^, Martin Erkens^20^, Katja Eschbach^7^, Cinzia Esposito^43^, André Fedier^24^, Pedro F Ferreira^7^, Joanna Ficek-Pascual^1,9,17,23,32^, Anja L Frei^37^, Bruno Frey^19^, Sandra Goetze^10^, Linda Grob^12,23^, Gabriele Gut^43^, Detlef Günther^8^, Pirmin Haeuptle^3^, Viola Heinzelmann-Schwarz^24,29^, Sylvia Herter^22^, Rene Holtackers^43^, Tamara Huesser^22^, Alexander Immer^9,18^, Anja Irmisch^34^, Francis Jacob^24^, Andrea Jacobs^41^, Tim M Jaeger^14^, Alva R James^9,23,32^, Philip M Jermann^30^, André Kahles^9,23,32^, Abdullah Kahraman^15,23,37^, Viktor H Koelzer^30,37,42^, Werner Kuebler^31^, Jack Kuipers^7,23^, Christian P Kunze^28^, Christian Kurzeder^27^, Kjong-Van Lehmann^2,4,9,16^, Mitchell Levesque^34^, Ulrike Lischetti^24^, Flavio C Lombardo^24^, Sebastian Lugert^14^, Gerd Maass^19^, Markus G Manz^36^, Philipp Markolin^9^, Martin Mehnert^10^, Julien Mena^5^, Julian M Metzler^35^, Nicola Miglino^36,42^, Emanuela S Milani^10^, Holger Moch^37^, Simone Muenst^30^, Riccardo Murri^44^, Charlotte KY Ng^30,40^, Stefan Nicolet^30^, Marta Nowak^37^, Monica Nunez Lopéz^24^, Patrick GA Pedrioli^6^, Lucas Pelkmans^43^, Salvatore Piscuoglio^24,30^, Michael Prummer^12,23^, Laurie Prélot^9,23,32^, Natalie Rimmer^24^, Mathilde Ritter^24^, Christian Rommel^20^, María L Rosano-González^12,23^, Gunnar Rätsch^1,6,9,23,32^, Natascha Santacroce^7^, Jacobo Sarabia del Castillo^43^, Ramona Schlenker^21^, Petra C Schwalie^20^, Severin Schwan^14^, Tobias Schär^7^, Gabriela Senti^33^, Wenguang Shao^10^, Franziska Singer^12,23^, Sujana Sivapatham^41^, Berend Snijder^5,23^, Bettina Sobottka^37^, Vipin T Sreedharan^12,23^, Stefan Stark^9,23,32^, Daniel J Stekhoven^12,23^, Tanmay Tanna^7,9^, Alexandre PA Theocharides^36^, Tinu M Thomas^9,23,32^, Markus Tolnay^30^, Vinko Tosevski^22^, Nora C Toussaint^13^, Mustafa A Tuncel^7,23^, Marina Tusup^34^, Audrey Van Drogen^10^, Marcus Vetter^26^, Tatjana Vlajnic^30^, Sandra Weber^33^, Walter P Weber^25^, Rebekka Wegmann^5^, Michael Weller^39^, Fabian Wendt^10^, Norbert Wey^37^, Andreas Wicki^36,42^, Mattheus HE Wildschut^5,36^, Bernd Wollscheid^10^, Shuqing Yu^12,23^, Johanna Ziegler^34^, Marc Zimmermann^9^, Martin Zoche^37^, Gregor Zuend^38^

^1^AI Center at ETH Zurich, Andreasstrasse 5, 8092 Zurich, Switzerland, ^2^Cancer Research Center Cologne-Essen, University Hospital Cologne, Cologne, Germany, ^3^Cantonal Hospital Baselland, Medical University Clinic, Rheinstrasse 26, 4410 Liestal, Switzerland, ^4^Center for Integrated Oncology Aachen (CIO-A), Aachen, Germany, ^5^ETH Zurich, Department of Biology, Institute of Molecular Systems Biology, Otto-Stern-Weg 3, 8093 Zurich, Switzerland, ^6^ETH Zurich, Department of Biology, Wolfgang-Pauli-Strasse 27, 8093 Zurich, Switzerland, ^7^ETH Zurich, Department of Biosystems Science and Engineering, Mattenstrasse 26, 4058 Basel, Switzerland, ^8^ETH Zurich, Department of Chemistry and Applied Biosciences, Vladimir-Prelog-Weg 1-5/10, 8093 Zurich, Switzerland, ^9^ETH Zurich, Department of Computer Science, Institute of Machine Learning, Universitätstrasse 6, 8092 Zurich, Switzerland, ^10^ETH Zurich, Department of Health Sciences and Technology, Otto-Stern-Weg 3, 8093 Zurich, Switzerland, ^11^ETH Zurich, Institute of Molecular Health Sciences, Otto-Stern-Weg 7, 8093 Zurich, Switzerland, ^12^ETH Zurich, NEXUS Personalized Health Technologies, Wagistrasse 18, 8952 Zurich, Switzerland, ^13^ETH Zurich, Swiss Data Science Center, Wasserwerkstrasse 10, 8092 Zurich, Switzerland, ^14^F. Hoffmann-La Roche Ltd, Grenzacherstrasse 124, 4070 Basel, Switzerland, ^15^FHNW, School of Life Sciences, Institute of Chemistry and Bioanalytics, Muttenz, Switzerland, ^16^Joint Research Center Computational Biomedicine, University Hospital RWTH Aachen, Aachen, Germany, ^17^Life Science Zurich Graduate School, Biomedicine PhD Program, Winterthurerstrasse 190, 8057 Zurich, Switzerland, ^18^Max Planck ETH Center for Learning Systems, ^19^Roche Diagnostics GmbH, Nonnenwald 2, 82377 Penzberg, Germany, ^20^Roche Pharmaceutical Research and Early Development, Roche Innovation Center Basel, Grenzacherstrasse 124, 4070 Basel, Switzerland, ^21^Roche Pharmaceutical Research and Early Development, Roche Innovation Center Munich, Roche Diagnostics GmbH, Nonnenwald 2, 82377 Penzberg, Germany, ^22^Roche Pharmaceutical Research and Early Development, Roche Innovation Center Zurich, Wagistrasse 10, 8952 Schlieren, Switzerland, ^23^SIB Swiss Institute of Bioinformatics, Lausanne, Switzerland, ^24^University Hospital Basel and University of Basel, Department of Biomedicine, Hebelstrasse 20, 4031 Basel, Switzerland, ^25^University Hospital Basel and University of Basel, Department of Surgery, Brustzentrum, Spitalstrasse 21, 4031 Basel, Switzerland, ^26^University Hospital Basel, Brustzentrum & Tumorzentrum, Petersgraben 4, 4031 Basel, Switzerland, ^27^University Hospital Basel, Brustzentrum, Spitalstrasse 21, 4031 Basel, Switzerland, ^28^University Hospital Basel, Department of Information- and Communication Technology, Spitalstrasse 26, 4031 Basel, Switzerland, ^29^University Hospital Basel, Gynecological Cancer Center, Spitalstrasse 21, 4031 Basel, Switzerland, ^30^University Hospital Basel, Institute of Medical Genetics and Pathology, Schönbeinstrasse 40, 4031 Basel, Switzerland, ^31^University Hospital Basel, Spitalstrasse 21/Petersgraben 4, 4031 Basel, Switzerland, ^32^University Hospital Zurich, Biomedical Informatics, Schmelzbergstrasse 26, 8006 Zurich, Switzerland, ^33^University Hospital Zurich, Clinical Trials Center, Rämistrasse 100, 8091 Zurich, Switzerland, ^34^University Hospital Zurich, Department of Dermatology, Gloriastrasse 31, 8091 Zurich, Switzerland, ^35^University Hospital Zurich, Department of Gynecology, Frauenklinikstrasse 10, 8091 Zurich, Switzerland, ^36^University Hospital Zurich, Department of Medical Oncology and Hematology, Rämistrasse 100, 8091 Zurich, Switzerland, ^37^University Hospital Zurich, Department of Pathology and Molecular Pathology, Schmelzbergstrasse 12, 8091 Zurich, Switzerland, ^38^University Hospital Zurich, Rämistrasse 100, 8091 Zurich, Switzerland, ^39^University Hospital and University of Zurich, Department of Neurology, Frauenklinikstrasse 26, 8091 Zurich, Switzerland, ^40^University of Bern, Department of BioMedical Research, Murtenstrasse 35, 3008 Bern, Switzerland, ^41^University of Zurich, Department of Quantitative Biomedicine, Winterthurerstrasse 190, 8057 Zurich, Switzerland, ^42^University of Zurich, Faculty of Medicine, Zurich, Switzerland, ^43^University of Zurich, Institute of Molecular Life Sciences, Winterthurerstrasse 190, 8057 Zurich, Switzerland, ^44^University of Zurich, Services and Support for Science IT, Winterthurerstrasse 190, 8057 Zurich, Switzerland, ^45^University of Zurich, VP Medicine, Künstlergasse 15, 8001 Zurich, Switzerland

