## Supplementary Material for "RIPPLET: Mutation-Only Gene and Pathway Profiling for Precision Oncology"

#### Cohorts and cohort-specific data

**Supplementary Table 1 Cohort composition and network parameters.** For each cohort (abbreviation: full name), we enumerate the number of TCGA tumor samples with paired mutation and expression data used to contextualize the cancer-specific network, followed by the count of matched normal samples employed in network contextualization and comparative analyses in the case of PRODIGY (which requires matched normal). We then give the size of the cohort-specific driver gene set (KCG-SPEC), and finally describe the resulting network’s dimensions: N indicates the total number of nodes, while E denotes the number of edges in each network (each unique gene–gene interaction counted once). All cohort-specific networks are extracted subsets of the global STRING v12 reference network[1]. The SKCM-MET cohort uses the same reference gene sets as the TCGA-SKCM cohort, but a unique network (only metastatic samples TCGA-SKCM for tumor data).

| Study | Study Name | n[TCGA] | n[GTEx] | n[KCG-SPEC] | NET n[N] | NET n[E] |
|---|---|---|---|---|---|---|
| **Global/Generic** | - | - | - | - | 15,722 | 225,835 |
| ACC | Adrenocortical carcinoma | 77 | 126 | 22 | 11,177 | 164,572 |
| BLCA | Bladder Urothelial Carcinoma | 407 | 9 | 109 | 11,868 | 174,602 |
| BRCA | Breast invasive carcinoma | 1,099 | 305 | 196 | 12,044 | 178,651 |
| CESC | Cervical squamous cell carcinoma and endocervical adenocarcinoma | 305 | 10 | 48 | 12,077 | 177,833 |
| CHOL | Cholangiocarcinoma | 36 | 110 | 87 | 11,649 | 174,644 |
| COAD | Colon adenocarcinoma | 289 | 307 | 95 | 11,940 | 175,482 |
| DLBC | Lymphoid Neoplasm Diffuse Large B-cell Lymphoma | 47 | 337 | 98 | 10,512 | 161,780 |
| ESCA | Esophageal carcinoma | 182 | [652] | 111 | 12,092 | 178,609 |
| GBM | Glioblastoma multiforme | 166 | 1137 | 57 | 12,189 | 178,192 |
| HNSC | Head and Neck squamous cell carcinoma | 520 | NA | 109 | 11,198 | 167,446 |
| KICH | Kidney Chromophobe | 66 | 28 | 18 | 11,457 | 165,442 |
| KIRC | Kidney renal clear cell carcinoma | 531 | 28 | 62 | 11,808 | 173,901 |
| KIRP | Kidney renal papillary cell carcinoma | 289 | 28 | 32 | 11,653 | 170,168 |
| LAML | Acute Myeloid Leukemia | 173 | 337 | 155 | 10,570 | 160,268 |
| LGG | Brain Lower Grade Glioma | 523 | 1137 | 54 | 11,947 | 173,048 |
| LIHC | Liver hepatocellular carcinoma | 371 | 110 | 125 | 10,835 | 163,021 |
| LUAD | Lung adenocarcinoma | 515 | 288 | 138 | 12,218 | 181,508 |
| LUSC | Lung squamous cell carcinoma | 498 | 288 | 79 | 12,373 | 182,718 |
| MESO | Mesothelioma | 87 | NA | 35 | 11,235 | 167,405 |
| OV | Ovarian serous cystadenocarcinoma | 427 | 88 | 68 | 12,003 | 176,877 |
| PAAD | Pancreatic adenocarcinoma | 179 | 167 | 92 | 11,988 | 177,100 |
| PCPG | Pheochromocytoma and Paraganglioma | 182 | 126 | 8 | 11,634 | 168,549 |
| PRAD | Prostate adenocarcinoma | 495 | 100 | 134 | 11,996 | 175,961 |
| READ | Rectum adenocarcinoma | 93 | 307 | 99 | 11,948 | 175,787 |
| SARC | Sarcoma | 262 | 395 | 63 | 11,507 | 172,049 |
| SKCM | Skin Cutaneous Melanoma | 469 | 555 | 137 | 12,031 | 177,852 |
| **SKCM-MET** | Metastatic Skin Cutaneous Melanoma | 320 | 555 | 137 | 12,057 | 178,196 |
| STAD | Stomach adenocarcinoma | 413 | 174 | 133 | 11,886 | 176,479 |
| TGCT | Testicular Germ Cell Tumors | 154 | 165 | 11 | 13,368 | 195,937 |
| THCA | Thyroid carcinoma | 512 | 279 | 80 | 11,762 | 170,605 |
| THYM | Thymoma | 119 | NA | 24 | 11,153 | 166,720 |
| UCEC | Uterine Corpus Endometrial Carcinoma | 181 | 78 | 95 | 12,015 | 177,385 |
| UCS | Uterine Carcinosarcoma | 57 | 78 | 29 | 12,125 | 176,927 |
| UVM | Uveal Melanoma | 79 | NA | 18 | 9,832 | 145,218 |

**Supplementary Table 2 Matching terms across data sources.** For each cohort (abbreviation), this table lists the strings used to match entries across GTEx, NCG, COSMIC, IntOGen, Bailey et al.[2], and GDSC[3]. The first two rows show terms for the pan-cancer driver (KCG-PAN) and immune-modulating gene (KTG) reference sets. CGC entries were limited to somatic alterations (Somatic = yes); NCG entries to coding genes (coding_status = "coding").

| Study | GTEx | NCG | COSMIC | IntOGen | Bailey | GDSC |
|---|---|---|---|---|---|---|
| KCG-PAN | - | canonical_drivers_Pan-cancer adult | - | - | PANCAN | - |
| KTG | - | TIME_canonical_drivers | - | - | - | - |
| ACC | Adrenal Gland | adrenocortical_carcinoma | adrenocortical; adrenocortical carcinoma; adrenal adenoma; adrenal aldosterone producing adenoma; adrenal carcinoma; adrenal aldosterone producing adenoma | ACC | ACC | ACC |
| BLCA | Bladder | bladder_cancer | bladder cancer; urothelial carcinoma; bladder carcinoma | BLCA | BLCA | BLCA |
| BRCA | Breast - Mammary Tissue | breast_cancer;male_breast_cancer; triple_negative_breast_cancer | breast;cancer; lobular breast; breast carcinoma; phyllodes tumour of the breast; secretory breast | BRCA | BRCA | BRCA |
| CESC | Cervix - Ectocervix; Cervix - Endocervix | cervical_cancer_(all_histologies) | cervical carcinoma | CESC | CESC | CESC |
| CHOL | Liver | cholangiocarcinoma; hepatocellular_carcinoma, cholangiocarcinoma | cholangiocarcinoma; biliary tract; biliary tract carcinoma | CHOL | CHOL | - |
| COAD | Colon - Sigmoid; Colon - Transverse | colorectal_adenocarcinoma | colon cancer; colorectal carcinoma; colorectal cancer; colorectal adenocarcinoma; colon carcinoma; colon adenocarcinoma; large intestine carcinoma; CRC | COAD | COADREAD | COREAD |
| DLBC | Whole Blood | diffuse_large_B-cell_lymphoma; dlblc, follicular_lymphoma | DLBCL; diffuse large B-cell lymphoma; ABC-DLBCL | DLBCLNOS | DLBC | DLBC |
| ESCA | Esophagus - Gastroesophageal Junction; Esophagus - Mucosa;Esophagus - Muscularis | esophageal_(squamous_and_adenocarcinoma); esophageal_adenocarcinoma; esophageal_squamous_carcinoma | oesophagus; oesophageal SCC; oesophagus cancer; oesophageal squamous cell carcinoma | ESCA | ESCA | ESCA |
| GBM | Brain - Amygdala; Brain - Anterior cingulate cortex (BA24); Brain - Caudate (basal ganglia); Brain - Cerebellar Hemisphere; Brain - Cerebellum;Brain - Cortex; Brain - Frontal Cortex (BA9); Brain - Hippocampus; Brain - Hypothalamus; Brain - Nucleus accumbens (basal ganglia); Brain - Putamen (basal ganglia); Brain - Spinal cord (cervical c-1); Brain - Substantia nigra | glioblastoma; paediatric_high-grade_glioma | GBM;glioblastoma; paediatric glioblastoma | GMB; HGGNOS | GBM | GBM |
| HNSC | - | oral_squamous_cell_carcinoma; squamous_head_and_neck_cancer | HNSCC; head and neck; head and neck cancer;oral squamous cell; head and neck SCC; oral SCC | HNSC | HNSC | HNSC |
| KICH | Kidney - Cortex | chromophobe_renal_cell_carcinoma | chromophobe renal carcinoma; renal | CHRCC | KICH | - |
| KIRC | Kidney - Cortex | clear_cell_renal_cancer | CCRCC;clear cell renal carcinoma;renal cell carcinoma;clear cell renal cell carcinoma | CCRCC; RCC | KIRC | KIRC |
| KIRP | Kidney - Cortex | papillary_renal_cell_carcinoma | papillary renal carcinoma;renal | PRCC | KIRP | - |
| LAML | Whole Blood | acute_myeloid_leukemia (AML); acute_lymphoblastic_leukemia; acute_lymphocytic_leukemia | AML; acute megakaryocytic leukaemia; de novo AML; MDS/MPN-U; myelodysplastic syndrome; CMML; AML(CMLblasttransformation); sAML | AML | LAML | LAML |
| LGG | See GMB | astrocytoma; low_grade_glioma | low-grade glioma; oligodendroglioma; pilocytic astrocytoma; other CNS; astrocytoma | LGGNOS | LGG | LGG |
| LIHC | Liver | hepatocellular_carcinoma | hepatocellular carcinoma; hepatocellular; fibrolamellar hepatocellular carcinoma; liver;hepatic adenoma | HCC | LIHC | LIHC |
| LUAD | Lung | lung_adenocarcinoma | lung adenocarcinoma; NSCLC;lung cancer; lung carcinoma; lung | LUAD | LUAD | LUAD |
| LUSC | Lung | lung_squamous_cell_carcinoma | lung SCC; squamous cell carcinoma; Lung SCC | LUSC | LUSC | LUSC |
| MESO | - | malignant pleural mesothelioma | mesothelioma; malignant mesothelioma | PLMESO | MESO | MESO |
| OV | Ovary | ovarian_cancer; ovarian_serous_carcinoma | ovarian; serous ovarian;clear cell ovarian carcinoma; epithelial ovarian; borderline ovarian; low grade serous ovarian cancer; ovarian mixed germ cell tumour; granulosa-cell tumour of the ovary | OVT | OV | OV |

| | | | | | | |
|---|---|---|---|---|---|---|
| PAAD | Pancreas | pancreatic_cancer_(all_histologies); pancreatic_ductal_adenocarcinoma | pancreatic cancer; pancreatic carcinoma; pancreatic; pancreas acinar carcinoma; pancreatic ductal adenocarcinoma; pancreatic neuroendocrine tumour; pancreatic intraductal papillary mucinous neoplasm | PAAD; PANCREAS | PAAD | PAAD |
| PCPG | Adrenal Gland | pheochromocytoma,_paraganglioma | pheochromocytoma; paraganglioma | | PCPG | - |
| PRAD | Prostate | prostate_cancer | prostate; prostate carcinoma; prostate cancer; prostae adenocarcinoma | PRAD | PRAD | PRAD |
| READ | Colon - Sigmoid;Colon - Transverse | colorectal_adenocarcinoma | rectal adenocarcinoma; colorectal carcinoma; CRC | COADREAD | COADREAD | COREAD |
| SARC | Muscle - Skeletal | angiosarcoma; leiomyoma; leiomysarcoma, liposarcoma; soft_tissue_sarcoma | sarcoma; soft tissue sarcoma; liposarcoma; osteosarcoma; extraskeletal myxoid chondrosarcoma; synovial sarcoma; clear cell sarcoma; angiosarcoma; leiomyoma | LMS; SOFT_TISSUE | SARC | - |
| SKCM | Skin - Not Sun Exposed (Suprapubic); Skin - Sun Exposed (Lower leg) | melanoma; desmoplastic_melanoma | melanoma; cutaneous melanoma | MEL | SKCM | SKCM |
| STAD | Stomach | gastric_adenocarcinoma; gastric_cancer; mucinous_gastric_cancer; pan-gastric | stomach carcinoma; gastric carcinoma; gastric adenocarcinoma;gastric; diffuse gastric | STAD | STAD | STAD |
| TGCT | Testis | testicular_germ_cell_cancer | testicular germ cell tumour;sex cord-stromal tumour | MGCT | TGCT | - |
| THCA | Thyroid | papillary_thyroid_cancer; anaplastic_thyroid_carcinoma; hurthle_cell_carcinoma | papillary thyroid; thyroid cancer; follicular thyroid; anaplastic thyroid cancer; thyroid cancer(PDTCandATC) | WDTC | THCA | THCA |
| THYM | - | thymic_carcinoma | thymoma | THYM | THYM | - |
| UCEC | Uterus | endometrial_cancer; clear_cell_endometrial_cancer; serous_endometrial_cancer; uterine_carcinosarcoma | endometrial cancer; endometrial carcinoma; endometrioid carcinoma; endometrioid adenocarcinoma | UCEC | UCEC | UCEC |
| UCS | Uterus | uterine_carcinosarcoma | uterine carcinosarcoma; uterine leiomyoma; uterine serous carcinoma; endometrial stromal tumour | UCS | UCS | - |
| UVM | - | uveal_melanoma | uveal melanoma | UM | UVM | - |

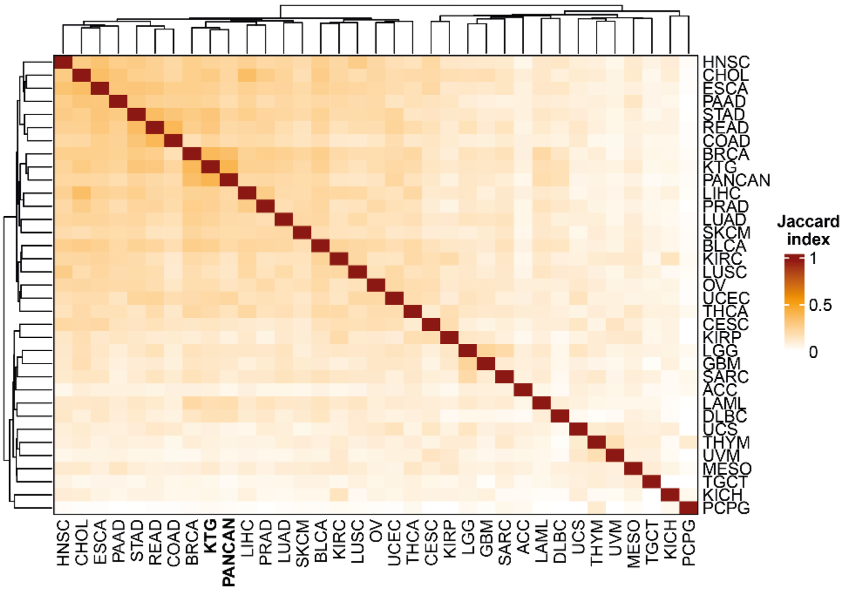


**Supplementary Fig. 1 Tissue specificity of driver mutations.** Although all cells share the same genome, tissue-specific differences in gene expression and protein function reshape network topology and pathway structures, resulting in distinct driver gene sets across 33 TCGA cancer types (KCG-SPEC) and the pan-cancer references (KCG-PAN, KTG; bold). Shown are pairwise Jaccard similarity indices among these sets (mean = 0.146 (excluding self-comparisons, KCG-SPEC only). Despite this heterogeneity, RIPPLET achieved the highest F1 score in 19 of 33 cancer types (56%). SKCM-MET uses the same cancer type specific driver genes as TCGA-SKCM.

### Binary mutation matrix

#### Supplementary Note 1: Long gene filter

Long genes accrue more mutations simply by length, skewing both gene-level and network analyses toward false positives. Long genes harbor more alterations, including many variants of unknown significance (VUS), so false positives are likely to persist despite passenger filtering, inflating mutation frequency. Because cancer drivers are significantly longer than non-drivers (mean lengths: non-drivers 1,733 bp versus KCG-PAN 3,134 bp; KCG-SPEC (all cancer types combined) 3,551 bp; KCG-MEL 4,170 bp; all Wilcoxon $P < 2.2\times10^{-16}$), excluding long genes risks discarding true positives, that have previously also shown to be highly interconnected with drivers and potentially relevant[4]. To address this, we identified a set of very long genes prone to random mutations and perform gene-level filtering. Specifically, we log-transformed lengths of all protein-coding genes (fit to a log-normal distribution: $\mu = 3.11$, $\sigma = 0.32$) and applied Tukey's rule ($Q_3 + 1.5$ IQR) to flag 195 outliers. After removing consensus drivers (KCG-PAN), 182 very long genes remained, exhibiting inflated mutation rates in SKCM-MET (14.8% vs. 2.2%; Wilcoxon $P < 2.2\times10^{-16}$). From Shyr et al.[5], we incorporated 83 FLAG genes, also enriched for mutations (19.9 % vs. 2.3 %; $P < 2.2\times10^{-16}$), of which 78 overlapped our long-gene set ($n_{final}$ = 197).

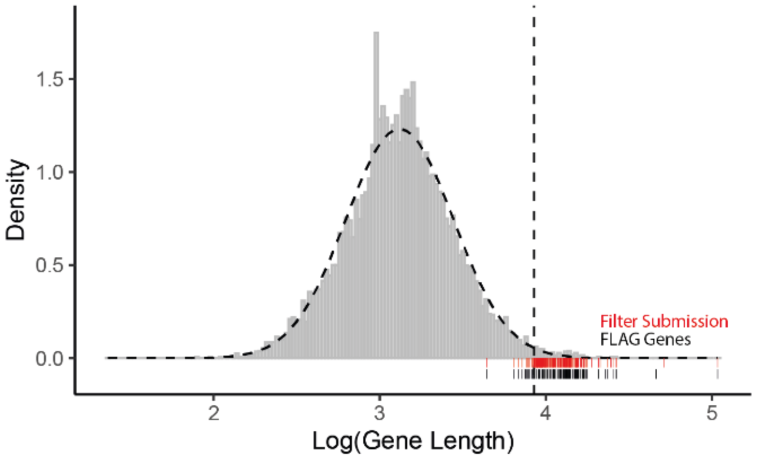


**Supplementary Fig. 2 Log-transformed length distribution of human protein-coding genes follows a log-normal model** ($\mu$ = 3.11, $\sigma$ = 0.32). The vertical line at log-length = 3.929 (8,482 bp) marks the Tukey outlier threshold. Black ticks show FLAG genes; red ticks denote all non-KCG-PAN FLAG and "Very Long" genes subjected to filtering.

To assess functional relevance, we compared each very-long gene's interactome mutation burden to a degree-preserving null model by summing mutations across first-degree interactors and generating null distributions via permutation. 26 genes showed significant interactome enrichment (BH-adjusted P ≤ 0.05), distinguishing them from "star-like" patterns of passenger bias in which only the index gene is mutated without supporting mutations in its PPI network neighborhood[6,7] (**Supplementary Fig. 3**).

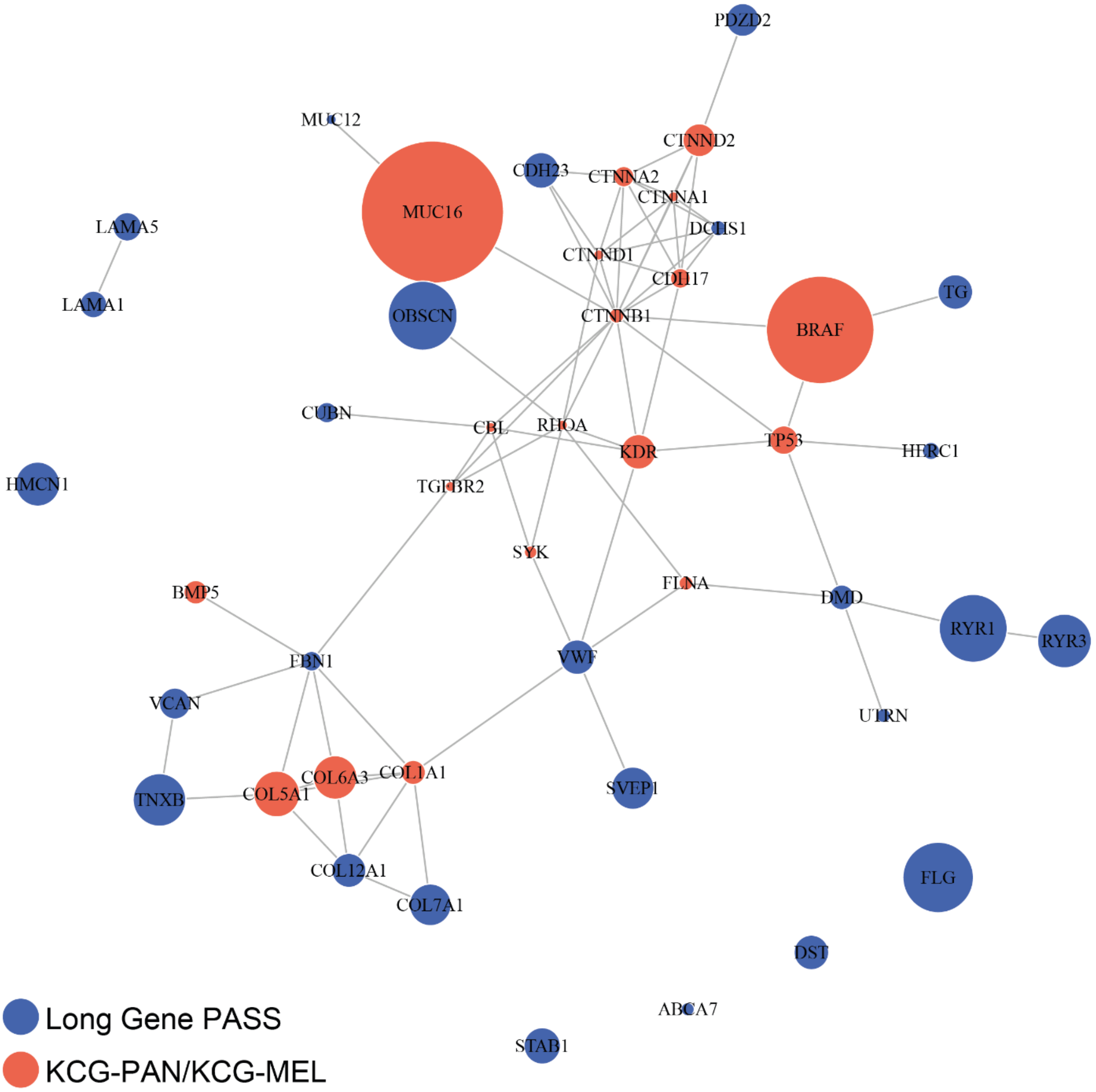


**Supplementary Fig. 3 Permutation-based long-gene filter highlights direct interactors of established cancer drivers.** Shown in red are the KCG-MEL and KCG-PAN proteins that directly (first order) interact with long genes (blue) passing the mutated-interactome permutation test in SKCM-MET (n = 26/197).

Pathway analysis via over-representation analysis (ORA, R *ReactomePA* v1.42.0) revealed enrichment in cytoskeletal organization, ECM interactions, cell motility (**Supplementary Table 3),** critical for melanoma cell migration and invasion, connecting directly to key metastatic regulators *RHOA, CTNNA1*, *COL1A1*, and *SYK* and cancer genes *BRAF*, *TP53* among others (**Supplementary Fig. 3**). We retained all their mutations for downstream analyses and set other "long genes" to null.

**Supplementary Table 3: Over-representation analysis of long genes.** Reactome pathway enrichment of long genes highlights cytoskeletal organization, ECM interactions, and cell motility–related processes.

| ID | Description | q value | Genes |
|---|---|---|---|
| R-HSA-1474244 | Extracellular matrix organization | $4.460\times10^{-9}$ | LAMA5, VWF, VCAN, DST, COL12A1, FBN1, COL7A1, DMD, LAMA1, TNXB |
| R-HSA-3000178 | ECM proteoglycans | $4.588\times10^{-4}$ | LAMA5, VCAN, LAMA1, TNXB |
| R-HSA-3000157 | Laminin interactions | $6.201\times10^{-4}$ | LAMA5, COL7A1, LAMA1 |
| R-HSA-1474228 | Degradation of the extracellular matrix | $2.530\times10^{-3}$ | LAMA5, COL12A1, FBN1, COL7A1 |
| R-HSA-3000171 | Non-integrin membrane-ECM interactions | $2.645\times10^{-3}$ | LAMA5, DMD, LAMA1 |
| R-HSA-2022090 | Assembly of collagen fibrils and other multimeric structures | $2.645\times10^{-3}$ | DST, COL12A1, COL7A1 |
| R-HSA-216083 | Integrin cell surface interactions | $6.039\times10^{-3}$ | VWF, FBN1, COL7A1 |
| R-HSA-1474290 | Collagen formation | $6.246\times10^{-3}$ | DST, COL12A1, COL7A1 |
| R-HSA-8874081 | MET activates PTK2 signaling | $1.271\times10^{-2}$ | LAMA5, LAMA1 |
| R-HSA-8875878 | MET promotes cell motility | $2.129\times10^{-2}$ | LAMA5, LAMA1 |
| R-HSA-8948216 | Collagen chain trimerization | $2.226\times10^{-2}$ | COL12A1, COL7A1 |

#### Supplementary Note 2: Calibrating REVEL thresholds for missense variants

Cancer driver mutations confer a selective advantage by disrupting key cellular functions. We rely on REVEL as an orthogonal strategy to subsequent frequency-based approaches to estimate the impact of missense variants on protein function and exclude variants with minimal functional impact, a crucial step for noise reduction in hypermutated tumors like metastatic melanoma. To define an optimal cancer-specific REVEL threshold for cutaneous metastatic melanoma, we used ClinVar classifications for 1,710 missense variants (937 benign/likely benign (BLB), 773 pathogenic/likely pathogenic (PLP)). Mean REVEL scores were significantly lower for BLB (0.276) than PLP (0.768; Wilcoxon test, $P < 2.2\times10^{-16}$). ROC analysis of our logistic regression model predicting missense PLP variants showed a high AUC (0.918). The maximal Youden index was identified as an optimal REVEL threshold of 0.557 (FPR = 0.133, TPR = 0.845), closely aligning with the genome-wide recommendation of 0.5[8]. However, this threshold excluded 15.8% of PLP variants. For a comprehensive gene perturbation assessment, we set a REVEL threshold of 0.276, allowing 5% PLP false-negative rate based on the cumulative empirical distributions, while filtering out 58.5% of BLB variants (**Supplementary Fig. 4**). Missense variants below this threshold were excluded. We followed the same procedure for pan-cancer analysis ($n_{BLB} = 20{,}893$, $n_{PLP} = 15{,}454$), and used this single REVEL threshold (= 0.386) to maintain a 5% PLP false-negative rate across all 33 TCGA cancer types.

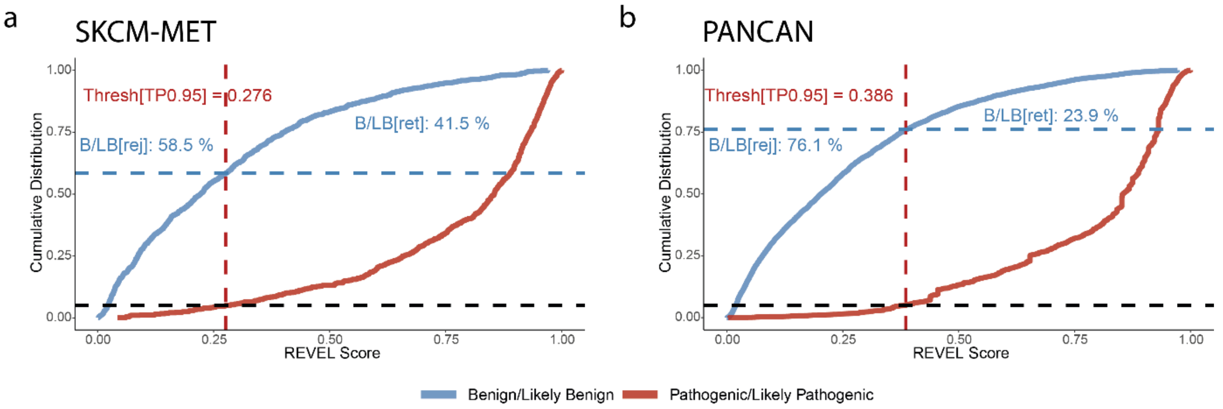


**Supplementary Fig. 4 Calibration of REVEL score thresholds for missense pathogenicity in SKCM-MET and PANCAN. (a)** Cumulative distributions of ClinVar-annotated benign/likely benign (BLB; n = 937) and pathogenic/likely pathogenic (PLP; n = 773) missense variants in SKCM-MET. A REVEL cutoff of 0.275 (allowing 5% false negatives among PLP variants) excludes 58.5% of BLB variants. **(b)** Equivalent analysis in PANCAN (BLB: n = 20,893; PLP: n = 15 454) identifies a 0.386 threshold, which removes 76.1% of BLB variants.

#### Supplementary Note 3: Allelic heterogeneity filter

Striker et al.[9] introduced an approach to identify cancer driver candidates within copy number alteration (CNA) regions by leveraging allelic heterogeneity (AH), where diverse mutations can produce similar functional effects. Allelic heterogeneity is particularly common in tumor suppressor genes, where diverse mutations across multiple sites within the gene can impair its function[10]. We expanded this concept to nominate potentially multiple high-AH genes per locus, since somatic CNAs may contain multiple functional targets[11]. To establish cancer-specific AH cutoffs, we varied locus size and IFF thresholds and measured overlap with the KCG-MEL benchmark, finding significant enrichment across several parameter combinations that grew stronger as thresholds tightened (**Supplementary Fig. 5**). This confirmed our ability to filter out passenger CNAs while enriching for biologically relevant events. Restricting AH filtering to loci containing more than ten genes, we kept the gene with the highest integrated final frequency (IFF) plus any additional genes exceeding 3.5 median absolute deviations above the locus median, a slightly more permissive cutoff ($n_{loci}$ = 10, MAD = 3.5, P = 0.102) than the two least significant results ($n_{loci}$ = 5, MAD = 3.5, P = 0.041; $n_{loci}$ = 10, MAD = 4, P = 0.042). This criterion dramatically shrank our candidate list: GISTIC2 initially flagged loci spanning 1–588 genes (mean = 55), but after AH filtering, locus sizes fell to 1-120 genes (mean = 8), reducing unique CNA-affected genes from 3,746 to 569. Amplification calls per patient dropped from 47.9 to 9.1 on average (P < 0.001), whereas deletions fell from 12.1 to 4.9 (P = 0.153). The same thresholds were then applied in our pan-cancer analyses.

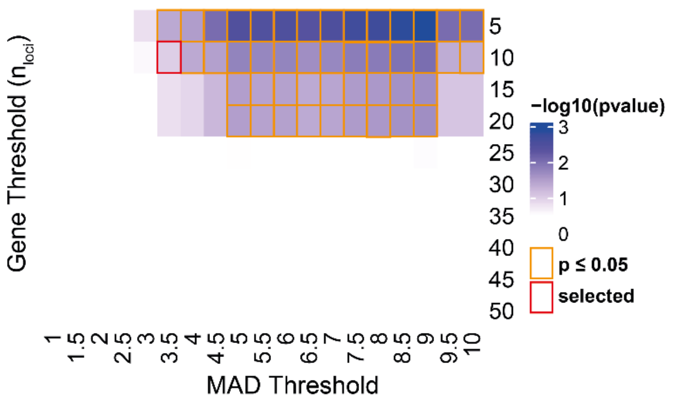


**Supplementary Fig. 5 Optimization of allelic heterogeneity filter parameters for copy-number alterations.** A two-dimensional grid search varied (1) the maximum number of genes per CNA locus (5–50, y-axis) and (2) the MAD threshold (x-axis), scoring each parameter pair by hypergeometric enrichment overlap with the KCG-MEL reference set. Significant enrichment was retained up to a 20-gene locus size or a MAD of 3.5. To maximize stringency just below this significance boundary, we set the locus-size cutoff to 10 genes and the MAD threshold to 3.5. These parameters were also applied uniformly in our pan-cancer comparisons.

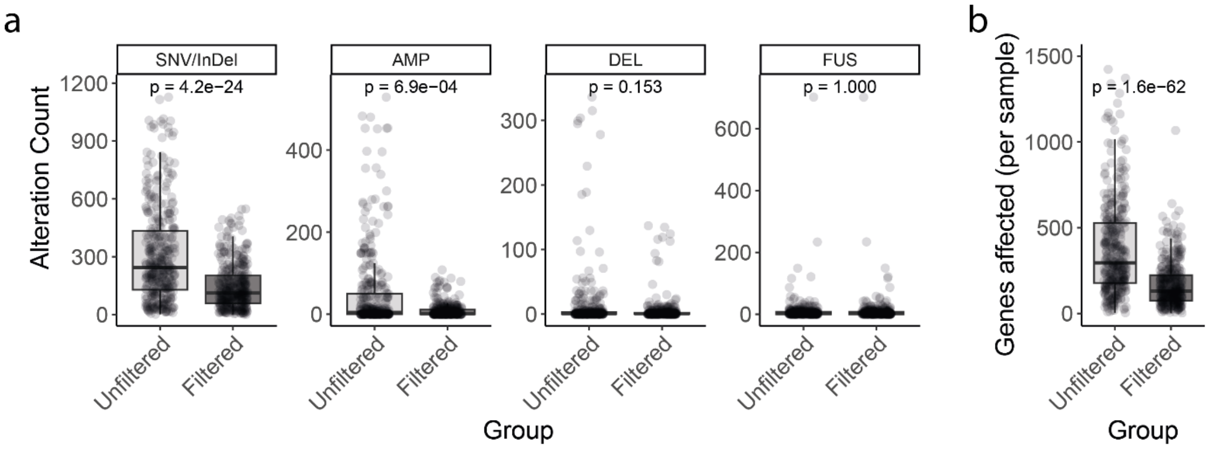


**Supplementary Fig. 6 Impact of Variant Filtering on Gene-Level Alterations across Mutation Matrix Layers. (A)** Boxplots show the per-patient counts of unique genes with at least one variant before (unfiltered) and after (filtered) quality-based filtering in each mutation matrix layer. For SNV/InDels, unfiltered refers to non-silent, protein-coding alterations. Each gene is counted only once per patient, regardless of the number of variants targeting it. For CNAs, unfiltered count includes the number of genes in GISTIC2 peaks before allelic heterogeneity filtering. Paired Wilcoxon signed-rank tests were used to compare unfiltered and filtered counts for each layer; resulting P-values are indicated above the corresponding boxplot pairs. After filtering, the SNV/InDel layer was reduced from 114,469 to 53,577 gene-level mutation events (–53.2%), copy-number amplifications from 17,834 to 3,392 (–81.0%), and deletions from 4,503 to 1,840 (–59.1%), while gene fusions remained unfiltered. P value of paired Wilcoxon rank sum test per layer indicated. **(B)** Number of genes affected (per sample) by at least one layer in the integrated unfiltered versus filtered mutation matrix. P value of paired Wilcoxon rank sum test. Overall, the integrated, binary mutation matrix decreased from 139,684 to 62,098 events (–55.5%) and the total number of affected genes fell from 13,943 to 9,327 (–33.1%).

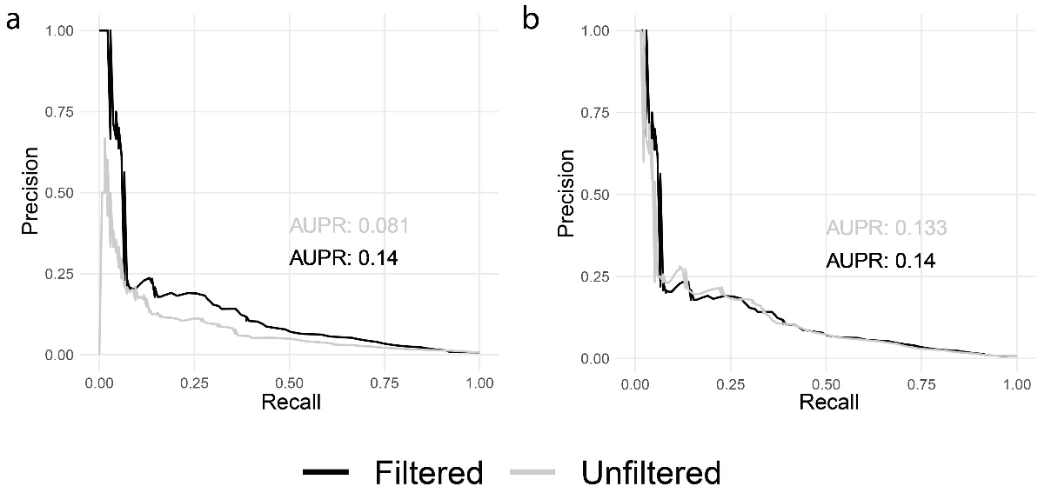


**Supplementary Fig. 7 Comprehensive mutation-matrix filtering and integrative modeling boost driver-gene prediction. (A)** Area under the precision–recall curve (AUPR) for predicting KCG-MEL driver genes using raw (unfiltered, gray) versus quality-filtered (black) mutation matrices based on mutation frequency alone. Quality filtering markedly increases AUPR, demonstrating that removing low-confidence calls substantially improves mutation-frequency–based gene ranking. **(B)** Incremental performance gains from adding copy-number variation (CNV) and gene-fusion (FUS) data to the single-nucleotide variant/indel (SNV/INDEL) layer in the RIPPLET framework. AUPR for SNV/INDEL only (grey) and SNV/INDEL + CNV + FUS (black) shows improvement in cross-cohort predictive accuracy, highlighting the value of comprehensive genomic profiling and integrated analysis. Though SNV/indels dominate numerically, adding CNV and fusion layers further boosts driver-relevant gene prioritization, raising KCG-MEL predictive performance (AUPR 0.133 to 0.140).

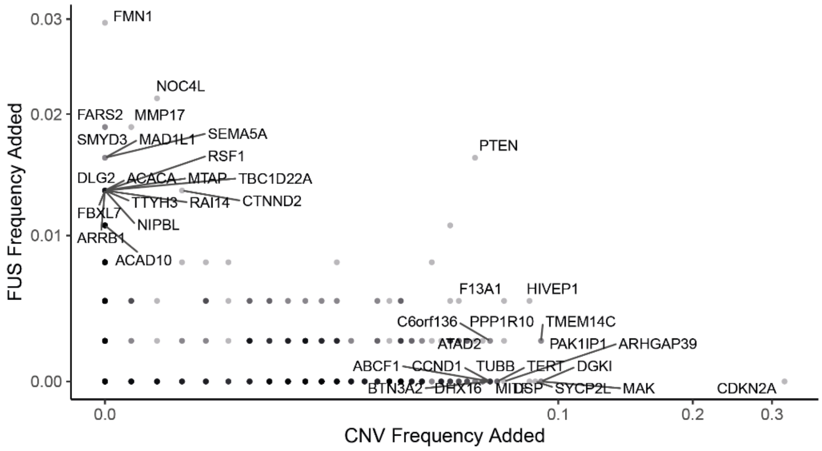


**Supplementary Fig. 8 Layer-specific contribution to mutation frequency in the integrated mutation matrix.** For each target layer (e.g., SNV/INDEL, CNV, FUS), we computed its added frequency by taking the full integrated mutation frequency and subtracting the frequency obtained when all but that layer was omitted (i.e., integrated – [all layers except the target]). The residual counts therefore represent the mutation-frequency attributable solely to the target layer.

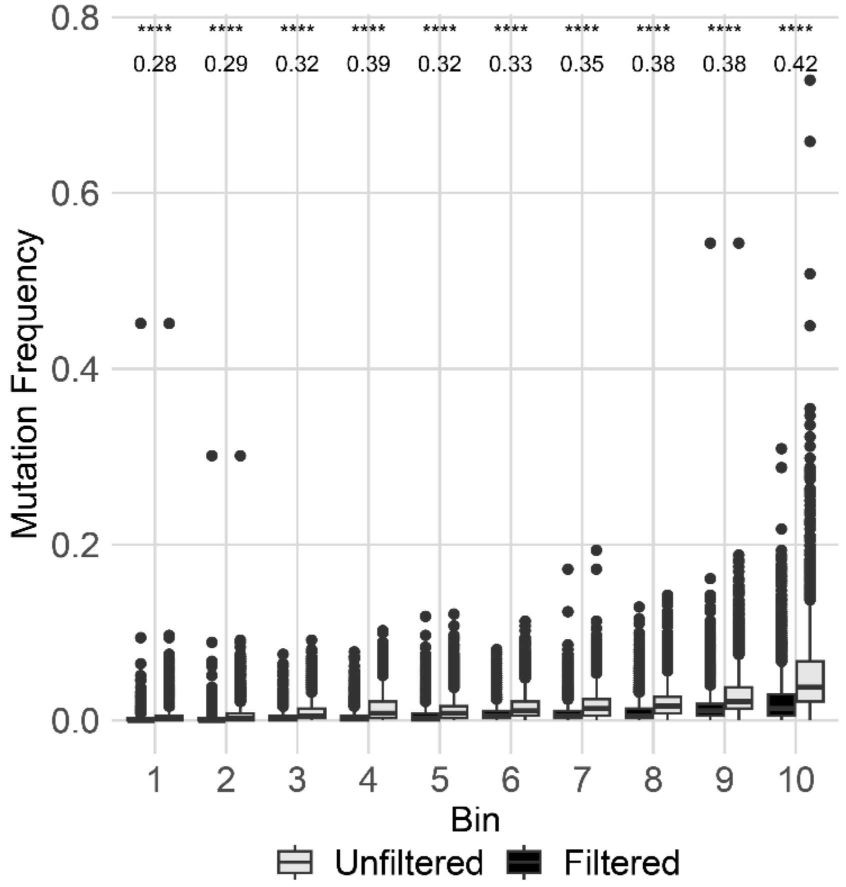


**Supplementary Fig. 9 Reduction in per-gene mutation frequency across gene-length deciles.** Genes were ranked by coding-sequence length and divided into ten equal-sized bins (bin 1 = shortest genes; bin 10 = longest genes). For each bin, boxplots compare per-gene mutation frequencies before (unfiltered) and after (filtered) quality-based filtering. Paired Wilcoxon signed-rank tests were performed within each bin using the *rstatix* R package (v0.7.2); both P-values and Wilcoxon effect sizes (r) are annotated above the boxplot pairs. The most pronounced decrease in mutation frequency after filtering is observed in bin 10, which contains the longest genes.

**Supplementary Table 4 Effect of mutation-matrix filtering on driver identification by cohort-wide mutation frequency.** With the exception of DLBC and THYM, filtering significantly improved performance in 31 of 33 TCGA cohorts. Shown are the area under the precision–recall curve (AUPRC) for unfiltered and filtered mutation frequencies derived from the integrated mutation matrix.

| Cancer Type [TCGA Label] | AUPRC [Unfiltered] | AUPRC [Filtered] |
|---|---|---|
| ACC | 0.189 | 0.245 |
| BLCA | 0.181 | 0.260 |
| BRCA | 0.080 | 0.151 |
| CESC | 0.063 | 0.139 |
| CHOL | 0.108 | 0.112 |
| COAD | 0.129 | 0.203 |
| DLBC | 0.161 | 0.136 |
| ESCA | 0.090 | 0.120 |
| GBM | 0.158 | 0.247 |
| HNSC | 0.139 | 0.193 |
| KICH | 0.188 | 0.236 |
| KIRC | 0.074 | 0.171 |
| KIRP | 0.152 | 0.319 |
| LAML | 0.134 | 0.137 |
| LGG | 0.159 | 0.302 |
| LIHC | 0.067 | 0.215 |
| LUAD | 0.069 | 0.123 |
| LUSC | 0.086 | 0.149 |
| MESO | 0.095 | 0.101 |
| OV | 0.052 | 0.068 |
| PAAD | 0.088 | 0.103 |
| PCPG | 0.288 | 0.312 |
| PRAD | 0.059 | 0.152 |
| READ | 0.091 | 0.142 |
| SARC | 0.075 | 0.096 |
| SKCM | 0.081 | 0.142 |
| STAD | 0.118 | 0.181 |
| TGCT | 0.100 | 0.207 |
| THCA | 0.141 | 0.202 |
| THYM | 0.134 | 0.132 |
| UCEC | 0.273 | 0.307 |
| UCS | 0.142 | 0.151 |
| UVM | 0.176 | 0.182 |

### Cohort-Informed Reweighting

#### Supplementary Note 4: Random walk with restart – Topology bias and network propagation characteristics

An inappropriately chosen graph normalization method can lead to topology bias as shown by Charmpi *et al*[12]. Topology bias is the biased increase or reduction of node scores exclusively due to the network structure and independent the prior information vector $F_0$ (equation 1). Using the procedure of Charmpi et al.[12], we set every node's initial value $F_0 = 1$ and applied network propagation using RWR. With row-wise normalization, as used in RIPPLET, all propagated values remain at 1, confirming no topology-driven bias. By contrast, column-wise normalization artificially inflates scores of high-degree nodes. Although these genes frequently overlap established cancer drivers (mean STRING v12 degree: KCG-MEL = 81.1, KCG-PAN = 81.80 versus 28.09 for non-cancer genes; all P[KCG vs. non-KCG] $< 2.2 \times 10^{-16}$), we intentionally avoid enriching for this signal[12], which could inflate recovery of known drivers and mask novel, low-connectivity candidates. Gene connectivity and positional importance are incorporated later in a pathway-specific context, enabling independent assessment of each gene's positional value within individual cellular processes. We demonstrated topology bias further using *igraph* v1.5.1 in R. We computed degree, betweenness, closeness and PageRank for every node and assigned each score to a decile bin. We then selected 1,000 random seed genes, propagated their scores through 10,000 permutations at restart probabilities $\alpha = \{0.3, 0.5, 0.7, 0.9\}$, and averaged the resulting node scores within each bin. The decile-wise means and variances (across corresponding restart parameters) remained flat across centrality (no shifts observed)[13]. We observe decreasing variances of centralities with decreasing restart probabilities as network scores converge (**Supplementary Fig. 10**).

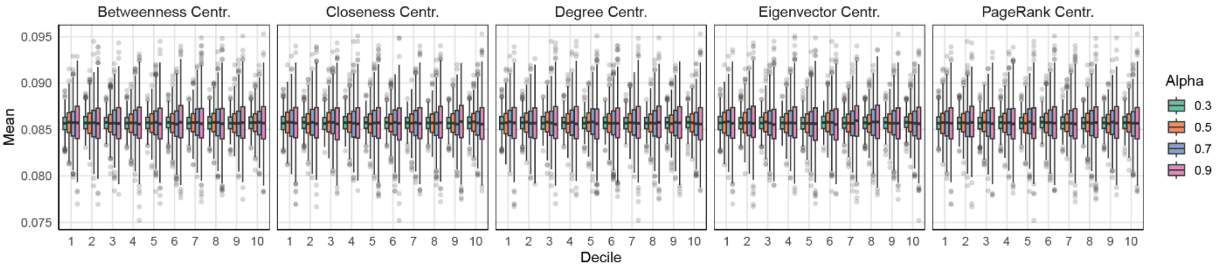


**Supplementary Fig. 10 Row-normalized random walk with restart (RWR) shows no topology bias across centrality metrics.** Permutation tests reveal stable mean centrality values across all metrics and bins, while variance across bins increases with higher restart probabilities ($\alpha$). As $\alpha$ decreases, RWR scores progressively converge toward a uniform distribution.

We fixed restart parameter $\alpha = 0.7$ to balance local versus global diffusion, consistent with prior studies[14–16]. We mapped the cumulative probability mass distribution at various restart parameters (**Supplementary Fig. 11**). For each restart probability α, we iteratively set a single random seed node to 1, run RWR to convergence, and sum the scores of all nodes within n hops from the seed to get the cumulative diffusion mass. We repeat this 1,000 times at each α and average the scores. At α = 0.7, ~75% of the cumulative probability mass is retained on the source node, and ~ 94% within 1 step. With decreasing α, the random walker is more likely to land on nodes further away from the seed node.

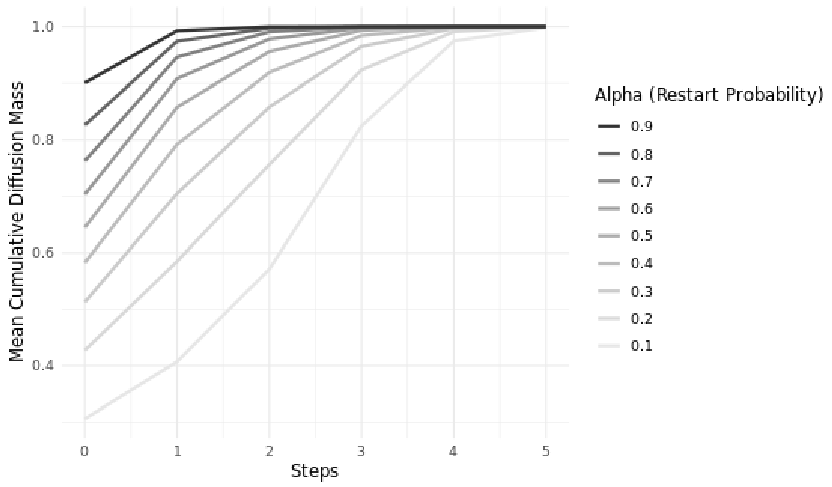


**Supplementary Fig. 11 Cumulative probability mass from the source node across restart probabilities.** For each α, random walk with restart (RWR) was run from a single source node to convergence, and scores were summed for nodes within n steps, yielding the aggregate probability of the walker residing within that step distance.

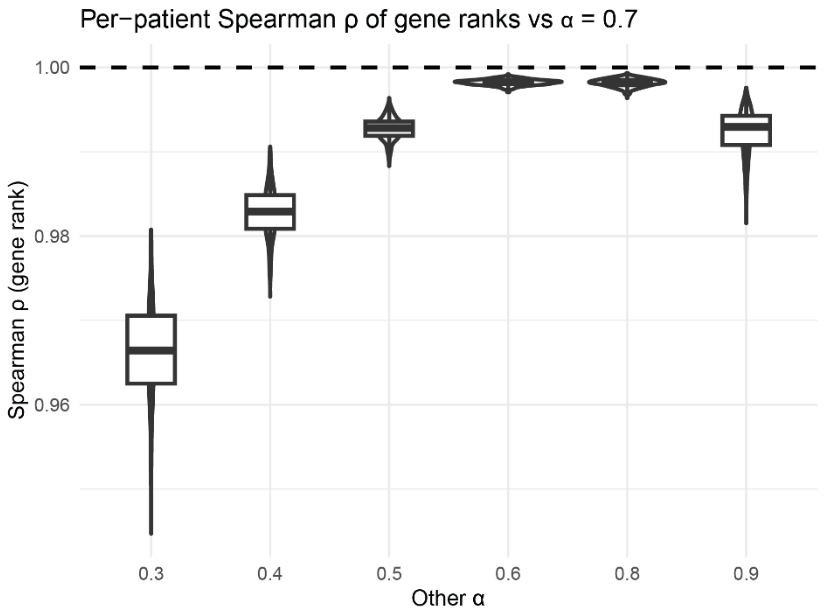


**Supplementary Fig. 12 Random Walk with Restart delivers robust rankings across a range of restart parameters.** We evaluated whether the restart probability parameter α alters patient-specific gene rankings derived from the random walk with restart (RWR). Result are based on the SKCM-MET cohort, using the corresponding contextualized network. For each α, RWR produced a gene×patient score matrix. For every patient and α, genes were ranked in descending score (ties averaged), and Spearman correlations were computed between all α–α′ rank vectors. For each α–α′ pair we visualized per-patient distributions versus reference α (α=0.7) used during final analysis. We observe high correlation coefficients (all mean > 0.966).

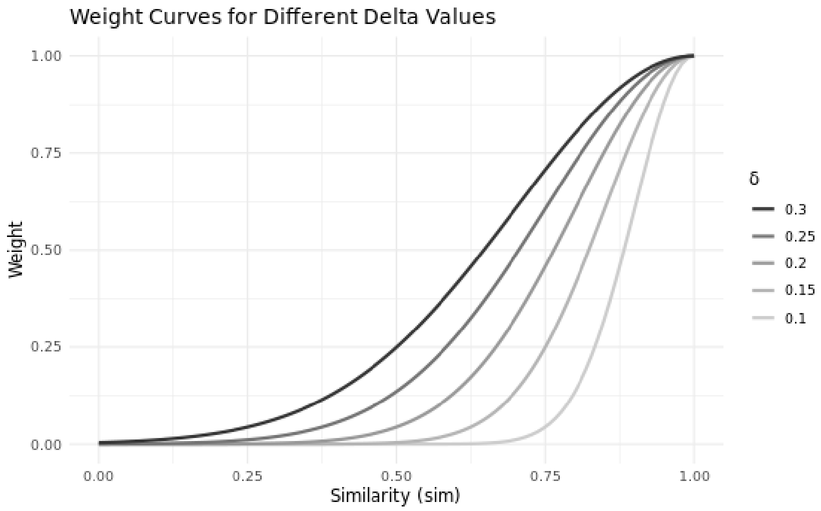


**Supplementary Fig. 13 Decay of angular-similarity weights as a function of the similarity parameter δ.** We use tumor-tumor similarity, formulated as angular similarity in equations 4-5 for the cohort-informed reweighting step. The x-axis shows δ (controlling the steepness of decay from 0 to 1), and the y-axis shows the resulting contribution assigned to an "information-lending" tumor relative to the index tumor. As δ increases, the decay curve flattens, meaning that more dissimilar tumors retain higher relative weight compared to lower δ settings. We default to δ = 0.2. With this δ, weights decay to 0.88 at $s_{ang}$ = 0.9, 0.61 at $s_{ang}$ = 0.8 and fall below 0.05 for $s_{ang} \leq 0.5$ (minimum achievable with our strictly positive vectors; half-maximum weight at sim ≈ 0.765).

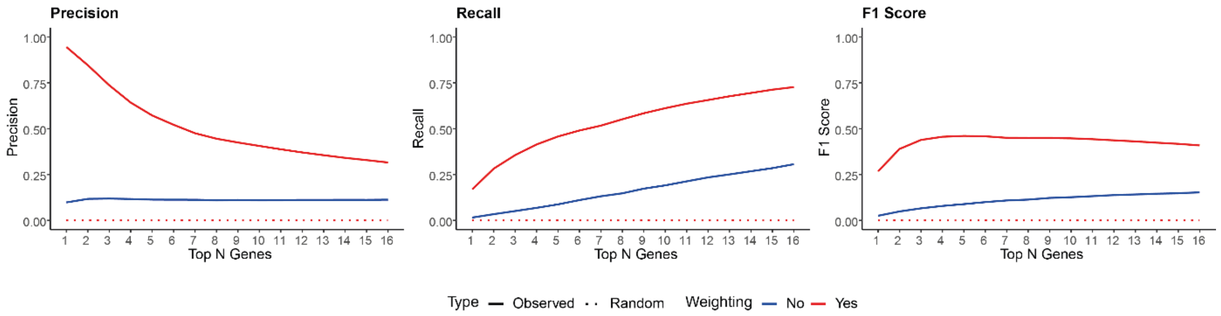


**Supplementary Fig. 14 Cohort-informed gene reweighting permits personalized driver prioritization.** Network propagation on unweighted mutation profiles (no inter-sample information sharing) only modestly exceeds a randomized baseline (gene–mutation links are shuffled, per-sample mutation counts preserved); true positives score higher simply by virtue of being mutated, boosting recall but yielding minimal precision and low F1. Shown are mean precision, recall, and F1 across the top 16 ranks (SKCM-MET threshold, on KCG-MEL). Incorporating cohort-level reweighting dramatically enriches known melanoma drivers among the highest ranks, achieving both high precision and robust F1.

#### Supplementary Note 5: Performance / evaluation metrics

For statistical evaluation, the precision, recall, and their harmonic mean (F-score) were used. These are expressed as:

$$Precision = \frac{TP}{TP + FP}$$

$$Recall = \frac{TP}{TP + FN}$$

$$F - score = 2 * \frac{Precision * Recall}{Precision + Recall}$$

where **TP**, **FP**, and **FN** denote true positives, false positives, and false negatives, respectively. We specifically compute precision, recall, and the F1-score at K. **Precision at K** measures the proportion of the top K retrieved items that are relevant, whereas **Recall at K** quantifies the fraction of all relevant items that appear within these top K positions. The **F1-score** is the harmonic mean of precision and recall, offering a balanced evaluation of both metrics. As such, we consider this the primary measure in performance comparison. Each of these scores ranges from 0 to 1, with higher values indicating better performance. It is important to note that these measures assess only the presence of relevant items within the top K results and do not account for their internal ranking (i.e., they are set-based rather than rank-aware metrics).

To enable fair comparisons across cancer cohorts with different rank thresholds, we define the normalized partial area under the curve (**npAUC**) for precision, recall, and F1. For any metric M∈{precision, recall, F1}, npAUC is calculated as:

$$npAUC_M = \frac{\sum_{i=1}^{K}(M(i) + M(i-1))/2}{K}$$

where K is the cohort's dynamic rank threshold and M(i) is the metric value at rank i. The numerator is the area under the M-vs. rank curve, computed by the trapezoidal rule. Dividing by K (the maximum possible area, since $M \leq 1$ at each rank) scales the result between 0 and 1. By normalizing each partial AUC to its theoretical maximum (K), npAUC produces a single, unitless score, ranging from 0 (worst) to 1 (best), that can be directly compared across cohorts, regardless of their differing rank cutoffs, for personalized driver prioritization.

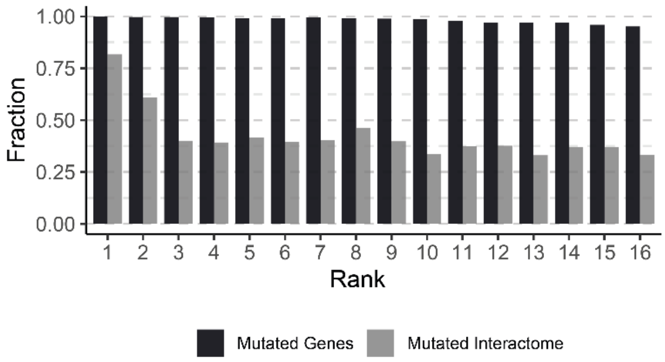


**Supplementary Fig. 15 Fraction of genes mutated (black) or with mutated interactome (grey) in the top 16 ranks across the SKCM-MET cohort.** Genes appearing at the very top of the rankings were predominantly those with mutations present in the input matrix, whereas genes without direct mutations scored highly only through the influence of their mutated network neighbors and thus appeared further down the personalized lists. RIPPLET is designed to prioritize these primary drivers over indirect network effects.

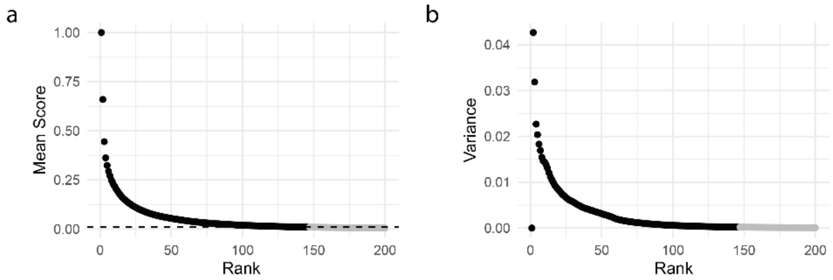


**Supplementary Fig. 16 Mean (A) and Variance (B) of SKCM-MET gene impact profiles across the cohort by gene rank**. The top-ranked driver is normalized to a value of 1, resulting in a variance of zero. In panel A, the dashed horizontal line denotes the default noise threshold (0.01), with genes above this cutoff rendered in black and those below in gray. Genes above threshold extend up to rank ~150, suggesting that beyond a set of high impact perturbations, a number of lower-impact alterations may contribute to the cancer phenotype.

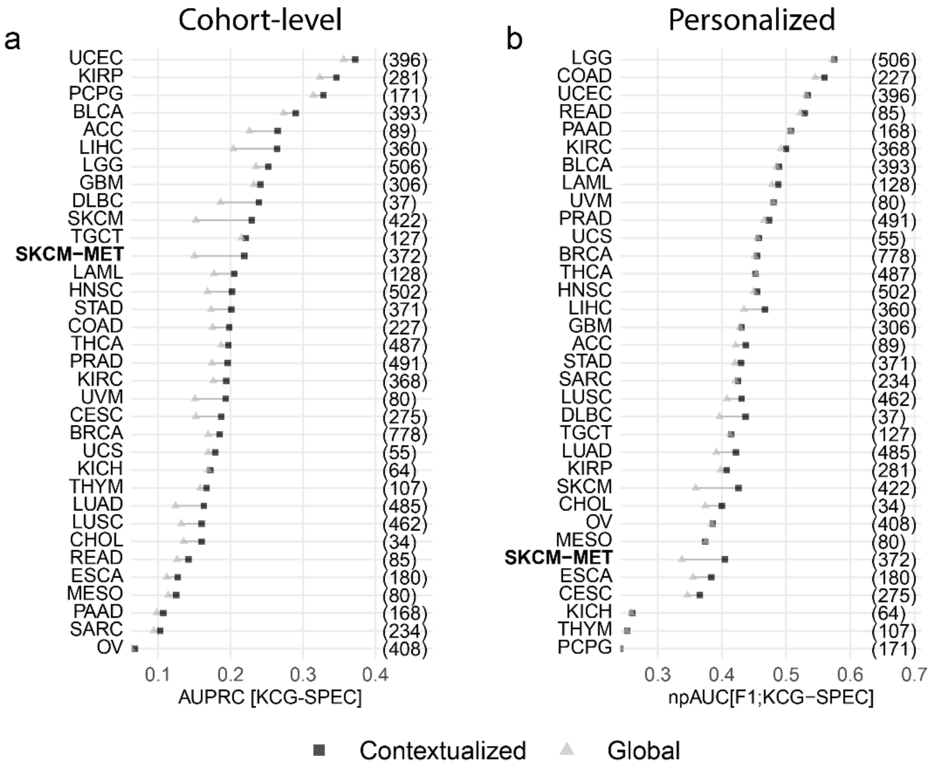


**Supplementary Fig. 17 Contextualized PPI networks result in higher driver prioritization performance. (A)** Cohort-level improvements are based on the average network-propagated score per gene (each propagated profile was column-normalized first to give equal weight to every sample). We ran propagation on both the global and cohort-specific (contextualized) PPI networks, then assessed prioritization accuracy against the KCG-SPEC reference gene sets using area under the precision-recall curve (AUPRC). In the plot, results for the contextualized network are marked by black squares and those for the global network by gray triangles; in all 33 TCGA cohorts the contextualized network outperforms on KCG-SPEC. Mean improvement = 14.969%, range: 1.775% to 50.658%. Contextualized gene importance outperforms global network in 33/33 TCGA and the SKCM-MET cohort. **(B)** Personalized prioritization performance (F1), reported as normalized partial AUC (npAUC) against KCG-SPEC using cohort-specific rank thresholds (see **Supplementary Table 5**), again shows that contextualized networks outperform global in 31 of 33 TCGA cancer cohorts and SKCM-MET. Mean improvement = 3.39%, range: -0.346% to 19.831%.

### Performance & Comparative trial

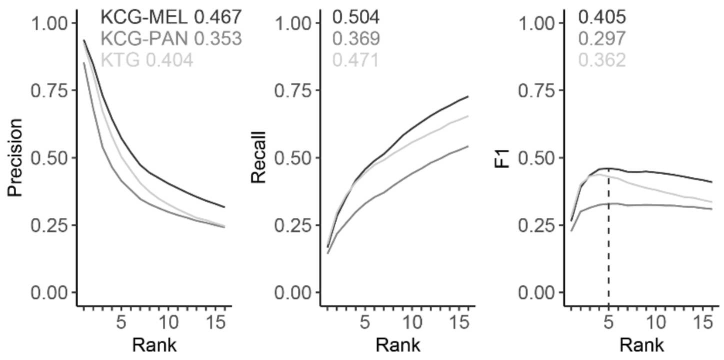


**Supplementary Fig. 18 Performance of RIPPLET across reference driver sets.** RIPPLET demonstrates robust precision, recall, and F1 performance in metastatic melanoma across three gold-standard references (KCG-MEL, KCG-PAN, KTG), evaluated over all 372 patients. Values represent normalized partial area under the curve (npAUC) for precision, recall, and F1. The dashed line in the right panel marks the maximal F1 at rank 5, which served as the cutoff for subsequent gene-level analyses of candidate driver characteristics.

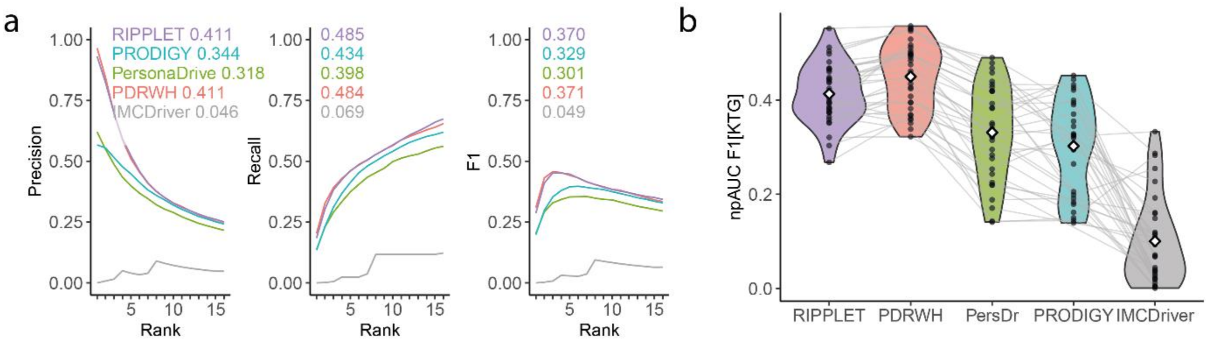


**Supplementary Fig. 19 RIPPLET identifies immune-modulatory genes in SKCM-MET. (a)** Comparative npAUC (precision, recall and F1) on the subset of SKCM-MET samples with matched mutation and expression data (SKCM-MET[sub], n = 320), using immune modulatory genes (KTG) as the benchmark. **(b)** Pan-cancer evaluation across 33 TCGA tumor types, using immune-modulatory genes (KTG), comparing prodigy to 4 state-of-the-art driver prediction tools.

**Supplementary Table 5 Performance comparison of RIPPLET and three state-of-the-art personalized gene prioritization tools.** For each cohort and reference set (KCG-SPEC/KCG-PAN/KTG), we report the cohort name, applied rank thresholds for personalized driver identification, and performance using F1 npAUC up to the assigned rank threshold. SKCM-MET[sub] denotes the SKCM-MET subset with paired expression data (320/322 TCGA samples). PRODIGY performance results are not available for HNSC, MESO, THYM, and UVM due to missing matched normal samples (GTEx).

| Study | Rank Thresh. [SPEC/PAN/KTG] | RIPPLET [F1 SPEC/PAN/KTG] | PersonaDrive [F1 SPEC/PAN/KTG] | PDRWH [F1 SPEC/PAN/KTG] | PRODIGY [F1 SPEC/PAN/KTG] | IMCDriver [F1 SPEC/PAN/KTG] |
|---|---|---|---|---|---|---|
| **SKCM-MET[sub]** | 16/16/16 | 0.410/0.303/0.370 | 0.256/0.296/0.301 | 0.336/0.364/0.371 | 0.285/0.334/0.329 | 0.046/0.049/0.049 |
| ACC | 6/8/8 | 0.442/0.34/0.321 | 0.236/0.231/0.225 | 0.317/0.319/0.322 | 0.178/0.184/0.179 | 0.168/0.105/0.119 |
| BLCA | 12/16/14 | 0.489/0.433/0.481 | 0.353/0.374/0.383 | 0.471/0.475/0.5 | 0.395/0.424/0.444 | 0.082/0.061/0.068 |
| BRCA | 8/8/8 | 0.455/0.428/0.439 | 0.425/0.411/0.419 | 0.501/0.492/0.496 | 0.326/0.321/0.326 | 0.027/0.026/0.028 |
| CESC | 8/10/10 | 0.366/0.358/0.359 | 0.285/0.304/0.295 | 0.376/0.386/0.382 | 0.356/0.376/0.367 | 0.03/0.019/0.02 |
| CHOL | 8/8/8 | 0.399/0.399/0.391 | 0.382/0.358/0.334 | 0.454/0.445/0.427 | 0.439/0.421/0.37 | 0.037/0.031/0.037 |
| COAD | 10/12/10 | 0.559/0.481/0.512 | 0.449/0.467/0.478 | 0.526/0.544/0.547 | 0.284/0.282/0.307 | 0.132/0.11/0.117 |
| DLBC | 14/12/12 | 0.437/0.327/0.403 | 0.243/0.245/0.254 | 0.355/0.37/0.358 | 0.242/0.285/0.328 | 0.109/0.165/0.161 |
| ESCA | 9/12/10 | 0.383/0.347/0.396 | 0.384/0.457/0.458 | 0.453/0.509/0.517 | 0.311/0.342/0.355 | 0.132/0.147/0.149 |
| GBM | 8/8/8 | 0.388/0.359/0.375 | 0.481/0.47/0.49 | 0.526/0.509/0.529 | 0.383/0.388/0.39 | 0.081/0.086/0.101 |
| HNSC | 10/12/10 | 0.456/0.394/0.464 | 0.383/0.403/0.419 | 0.427/0.454/0.494 | NA/NA/NA | 0.01/0.007/0.004 |
| KICH | 20/6/6 | 0.26/0.364/0.351 | 0.065/0.165/0.171 | 0.263/0.382/0.424 | 0.051/0.141/0.139 | 0.157/0.102/0.108 |
| KIRC | 6/8/6 | 0.499/0.461/0.46 | 0.243/0.265/0.283 | 0.468/0.438/0.504 | 0.146/0.142/0.183 | 0.121/0.115/0.114 |
| KIRP | 6/8/6 | 0.406/0.352/0.375 | 0.19/0.231/0.243 | 0.332/0.357/0.367 | 0.126/0.139/0.162 | 0.034/0.035/0.044 |
| LAML | 6/8/7 | 0.48/0.412/0.303 | 0.217/0.202/0.188 | 0.501/0.448/0.361 | 0.211/0.188/0.191 | 0.062/0.033/0.004 |
| LGG | 6/8/8 | 0.575/0.544/0.552 | 0.453/0.423/0.427 | 0.564/0.56/0.557 | 0.226/0.201/0.199 | 0.257/0.246/0.255 |
| LIHC | 8/8/8 | 0.466/0.353/0.383 | 0.321/0.319/0.329 | 0.438/0.426/0.435 | 0.263/0.252/0.265 | 0.021/0.009/0.023 |
| LUAD | 10/12/10 | 0.422/0.337/0.385 | 0.313/0.329/0.339 | 0.382/0.392/0.395 | 0.299/0.31/0.326 | 0.008/0.009/0.012 |
| LUSC | 10/14/12 | 0.43/0.309/0.437 | 0.327/0.353/0.351 | 0.372/0.386/0.445 | 0.38/0.4/0.453 | 0.005/0.016/0.005 |
| MESO | 6/8/6 | 0.374/0.435/0.448 | 0.189/0.287/0.322 | 0.353/0.465/0.475 | NA/NA/NA | 0.042/0.137/0.104 |
| OV | 6/10/8 | 0.39/0.42/0.467 | 0.396/0.399/0.419 | 0.437/0.45/0.497 | 0.405/0.382/0.426 | 0.012/0.026/0.019 |
| PAAD | 8/8/8 | 0.505/0.49/0.504 | 0.432/0.432/0.44 | 0.546/0.54/0.551 | 0.317/0.321/0.331 | 0.355/0.318/0.333 |
| PCPG | 20/8/6 | 0.242/0.333/0.365 | 0.053/0.113/0.143 | 0.226/0.329/0.338 | 0.056/0.12/0.159 | 0.136/0.104/0.093 |
| PRAD | 8/8/6 | 0.473/0.283/0.268 | 0.319/0.303/0.315 | 0.483/0.37/0.369 | 0.176/0.182/0.205 | 0.086/0.031/0.001 |
| READ | 9/10/10 | 0.528/0.46/0.498 | 0.479/0.458/0.468 | 0.559/0.538/0.552 | 0.299/0.28/0.295 | 0.229/0.204/0.227 |
| SARC | 6/8/8 | 0.425/0.328/0.373 | 0.392/0.323/0.348 | 0.51/0.409/0.451 | 0.378/0.305/0.324 | 0.086/0.07/0.069 |
| SKCM | 12/14/12 | 0.428/0.334/0.395 | 0.278/0.316/0.322 | 0.364/0.38/0.394 | 0.28/0.325/0.332 | 0.035/0.04/0.037 |
| STAD | 10/12/10 | 0.429/0.391/0.444 | 0.34/0.378/0.393 | 0.42/0.454/0.462 | 0.262/0.272/0.297 | 0.02/0.02/0.031 |
| TGCT | 6/6/6 | 0.415/0.341/0.353 | 0.285/0.248/0.276 | 0.408/0.387/0.408 | 0.231/0.227/0.231 | 0.219/0.149/0.192 |
| THCA | 6/6/6 | 0.452/0.436/0.425 | 0.143/0.145/0.141 | 0.457/0.461/0.45 | 0.147/0.148/0.15 | 0.293/0.271/0.283 |
| THYM | 20/7/6 | 0.258/0.35/0.38 | 0.098/0.175/0.219 | 0.25/0.341/0.351 | NA/NA/NA | 0.121/0.125/0.16 |
| UCEC | 12/14/14 | 0.476/0.455/0.468 | 0.368/0.374/0.383 | 0.47/0.466/0.49 | 0.293/0.301/0.319 | 0.081/0.066/0.071 |
| UCS | 7/10/10 | 0.457/0.388/0.413 | 0.427/0.421/0.433 | 0.499/0.497/0.511 | 0.39/0.409/0.432 | 0.336/0.234/0.287 |
| UVM | 6/10/6 | 0.481/0.377/0.445 | 0.2/0.146/0.22 | 0.453/0.449/0.484 | NA/NA/NA | 0/0.046/0.04 |

**Supplementary Table 6 Runtime comparison of RIPPLET and three state-of-the-art personalized gene prioritization tools.** For each cohort, we report the cohort name, number of patients ($N_{Pat}$), average number of integrated mutations per sample ($N_{Mut}$), and execution time (d:hh:mm:ss) measured on an 8-CPU, 64 GB RAM system. Runtimes for transcriptomics-based methods include differential expression analysis. SKCM-MET[sub] denotes the SKCM-MET subset with paired expression data (320/322 TCGA samples). PRODIGY runtimes are not available for HNSC, MESO, THYM, and UVM due to missing matched normal samples.

| Study | $N_{Pat}$ | Mean[$N_{Mut}$] | RIPPLET | PersonaDrive | PDRWH | PRODIGY | IMCDriver |
|---|---|---|---|---|---|---|---|
| **SKCM-MET[sub]** | 320 | 157.26 | 0:00:01:21 | 0:00:02:22 | 0:00:23:24 | 0:19:59:51 | 0:00:06:24 |
| ACC | 75 | 31.12 | 0:00:00:27 | 0:00:00:11 | 0:00:00:09 | 0:05:33:17 | 0:00:08:42 |
| BLCA | 387 | 80.94315 | 0:00:02:00 | 0:00:05:58 | 0:00:29:12 | 6:19:34:23 | 0:07:11:10 |
| BRCA | 772 | 35.11917 | 0:00:04:26 | 0:00:12:59 | 0:01:08:56 | 4:09:10:32 | 0:19:40:21 |
| CESC | 271 | 52.85609 | 0:00:01:18 | 0:00:01:00 | 0:00:04:28 | 3:09:22:50 | 0:03:36:28 |
| CHOL | 34 | 20.94118 | 0:00:00:08 | 0:00:00:03 | 0:00:00:02 | 0:05:14:21 | 0:00:01:27 |
| COAD | 214 | 48.74766 | 0:00:00:58 | 0:00:02:57 | 0:00:05:22 | 0:19:36:35 | 0:01:50:03 |
| DLBC | 37 | 75.97297 | 0:00:00:09 | 0:00:00:04 | 0:00:00:07 | 1:04:48:33 | 0:00:04:39 |
| ESCA | 176 | 59.38636 | 0:00:00:48 | 0:00:02:41 | 0:00:03:15 | 0:21:20:06 | 0:01:24:04 |
| GBM | 145 | 32.21379 | 0:00:00:38 | 0:00:00:53 | 0:00:01:18 | 1:06:25:13 | 0:00:32:37 |
| HNSC | 492 | 55.15447 | 0:00:02:20 | 0:00:12:06 | 0:00:44:33 | NA | 0:07:28:48 |
| KICH | 64 | 8.96875 | 0:00:00:15 | 0:00:00:09 | 0:00:00:03 | 0:02:14:28 | 0:00:04:48 |
| KIRC | 365 | 28.10959 | 0:00:01:41 | 0:00:01:06 | 0:00:08:25 | 0:17:45:10 | 0:03:13:14 |
| KIRP | 279 | 27.2043 | 0:00:01:13 | 0:00:00:32 | 0:00:01:18 | 0:14:14:13 | 0:02:00:36 |
| LAML | 113 | 18.11504 | 0:00:00:26 | 0:00:00:14 | 0:00:00:09 | 1:10:33:28 | 0:00:11:12 |
| LGG | 499 | 19.17234 | 0:00:02:25 | 0:00:07:01 | 0:00:27:49 | 2:08:33:57 | 0:02:27:39 |
| LIHC | 353 | 41.74788 | 0:00:01:33 | 0:00:02:30 | 0:00:06:57 | 1:22:17:54 | 0:01:19:36 |
| LUAD | 479 | 91.79749 | 0:00:02:31 | 0:00:08:42 | 0:00:45:29 | 3:18:42:40 | 0:12:31:49 |
| LUSC | 457 | 97.18381 | 0:00:02:25 | 0:00:15:29 | 0:00:56:13 | 9:07:17:34 | 0:12:57:17 |
| MESO | 80 | 19.0125 | 0:00:00:21 | 0:00:00:11 | 0:00:00:11 | NA | 0:00:10:26 |
| OV | 284 | 53.79577 | 0:00:01:20 | 0:00:06:59 | 0:00:11:33 | 4:20:53:03 | 0:03:22:49 |
| PAAD | 161 | 21.91304 | 0:00:00:44 | 0:00:01:24 | 0:00:01:11 | 0:20:42:24 | 0:00:41:13 |
| PCPG | 169 | 8.08284 | 0:00:00:47 | 0:00:00:16 | 0:00:00:07 | 0:12:25:29 | 0:00:40:34 |
| PRAD | 488 | 21.94467 | 0:00:02:36 | 0:00:02:13 | 0:00:13:20 | 1:12:13:51 | 0:06:43:54 |
| READ | 84 | 41.42857 | 0:00:00:27 | 0:00:00:41 | 0:00:00:31 | 0:08:41:03 | 0:00:12:28 |
| SARC | 231 | 39.60173 | 0:00:01:11 | 0:00:01:39 | 0:00:02:22 | 4:10:39:32 | 0:01:31:01 |
| SKCM | 418 | 113.2273 | 0:00:02:14 | 0:00:03:01 | 0:00:37:22 | 6:01:41:15 | 0:09:57:00 |
| STAD | 345 | 65.26957 | 0:00:01:41 | 0:00:04:13 | 0:00:13:10 | 1:18:25:55 | 0:06:05:11 |
| TGCT | 126 | 12.25397 | 0:00:00:37 | 0:00:00:17 | 0:00:00:11 | 0:12:16:17 | 0:00:20:24 |
| THCA | 484 | 5.419421 | 0:00:02:25 | 0:00:01:29 | 0:00:05:28 | 0:17:23:00 | 0:02:32:12 |
| THYM | 102 | 7.392157 | 0:00:00:26 | 0:00:00:09 | 0:00:00:03 | NA | 0:00:10:17 |
| UCEC | 154 | 119.1623 | 0:00:00:45 | 0:00:01:29 | 0:00:04:01 | 2:13:29:46 | 0:02:05:19 |
| UCS | 55 | 53.21818 | 0:00:00:16 | 0:00:00:25 | 0:00:00:13 | 0:17:20:21 | 0:00:09:04 |
| UVM | 79 | 33.55696 | 0:00:00:20 | 0:00:00:12 | 0:00:00:11 | NA | 0:00:04:51 |

### Topological Pathway Scoring

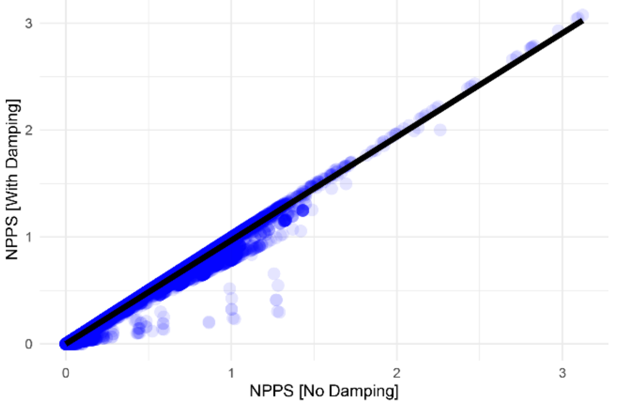


**Supplementary Fig. 20: Damping rescues singular pathways without changing SPIA scores.** We extended the original SPIA workflow[17], which requires a non-singular (I-B) during processing, where B is the normalized adjacency matrix. The original implementation discards pathways with $|\det(B-I)| \leq 10^{-7}$. Following Tarca's suggestion that simple transformations can remove singularities, we rescue only the failing pathways by dampening B → $(1-\epsilon)B$ (here: $\epsilon = 0.01$), which shrinks eigenvalues and renders (I-B) invertible in most cases. Pathways that still fail $|\det(B-I)| > 10^{-7}$ after dampening are excluded. This enabled inclusion of 228 additional pathways. To verify no bias, we recomputed normalized pathway perturbation scores (NPPS) in SKCM-MET for pathways already non-singular and compared undamped (x-axis) versus damped (y-axis) runs: scores were highly concordant for most pathways (Pearson r = 0.998).

### Pathway Convergence

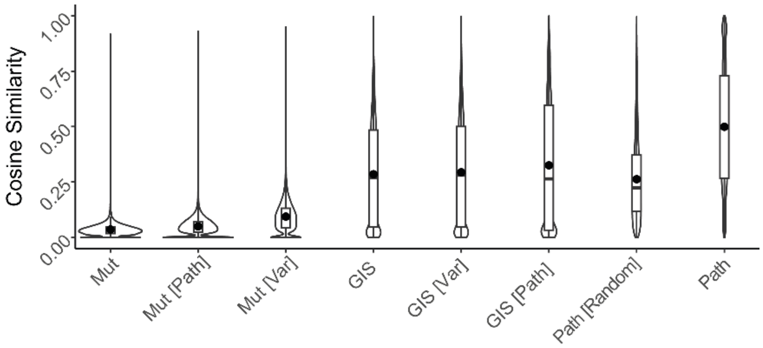


**Supplementary Fig. 21 Average pairwise cosine similarities between tumors across three data layers:** Binary mutation profiles (Mut), RIPPLET gene impact scores (GIS), and RIPPLET pathway perturbation scores (Path). Gene-level comparisons were performed on (1) all 12,057 SKCM-MET genes scored in the PPI network, (2) the 5,109 genes contained in our 579 curated pathways, and (3) the 579 most variable genes. To ensure that the biological meaning of pathways (i.e. the specific genes in each pathway) are driving the signal, we randomized the pathway memberships by shuffling the gene identities but keeping the structure and number of genes in the pathway the same. The resulting average similarities were: Mut = 0.04, Mut[Path] = 0.05, Mut[Var] = 0.09, GIS = 0.28, GIS[Var] = 0.29, GIS[Path] = 0.33, randomized pathways = 0.26, and true pathways = 0.50.

### Drug Sensitivity Prediction

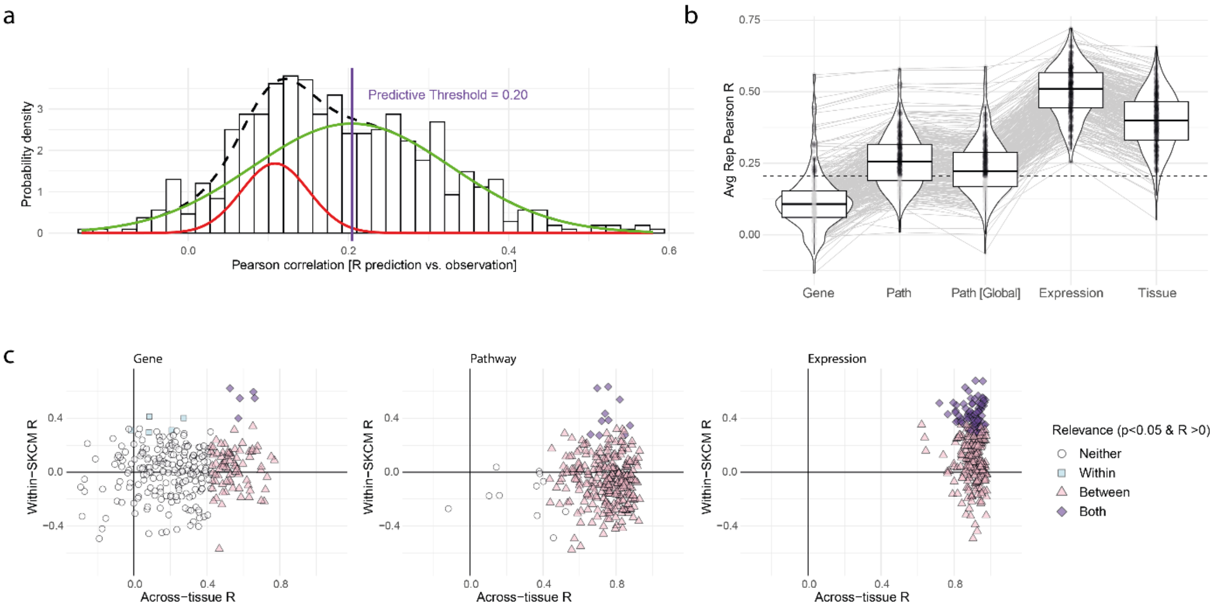


**Supplementary Fig. 22 Tissue-agnostic pan-cancer Elastic Net model performance.** Elastic Net (EN) models were trained on 630 cancer cell lines spanning 23 tissue types. **(A)** Distribution of Pearson correlation coefficients (R) between predicted and observed drug responses for mutation-only models (gene and pathway). A two-component Gaussian mixture model (*mixtools* v2.0.0.1 in R) was fitted to this distribution; the higher-mean component defines "informative" models, and the predictive threshold ($\theta$) is set at the intersection of the two Gaussian densities = 0.205[18]. **(B)** Average 5-fold cross-validated Pearson R for five feature sets: gene mutations (Gene), RIPPLET pathway scores (Path), RIPPLET pathway scores with a global gene impact model (Path [Global]), RNA expression (Expression), and one-hot encoded tissue-of-origin (Tissue). Models with $R > \theta$ (predictive) are shown in black; those with $R \leq \theta$ (non-predictive) in gray. **(C)** Between- versus within-tissue performance, illustrated for gene, pathway and expression models. *Between-tissue* performance was assessed by averaging observed and predicted responses across cell lines within each tissue and evaluating the model's ability to recover drug response differences across all 23 tissues. *Within-tissue* performance was calculated as the Pearson R between predicted and observed $\log(IC_{50})$ values for melanoma cell lines (SKCM) using the pan-cancer model[19]. Circles indicate no significant positive correlation; squares indicate significance only within-tissue; triangles indicate significance only between-tissue; and diamonds indicate significance at both levels ($P < 0.05$ and $R > 0$). Mean Pearson R across models: Gene[Between] = 0.28, Gene[Within] = -0.05; Pathway[Between] = 0.73, Pathway[Within] = 0.019; Expression[Between] = 0.903, Expression[Within] = 0.173. Mutation-based pan-cancer models exhibit low cross-tissue accuracy, which is substantially improved by pathway features and maximized by expression data, known to be an excellent tissue-of-origin (TOO) predictor[20]. Nonetheless, pathway-level features capture the same signals, matching its correlation index to TOO performance. By contrast, gene-level mutation data correlate poorly, likely because cancer heterogeneity obscures tissue information. Still, within-tissue performance remains modest and only a subset of drugs exceeded the predictive threshold, thus motivating the development of tissue-guided elastic net (TG-EN) models.

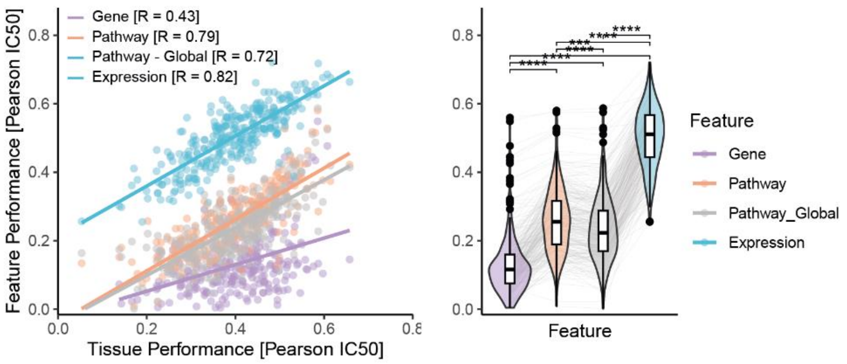


**Supplementary Fig. 23 Tissue contextualization minimally affects the correlation between pathway- and tissue-level predictive performance.** To assess whether contextualization, i.e. removing non-expressed nodes using tissue RNA-seq, drives the strong correlation between pathway and tissue-of-origin predictive performance, we recalculated gene impact scores using the global (non-contextualized) network. Predictive accuracy declined modestly (0.79 to 0.72; $p_{adj} = 8.5\times10^{-4}$, Benjamini-Hochberg), indicating that most of the predictive power reflects pathway models capturing truly tissue-specific perturbations.

**Supplementary Table 7 Summary of cell-line cohorts for TCGA-guided transfer learning.** For each TCGA tumor type (as mapped by GDSC), the number of matched cancer cell lines is listed, along with an indicator of whether that cohort was included in the downstream tissue-guided elastic net (TG-EN) analysis (see next section, only types with ≥ 15 lines).

| Cancer Type [TCGA Label] | #CCL | TG-EN |
|---|---|---|
| ACC | 1 | No |
| BLCA | 18 | Yes |
| BRCA | 51 | Yes |
| CESC | 14 | No |
| COAD | 45 | Yes |
| DLBC | 34 | Yes |
| ESCA | 35 | Yes |
| GBM | 34 | Yes |
| HNSC | 39 | Yes |
| KIRC | 32 | Yes |
| LAML | 26 | Yes |
| LGG | 17 | Yes |
| LIHC | 15 | Yes |
| LUAD | 62 | Yes |
| LUSC | 14 | No |
| MESO | 21 | Yes |
| OV | 34 | Yes |
| PAAD | 29 | Yes |
| PRAD | 6 | No |
| SKCM | 54 | Yes |
| STAD | 24 | Yes |
| THCA | 16 | Yes |
| UCEC | 9 | No |

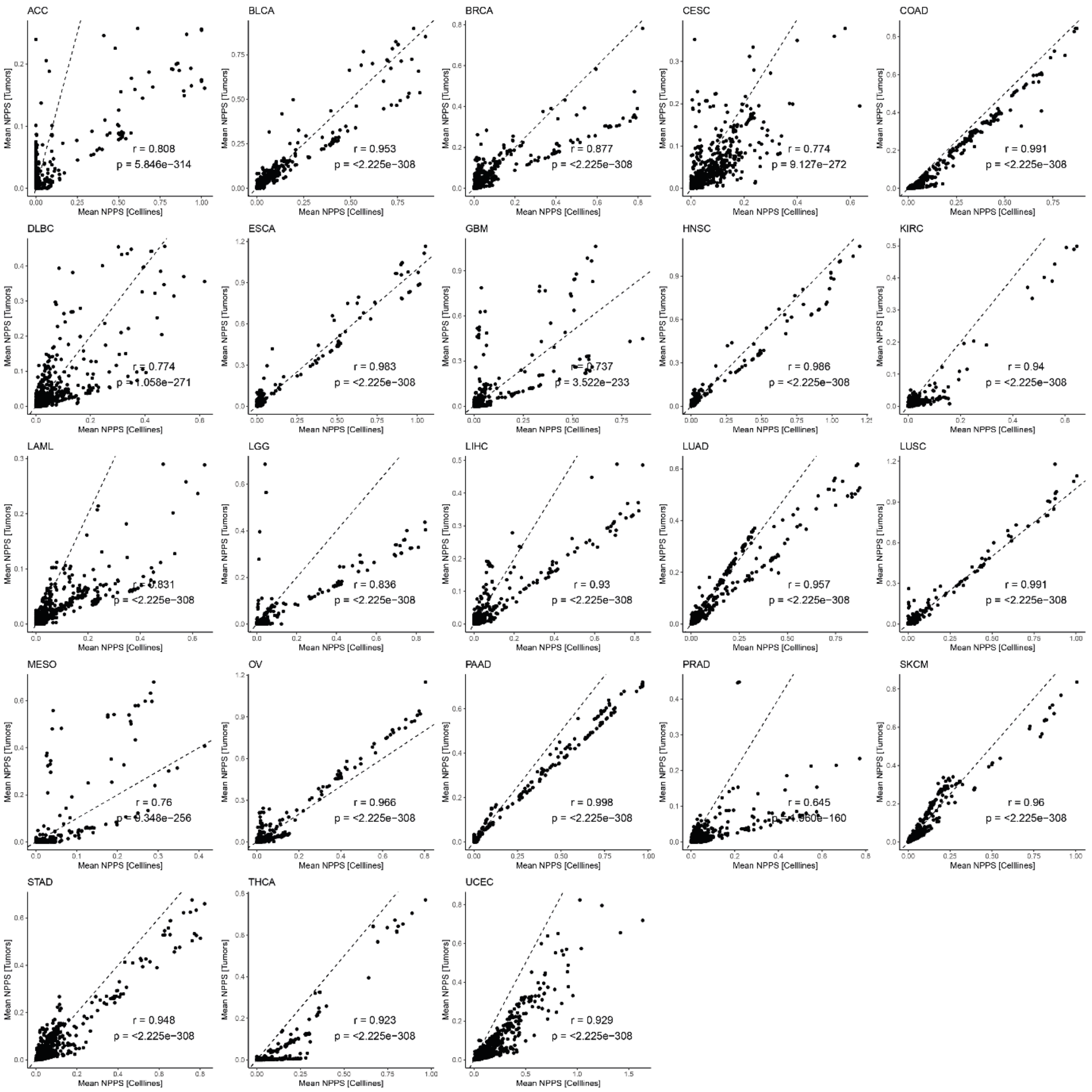


**Supplementary Fig. 24 Transfer learning for gene impact calculation between cancer cell lines (CCL) and TCGA tumor samples yields highly correlated pathway perturbation profiles.** Each point represents a mean RIPPLET-normalized pathway perturbation score (NPPS), with cell line NPPS on the x axis and matched TCGA tumor NPPS on the y axis. CCL gene impact scores (GIS) derive from transfer learning via RIPPLET: network-propagated CCL mutation matrices are compared against all tumor profiles in the corresponding TCGA cohort (GDSC assignments), enabling cohort-informed reweighting of CCL GIS. Reweighted scores are collapsed into pathway features and normalized to produce NPPS values. Because R uses IEEE 754 double precision, the smallest nonzero P value that can be reported is $2.225\times10^{-308}$.

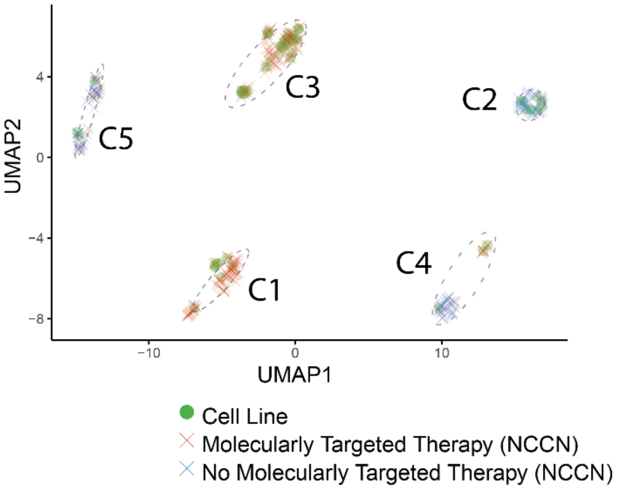


**Supplementary Fig. 25 UMAP embedding of 372 SKCM-MET tumors (**crosses; blue if mutation in BRAF, MAP2K1/2, KIT = National Comprehensive Cancer Network (NCCN)-indicated drug targets[21]; else red**) and 50 GDSC melanoma cell lines (green circles) in the shared RIPPLET pathway space.** We embedded both tumor-derived and cell-line pathway profiles in two dimensions using R *umap* (v0.2.10.0). Five clusters (C1-C5) contain mixed tumors and cell lines; clusters C1/C3 are enriched for samples with NCCN indicated mutations, while C2/C4/C5, particularly enriched for NRAS mutants, lack standard targeted options.

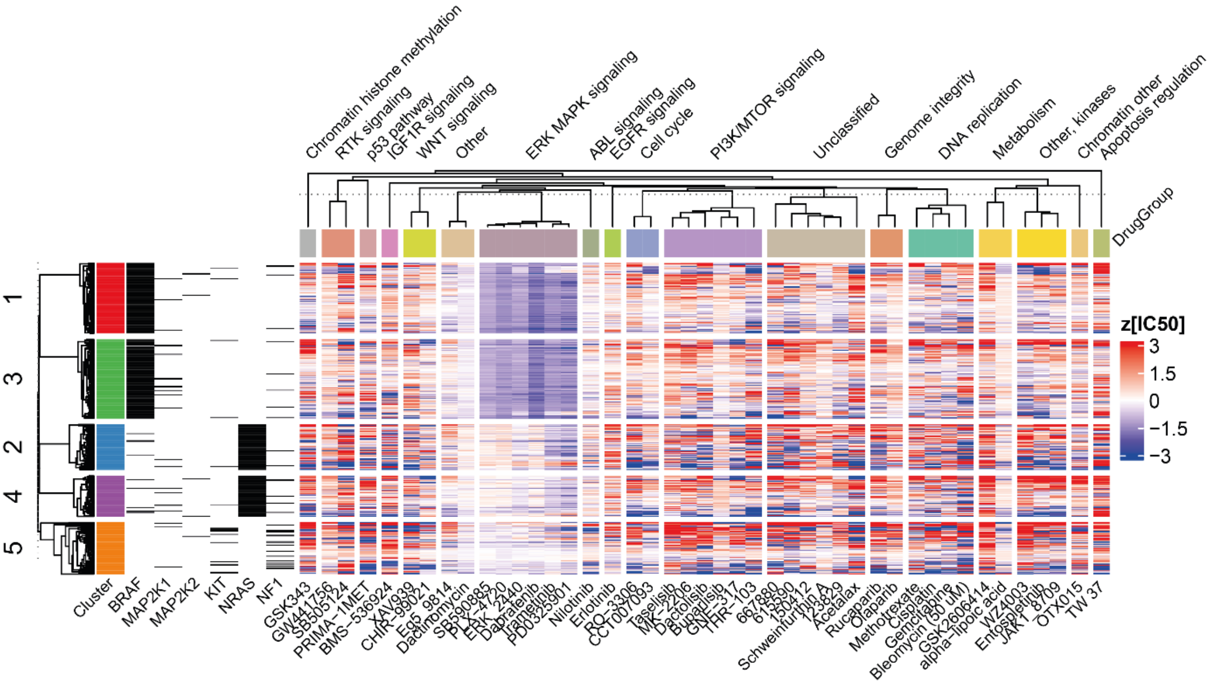


**Supplementary Fig. 26 Mutation Profiles and Predicted Drug Sensitivities Across Patient Clusters.** Left: Cluster annotations from UMAP (**Supplementary Fig. 25**) with binary mutation status for BRAF, MEK (MAP2K1, MAP2K2), KIT, NRAS, and NF1. Right: per-patient drug sensitivity predictions (z-scored $IC_{50}$ relative to the cancer cell line $IC_{50}$ distribution), grouped by drug pathway target. Blue denotes increased predicted sensitivity. Shown are predictive drug models (n = 133) with sensitivities in ≥5% of the cohort (n = 44).

**Supplementary Table 8 Tissue-guided elastic net (TG-EN) performance across cancer types.** Number of predictive TG-EN models (R > θ = 0.205) and average within-tissue Pearson correlation.

| Study | $N_{pred}$[Gene] / Mean R | $N_{pred}$[Path] / Mean R | $N_{pred}$[Expression] / Mean R |
|---|---|---|---|
| BLCA | 231 / 0.376 | 188 / 0.292 | 251 / 0.488 |
| BRCA | 168 / 0.242 | 118 / 0.198 | 251 / 0.365 |
| COAD | 215 / 0.293 | 209 / 0.278 | 278 / 0.506 |
| DLBC | 148 / 0.203 | 241 / 0.343 | 230 / 0.360 |
| ESCA | 202 / 0.287 | 210 / 0.299 | 263 / 0.442 |
| GBM | 148 / 0.234 | 168 / 0.246 | 272 / 0.478 |
| HNSC | 137 / 0.208 | 212 / 0.298 | 205 / 0.306 |
| KIRC | 184 / 0.256 | 208 / 0.308 | 250 / 0.394 |
| LAML | 158 / 0.240 | 213 / 0.316 | 205 / 0.315 |
| LGG | 51 / 0.149 | 112 / 0.376 | 126 / 0.654 |
| LIHC | 146 / 0.266 | 200 / 0.392 | NA / NA |
| LUAD | 81 / 0.158 | 114 / 0.181 | 259 / 0.365 |
| MESO | 162 / 0.248 | 212 / 0.310 | NA / NA |
| OV | 192 / 0.295 | 132 / 0.214 | 261 / 0.462 |
| PAAD | 176 / 0.248 | 141 / 0.221 | 269 / 0.504 |
| SKCM | 171 / 0.236 | 133 / 0.204 | 235 / 0.329 |
| STAD | 144 / 0.221 | 199 / 0.306 | 269 / 0.540 |
| THCA | 84 / 0.337 | 90 / 0.310 | NA / NA |

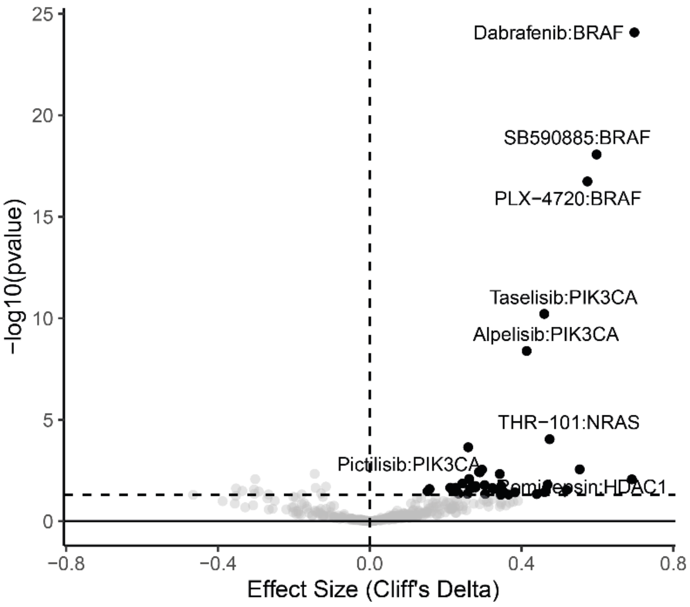


**Supplementary Fig. 27:** To assess whether mutations in a drug's nominal target confer altered sensitivity, we examined 630 GDSC cancer cell lines across 401 drug–target pairs from 222 drug compounds with assigned gene target. We retained only drug-target pairs with at least five mutant and five wild-type lines, resulting in a final set of 191 compounds (321 drug-target pairs). For these remaining pairs, cell lines were grouped by target-gene mutation status and compared using a Wilcoxon rank-sum test. Associations were considered mutation-dependent only if mutants showed significantly greater sensitivity (Cliff's δ > 0 and P < 0.05). 38/191 = 19.9% of compounds showed significantly greater drug sensitivity in mutant cell lines.

### Survival analysis

**Supplementary Table 9: Prognostic pathway signature in metastatic melanoma.** Listed are Reactome[22] pathway names, LASSO coefficients used to compute the pathway risk score (PRS), and univariate P values for each SKCM-MET sub-cohort.

| Pathway (Reactome) | LASSO Coefficient | Univariate P [TCGA] | Univariate P [TUPRO] |
|---|---|---|---|
| Toll Like Receptor 4 (TLR4) Cascade | -3.3781262 | 0.032 | 0.1208 |
| Retrograde neurotrophin signalling | -2.6357577 | 0.061 | 0.390 |
| Regulation of FZD by ubiquitination | -3.0089894 | 0.138 | 0.818 |
| WNT5A-dependent internalization of FZD4 | -3.7236710 | 0.098 | 0.462 |
| RET signaling | -2.2126623 | 0.145 | 0.120 |
| Transcriptional Regulation by MECP2 | -0.5785792 | 0.061 | 0.080 |

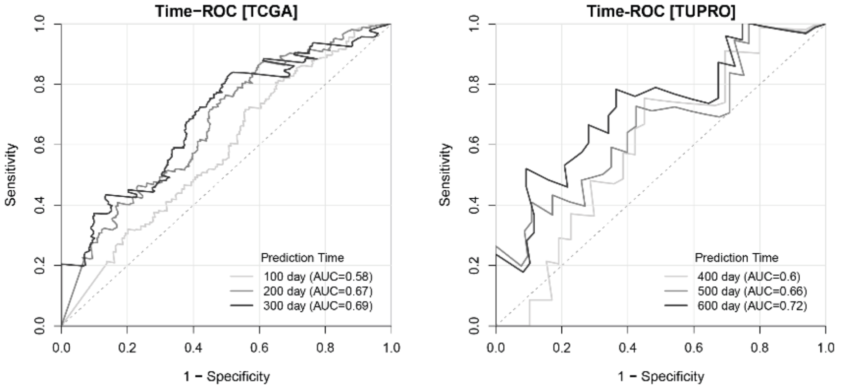


**Supplementary Fig. 28 Time-ROC of TCGA (100, 200, 300 days) and TUPRO (400, 500, 600 days).** Set points were chosen to match each cohort's follow-up. Median OS: TCGA = 102 days; TUPRO = 680 days.

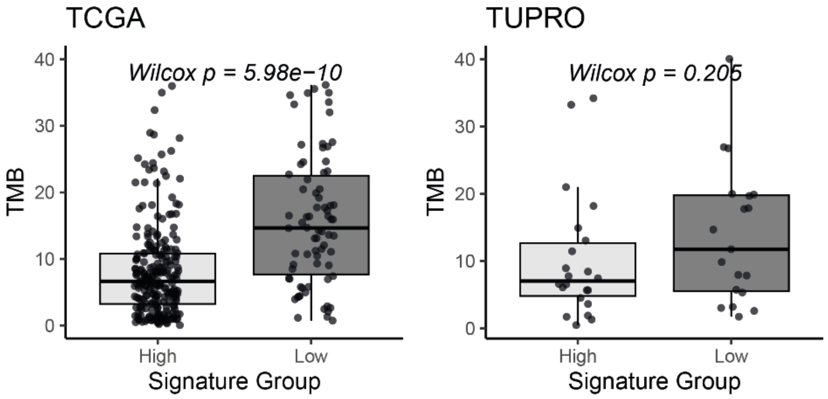


**Supplementary Fig. 29 Tumor Mutational Burden Stratified by Pathway Risk Score in SKCM-MET Sub-Cohorts.** Indicated is the P value of Wilcoxon rank sum test. TMB is calculated as non-silent mutations/Mb of genome. We see significantly higher TMB in TCGA in patients with lower risk scores ($P = 5.98\times10^{-10}$). The same trend is observed in TUPRO, although not significant (P = 0.205)

### Immune analysis

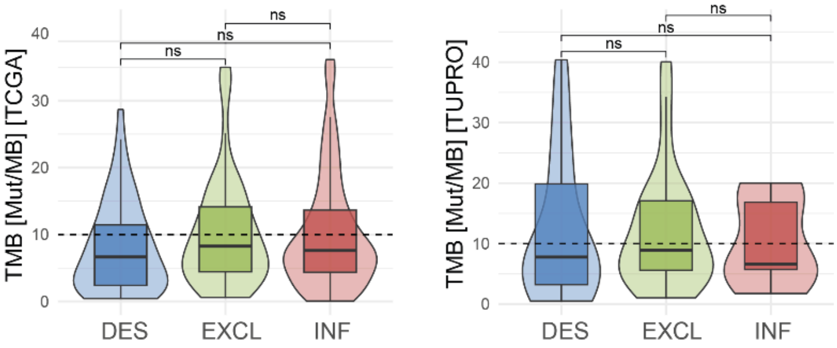


**Supplementary Fig. 30** Tumor mutational burden by immune phenotype in the SKCM-MET[TCGA] and –[TUPRO] cohorts. Significance by Wilcoxon rank-sum test (ns, not significant, threshold: P ≤ 0.05).

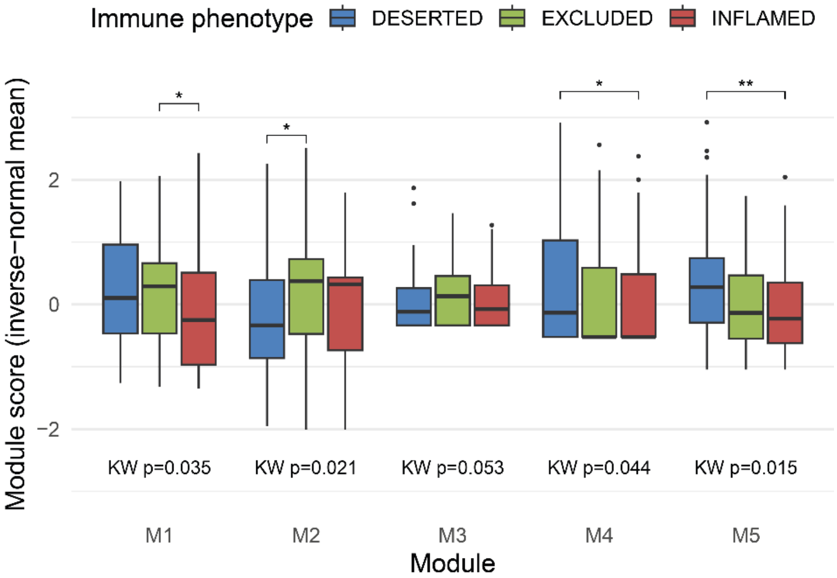


**Supplementary Fig. 31 Immune-modulating pathway modules.** Module scores were calculated for five unsupervised clusters (kmeans) of pathways significantly associated with immune phenotype. Shown are Kruskal–Wallis P-values and significant pairwise Wilcoxon tests (* *P*<0.05, ** *P*<0.001). Modules: M1 oxidative stress/p53 regulation, M2 MAPK signaling, M3 TGFβ/Notch developmental programs, M4 neuro-GPCR/GABAergic signaling, and M5 RHO GTPase–mediated cytoskeletal dynamics. M2 was less perturbed in deserted tumors versus excluded (P = 0.021), whereas M5 scored higher (P = 0.00086), implicating cytoskeletal remodeling alongside MAPK signaling as potential barriers to immune-cell infiltration.

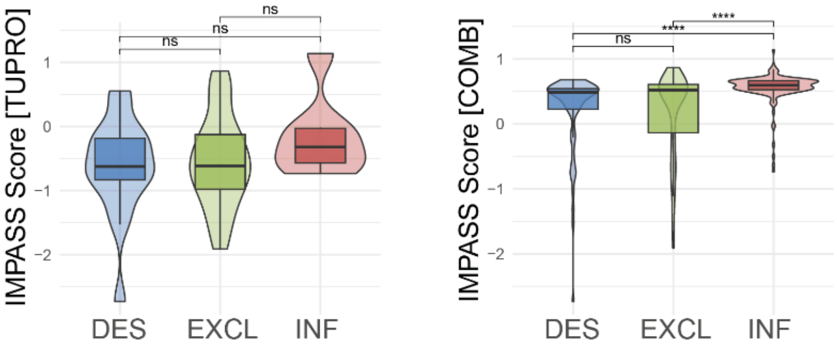


**Supplementary Fig. 32 IMPASS Signature in TUPRO and TUPRO + TCGA [COMB] across immune phenotypes.** Significance by Wilcoxon rank-sum test: ns, not significant; ****, P ≤ 0.0001.

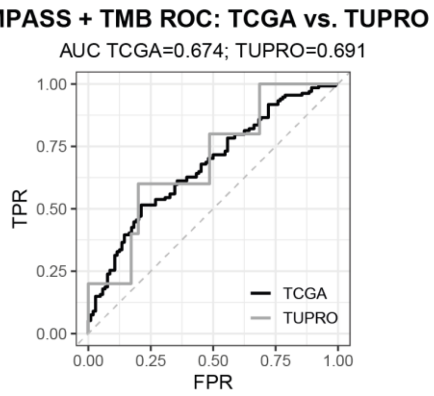


**Supplementary Fig. 33 Combined performance IMPASS + TMB on Inflamed vs. Non-Inflamed tumors of TCGA and TUPRO.** ROC curves for a logistic regression meta-model integrating the IMPASS signature score and tumor mutational burden (TMB) in the SKCM-MET[TCGA] and SKCM-MET[TUPRO] cohorts; corresponding area under the ROC curve (AUROC) values are indicated.

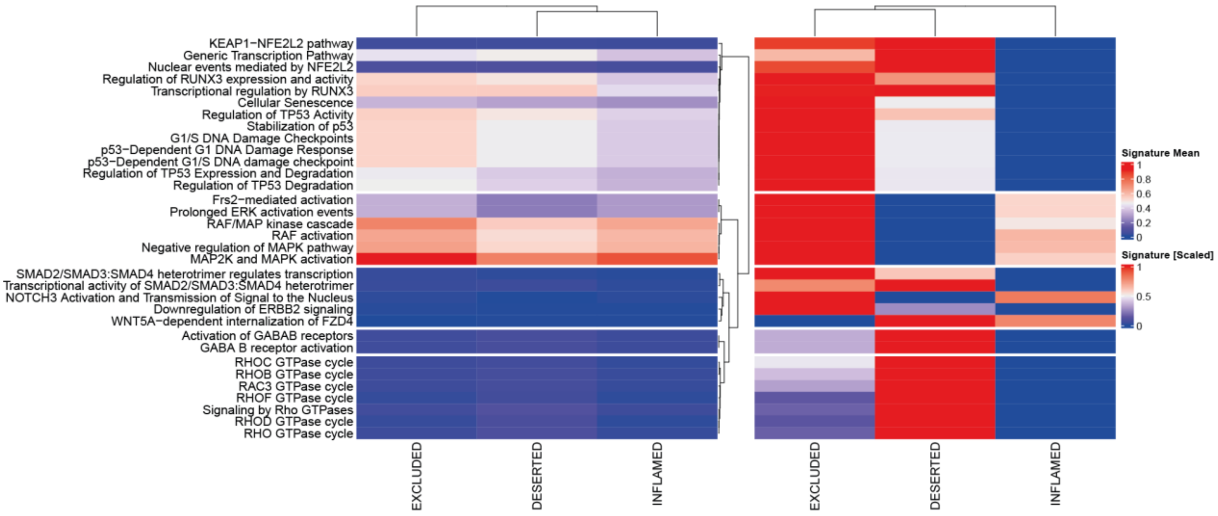


**Supplementary Fig. 34 IMPASS Signature Contributions**. **Left:** Unscaled mean RIPPLET pathway perturbation score across immune phenotypes. **Right:** Scaled (0 to 1) mean RIPPLET pathway perturbation score across immune phenotypes.

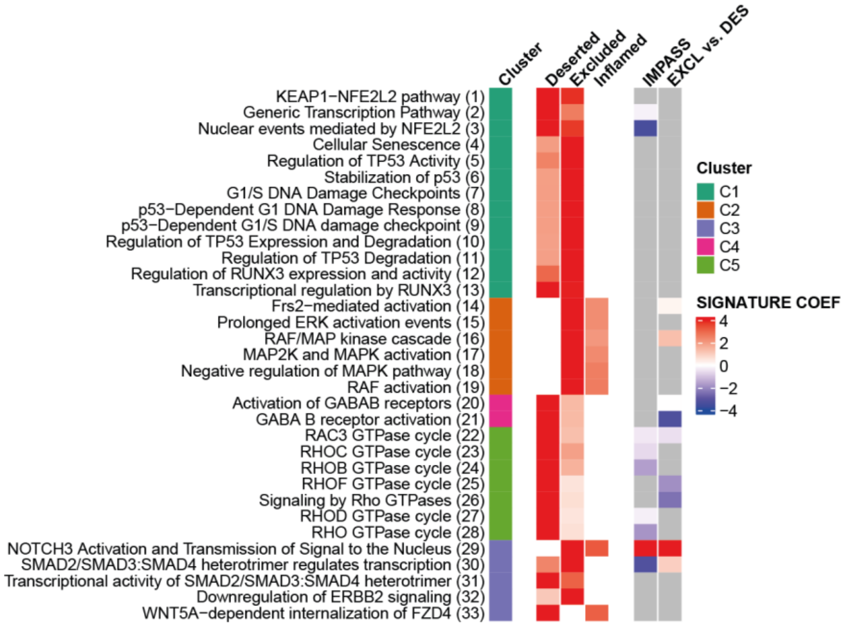


**Supplementary Fig. 35 Full Signature contributions for IMPASS (Classification: Inflamed vs. Non-Inflamed) and Classifier 2 (Excluded vs. Deserted on Non-Inflamed tumors). Left:** pathway names with cluster annotations; **Center:** min–max–scaled pathway perturbation scores across immune phenotypes; **Right:** heatmap of LASSO-derived coefficients at the optimal λ (selected by five-fold cross-validation on the SKCM-MET[TCGA] cohort), where positive weights drive classification toward Inflamed (or Excluded) and negative weights toward Non-Inflamed (or Deserted).

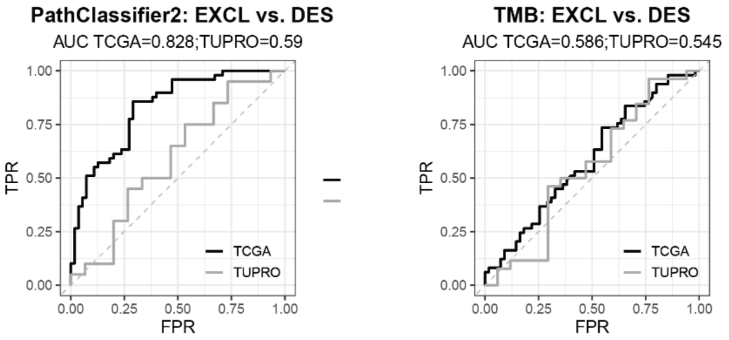


**Supplementary Fig. 36 Classification performance Classifier 2 vs TMB on non-inflamed (excluded vs. Deserted) tumors.** Both models were retrained specifically on this classification task; corresponding AUROC values are indicated.
